# Analysis of genome sequences of marine isolates of *Saccharomyces cerevisiae*

**DOI:** 10.64898/2026.10.26.761992

**Authors:** Muhammad Farhan Aslam, Abdelrahman Zaky, Xirong Hou, Yihan Chen, Yufei Hu, Christopher E. French

## Abstract

*Saccharomyces cerevisiae* is the workhorse of the fermentation industry, as well as a dominant laboratory model organism and chassis for synthetic biology. The great majority of known isolates are from terrestrial sources, but a number of strains are reported to have been isolated from seawater and marine sediments and organisms. Marine strains may have superior tolerance to salt and fermentation inhibitors, and other properties which may be useful in industry, especially in processes based on seawater and marine biomass, as well as casting new light on the physiology and evolution of this species. We have recently reported the isolation of several marine strains of *S. cerevisiae* and comparisons with terrestrial strains for ethanol production. Here we describe the genome sequences of three marine *S. cerevisiae* strains, AZ65, AZ118 and AZ123. To the best of our knowledge, these are the first genome sequences of marine *S. cerevisiae* strains for which published isolation data are available. Superior salt and inhibitor tolerance of the strains, compared to multiple terrestrial and industrial comparators, was confirmed. Analysis of the *ENA* locus, believed to be related to salt tolerance in *S. cerevisiae*, did not show strong evidence for modification of copy number, protein sequence, or promoter sequence of this locus in response to a marine habitat. The physiological and genetic basis of salt and inhibitor tolerance thus remains to be determined.

## 1. Introduction

*Saccharomyces cerevisiae* is one of humanity’s oldest domesticated species, having been used in the manufacture of alcoholic beverages for at least 9000 years (Bai et al., 2022). *S. cerevisiae* is arguably the most important organism in the modern fermentation industry, being responsible for the vast majority of potable ethanol and bioethanol production, as well as being engineered for large-scale production of other valuable commercial products such as artemisinin and farnesene (Benjamin et al., 2016). It is also a dominant model eukaryote for laboratory studies, being the first eukaryote to have its genome sequenced (Goffeau et al., 1996), and is also a dominant organism for synthetic biology experiments, including the first synthetic eukaryotic genome, Sc2.0 (Archer et al., 2026).

A number of recent studies have examined the evolutionary and ecological history of *S. cerevisiae* through large-scale genome sequencing and comparison. The dominant natural habitat of this species may be the bark of oak and related trees (Bai et al., 2022). However, isolates have been reported from a wide variety of habitats, including marine habitats. Kutty and Philip (2008) reviewed reports of marine yeast isolation, and concluded that yeasts in general are widespread in marine environments; however, *S. cerevisiae* specifically is mentioned only in the context of one study, where it was found to be common in the intestines of farmed rainbow trout, though the genus *Saccharomyces* was also reported from other marine environments. It appears that other genera of yeast are much more abundant in marine environments. Tian et al. (2021) reported attempts to isolate *S. cerevisiae* from the South China Sea; ten strains of *S. cerevisiae* were isolated from marine animals and plants but none from seawater or sediments. Baba et al. (2022) reported isolation of *S. cerevisiae* strains from tidal flats in the Ariake Sea near Japan. Other earlier reports of marine isolates of *S. cerevisiae* are described by Zaky et al (2014). The 1011 yeast strains investigated by Peter et al (2018) include 5 annotated as being from seawater (codes BQR, BEV, BET, AQI and AFE, or to give them their full names, DBVPG4381 from Italy, EXF-5247 and EXF-5248 from Slovenia, and CBS8291 and CBS8292 from Sweden) but none of these strains appear to be associated with published information.

The presence of *S. cerevisiae* in marine systems is interesting, since marine organisms are often noted to possess superior resistance to environmental stressors (Greetham et al., 2019). The dominant position of *S. cerevisiae* in the fermentation industry makes this characteristic of particular interest, since the presence of fermentation inhibitors, such as acetate and furfural, in media derived from lignocellulosic materials, is a major barrier to the expansion of bioethanol production from non-food materials. Furthermore, large-scale expansion of the fermentation industry to increase production of renewable chemicals from non-fossil carbon sources is likely to lead to problems with freshwater supply; processes based on seawater and marine resources may alleviate these problems (Zaky et al., 2021a). Thus, marine strains of *S. cerevisiae* may be valuable in the future fermentation industry.

Zaky et al. (2016) reported an improved method for isolation of marine yeasts, describing isolation of 116 strains from UK, US and Egyptian seawater samples. A number of these strains were identified as *S. cerevisiae*, including strains later designated as AZ65, AZ118 and AZ123. These three strains were originally isolated and identified by Dr Abdelrahman Zaky during his doctoral research, as previously reported in Zaky et al. (2016). Greetham et al. (2019) reported that these strains showed elevated tolerance to important fermentation inhibitors, and Zaky et al. (2018, 2020) reported that strain AZ65 showed superior ethanol production in a seawater-based system compared to an industrial comparator strain, NCYC2592. Life Cycle Analysis (Zaky et al, 2021b) showed that such a process would have considerable environmental advantages compared to a standard ethanol production process. It is therefore of interest to consider the physiological and genetic basis for salt and inhibitor tolerance in *S. cerevisiae*, and whether marine strains could be adapter for industrial use, or whether these characteristics could be transferred to industrial strains by genetic manipulation.

Sodium and osmotic-stress tolerance in *S. cerevisiae* have been well studied. The major locus involved in sodium tolerance is the *ENA* locus on chromosome 4, which typically includes 1 to 6 tandem repeats of the *ENA* gene (**E**xtrusion of **Na**+) located between the divergently transcribed *RSM10* and *KRS1* genes (Ruiz and Arino, 2007). Doniger et al. (2008) noted that *KRS1* and the *ENA* genes are among the most variable genes in *S. cerevisiae*, and Peter et al. (2018) noted that copy number variation at the *ENA* locus was one of the few genetic characteristics which showed a clear association with any physiological trait (in this case, Na+ resistance). The reference *S. cerevisiae* genome, from strain S288C, has three copies of *ENA*, designated *ENA1, ENA2* and *ENA5* (the numbering appears to refer to an earlier sequence of this locus from another strain which possessed 5 copies). There is some suggestion in the literature that *ENA1* may be more closely associated with Na+ tolerance and *ENA2* with Li+ tolerance (Ruiz and Arino, 2007). The promoter region of *ENA1* is well studied, and integrates three major stress-sensing pathways: the calcineurin–Crz1 pathway via CDRE motifs, the HOG MAPK pathway via CRE/Sko1 motifs, and the general stress response via Msn2/Msn4-binding STRE motifs (Ruiz and Arino, 2007). This architecture enables rapid, multi-input induction under saline, osmotic, or alkaline stress. However, there appears to be little or no published information about expression of *ENA2* or *ENA5*, or regulation of their promoter regions.

Here we present the genome sequences of marine *S. cerevisiae* strains AZ65, AZ118 and AZ123. To the best of our knowledge, these are the first genome sequences of marine *S. cerevisiae* strains for which published isolation information is available. We examine the *ENA* locus and find no clear evidence for a role in improved salt tolerance in these strains; thus, the physiological and genetic basis of salt and inhibitor tolerance requires further investigation.

## 2. Methods and Materials

### 2.1 Strains and culture conditions

*S. cerevisiae* was cultured on YPD medium (1% yeast extract, 2% peptone, 2% glucose) and stored as glycerol stocks at −80 °C. Marine strains AZ65, AZ118 and AZ123 were isolated from UK seawater as previously described (Zaky et al., 2016). The laboratory strain BY4741 (S288C-derived) and three industrial reference strains (NCYC2592, NCYC1187, and NCYC2780, obtained from NCYC) served as comparators for phenotypic assays; the three NCYC strains were selected to represent differing salt tolerances reported among *S. cerevisiae* isolates (Peter et al., 2018).

### 2.2 Phenotypic characterisation under salt and inhibitor stress

Salt tolerance was assessed in YPD broth supplemented with NaCl at 0.6 M (baseline seawater salinity), 1.0 M (moderate osmotic stress), or 1.5 M (severe osmotic stress), or with a multi-ion sea salt mixture (Sigma-Aldrich S9883; 40 g/L) simulating natural seawater ionic composition. Media were filter-sterilised through 0.22 µm membranes. Tolerance to lignocellulose-derived inhibitors was assessed in YPD supplemented with 5-hydroxymethylfurfural (HMF; 15 mM), furfural (10 mM), or acetic acid (60 mM, adjusted to pH 5.5). In all assays, cultures were inoculated at OD600 = 0.1 and incubated at 30 °C with orbital shaking at 200 rpm for 72 h, with YPD as the control. Final OD600 was recorded at the endpoint. Each condition was tested in biological triplicate, and strain effects were evaluated by one-way ANOVA with Tukey HSD post hoc comparisons (significance at p < 0.05) in SPSS v27.0.

### 2.3 Genome assembly and quality assessment

Cell pellets of AZ65, AZ118 and AZ123 were submitted to Plasmidsaurus for genomic DNA extraction, library preparation, and sequencing on both Oxford Nanopore and Illumina platforms. Hybrid assemblies integrating Nanopore long reads and Illumina short reads were generated for each strain. Assembly quality was assessed with QUAST using the *S. cerevisiae* S288C genome (∼12.1 Mb) as reference. Completeness was evaluated with BUSCO v5.4.4 using the saccharomycetes_odb10 dataset (2,137 single-copy orthologs).

### 2.4 ENA locus identification and copy number validation

Reference ENA coding sequences from BY4741 (ENA1, ENA2, ENA5) were used as queries in BLASTN and TBLASTN searches (BLAST+ v2.13.0; ≥85% identity, ≥80% query coverage). A tandem array was defined as ≥2 ENA genes within <10 kb on the same contig. Copy numbers were validated by mapping Illumina reads with Minimap2 (v2.24) and calculating per-base coverage with Bedtools (v2.30.0); coverage ratios at ENA loci were compared against genome-wide single-copy gene averages.

### 2.5 ENA sequence alignment and phylogenetic analysis

All 13 full-length ENA coding sequences (3,276 bp each) were aligned using CLUSTALW v1.2.4 (default parameters). Pairwise sequence identities and nucleotide composition statistics were calculated. Phylogenetic reconstruction used the UPGMA method in MEGA X. For strain-level comparison, ITS1/5.8S/ITS2 regions from reference *S. cerevisiae* strains were compared with those from the hybrid assemblies. Because the three marine isolates share essentially identical ITS sequences, the low within-species ITS variability precludes intraspecific resolution by maximum likelihood inference.

## 3. Results

### 3.1 Marine strains show enhanced salt and inhibitor tolerance compared to non-marine strains

Initially we sought to reassess the previously reported growth under salt and inhibitor stress of marine strains AZ65, AZ118 and AZ123 under standardised conditions which would allow easy comparison with published data from other sources (for example, Peter et al., 2018). In addition to laboratory strain BY4741 (a derivative of S288C), we used the comparator industrial strain NCYC2592, previously used by Zaky et al. (2021b) in fermentation studies, as well as two further strains, NCYC1187 and NCYC2780, which were chosen as they both appear in the dataset of Peter et al. (2018), one with relatively low salt tolerance and one with relatively high salt tolerance.

To establish the phenotypic basis for the genomic comparisons that follow, the three marine-derived isolates and a panel of reference strains were assessed for growth under salt stress. Strains were cultured in YPD medium supplemented with NaCl at 0.6 M (baseline seawater salinity), 1.0 M (moderate osmotic stress), and 1.5 M (severe osmotic stress), and with a multi-ion sea salt mixture (40 g/L) simulating natural seawater ionic composition. Growth was recorded as OD600 after 72 h at 30 °C. Across all conditions, the marine isolates consistently outperformed the laboratory and industrial reference strains, and AZ123 was the most salt-tolerant strain tested (Figure 1). At baseline salinity (0.6 M NaCl), AZ123 reached the highest growth of any strain (OD600 = 16.0), exceeding NCYC2780 (14.5), AZ65 (14.0), AZ118 and NCYC1187 (both 13.5), NCYC2592 (13.0), and the laboratory reference BY4741 (10.5). Under moderate osmotic stress (1.0 M NaCl), AZ123 again ranked highest (13.0), ahead of AZ65 (12.5), AZ118 and NCYC2780 (both 11.0), and NCYC1187 and NCYC2592 (both 9.5), while BY4741 declined to 8.0. In the multi-ion sea salt mixture, AZ65 (12.0) and AZ123 (11.5) ranked highest, followed by AZ118 (10.5), NCYC1187 (9.5), NCYC2592 (9.0), and BY4741 (8.5); NCYC2780 was the lowest at 7.0, notably below its NaCl-only performance and suggesting sensitivity to the additional divalent cations in the sea salt mixture. Under severe stress (1.5 M NaCl)—a condition at which most *S. cerevisiae* strains show severe inhibition—the marine isolates retained measurable growth (AZ118 = 4.5; AZ123 = 4.0; AZ65 = 3.5), as did NCYC2780 (4.0) and NCYC1187 (3.5), whereas NCYC2592 andBY4741 dropped to 2.5. Averaged across all four conditions, AZ123 ranked first (mean OD600 = 11.13), followed by AZ65 (10.50), AZ118 (9.88), NCYC2780 (9.13), NCYC1187 (9.00), and NCYC2592 (8.50), with BY4741 consistently the least tolerant (7.38). One-way ANOVA confirmed significant strain effects at every salt condition (p < 0.001).

**Figure 1:**
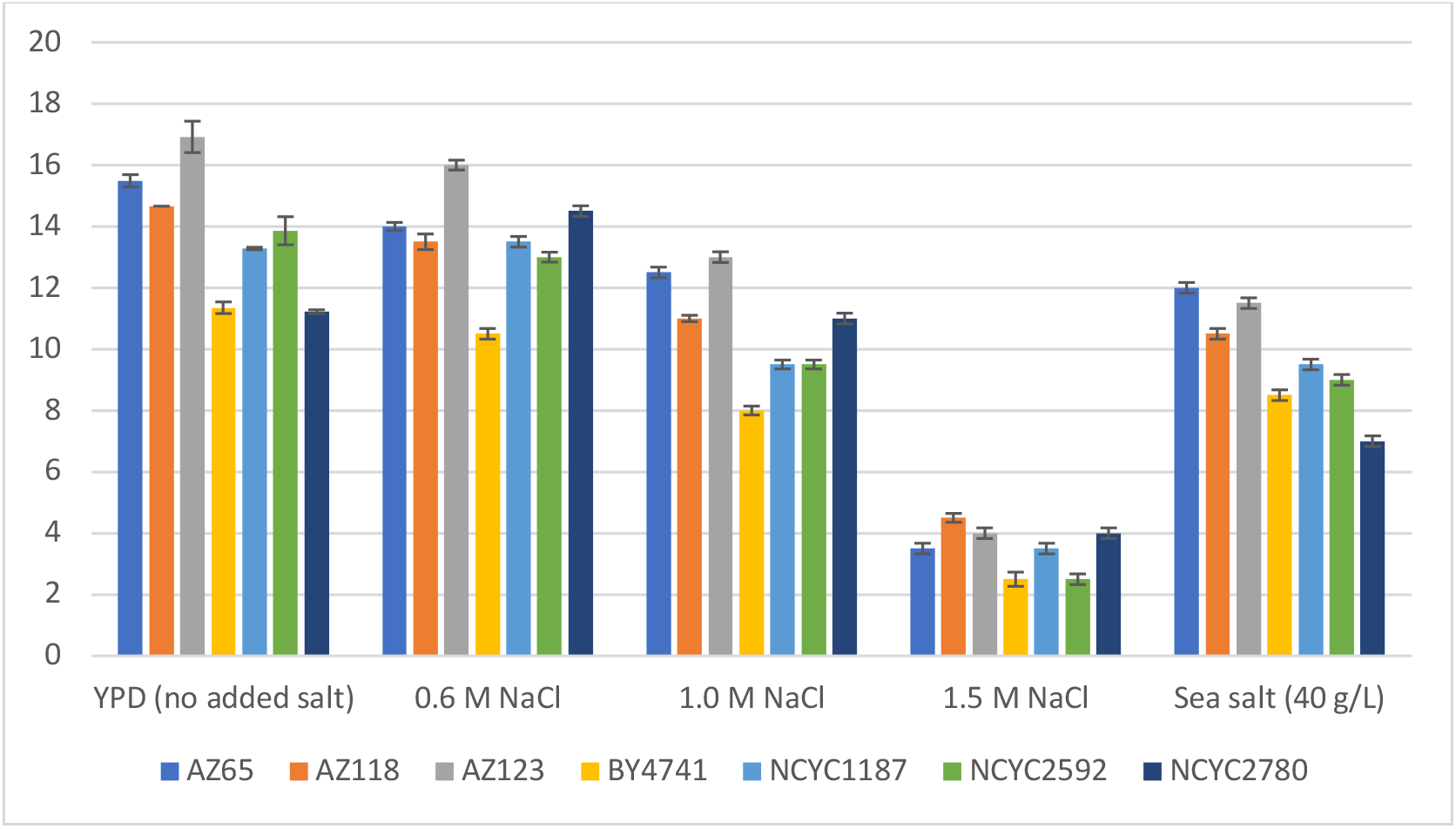
Salt tolerance of marine (AZ65, AZ118, AZ123) and comparator (BY4741, NCYC1187, NCYC2592, and NCYC2780) *S. cerevisiae* strains. Bars show OD (600 nm) at 72 h in YPD medium with the indicated amendment. All cultures were grown at 30°C with shaking at 200 rpm. Error bars indicate standard deviation from three biological replicates.

Lignocellulosic bioethanol production exposes fermenting yeasts to furan aldehydes and weak acids generated during biomass pre-treatment. To determine whether the marine isolates show broad stress tolerance beyond salinity, growth was assessed in YPD supplemented with HMF (15 mM), furfural (10 mM), or acetic acid (60 mM; pH 5.5) for 72 h at 30°C (Figure 2). One-way ANOVA revealed significant strain effects under all three inhibitor conditions (p < 0.001), and Tukey’s HSD post-hoc tests confirmed that marine isolates AZ65, AZ118, and AZ123 grew significantly better than BY4741 under all conditions (HMF: p < 0.01; furfural: p < 0.01; acetic acid: p < 0.05). AZ123 ranked highest overall, reaching OD600 = 16.92 ± 1.69 in the uninhibited glucose control, with OD600 = 14.16 ± 0.48 under acetic acid and 13.85 ± 0.89 under furfural; under HMF, however, AZ65 was marginally higher (12.03 ± 0.61) than AZ123 (11.39 ± 0.33), with AZ118 close behind (10.60 ± 0.55).

**Figure 2:**
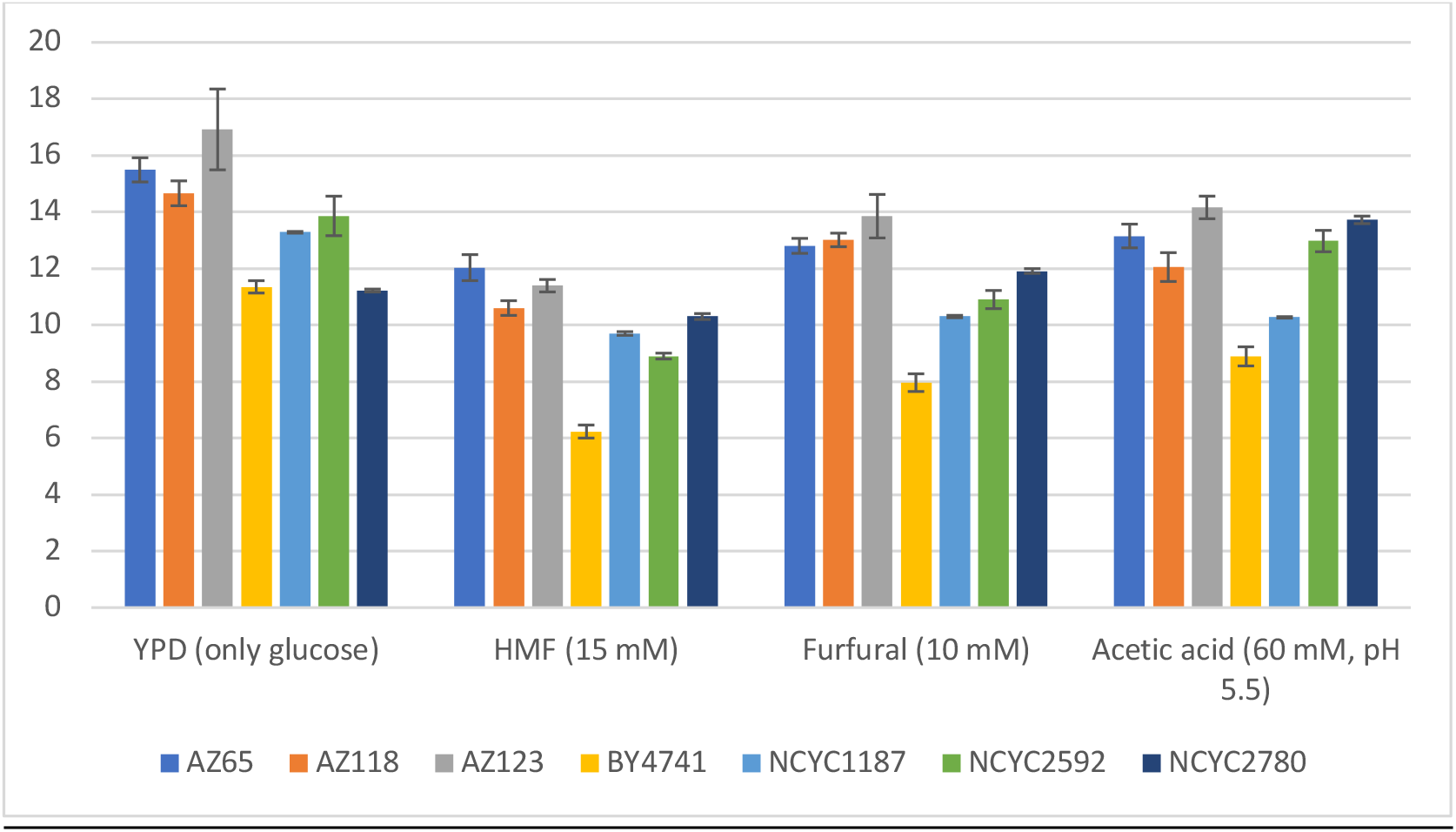
Inhibitor tolerance of marine (AZ65, AZ118, AZ123) and comparator (BY4741, NCYC1187, NCYC2592, and NCYC2780) *S. cerevisiae* strains. Bars show OD (600 nm) at 72 h in YPD medium with the indicated amendment. All cultures were grown at 30°C with shaking at 200 rpm. Error bars indicate standard deviation from three biological replicates.

AZ65 and AZ118 performed strongly under furfural and acetic acid (OD600 12.05–13.15). NCYC2780 showed selective acetic acid resilience (13.72 ± 0.08), comparable to AZ65, while performing less well under HMF (10.30 ± 0.15) and furfural (11.91 ± 0.13). The other two industrial references, NCYC2592 and NCYC1187, showed intermediate inhibitor tolerance below the marine isolates (HMF 8.90 ± 0.19 and 9.70 ± 0.11; furfural 10.90 ± 0.43 and 10.30 ± 0.07; acetic acid 12.97 ± 0.48 and 10.27 ± 0.04, respectively); AZ123 significantly outperformed both under HMF and furfural (p < 0.05). BY4741 showed the poorest performance across all inhibitors (HMF 6.23 ± 0.37; furfural 7.96 ± 0.45; acetic acid 8.89 ± 0.47), consistent with its laboratory domestication and known stress sensitivity. Taken together with the salt tolerance data, these results establish that AZ123’s phenotypic superiority extends across multiple stressor classes, providing a broader foundation for its proposed role as an industrial fermentation strain.

### 3.2 Genome sequencing

Genomic DNA was prepared from strains AZ65, AZ118 and AZ123 and sequenced using a combination of long read (Nanopore) and short-read (Illumina) sequencing. Nanopore sequencing yielded 1.2–1.6 Gb per strain at 99–134× estimated coverage. Hybrid assemblies ranged from 12.1 to 12.2 Mb with contig N50 values of 537–617 kb (Table 1). BUSCO analysis confirmed >99% completeness for all three genomes, indicating near-complete assemblies suitable for locus-level comparative analysis.

**Table 1.**
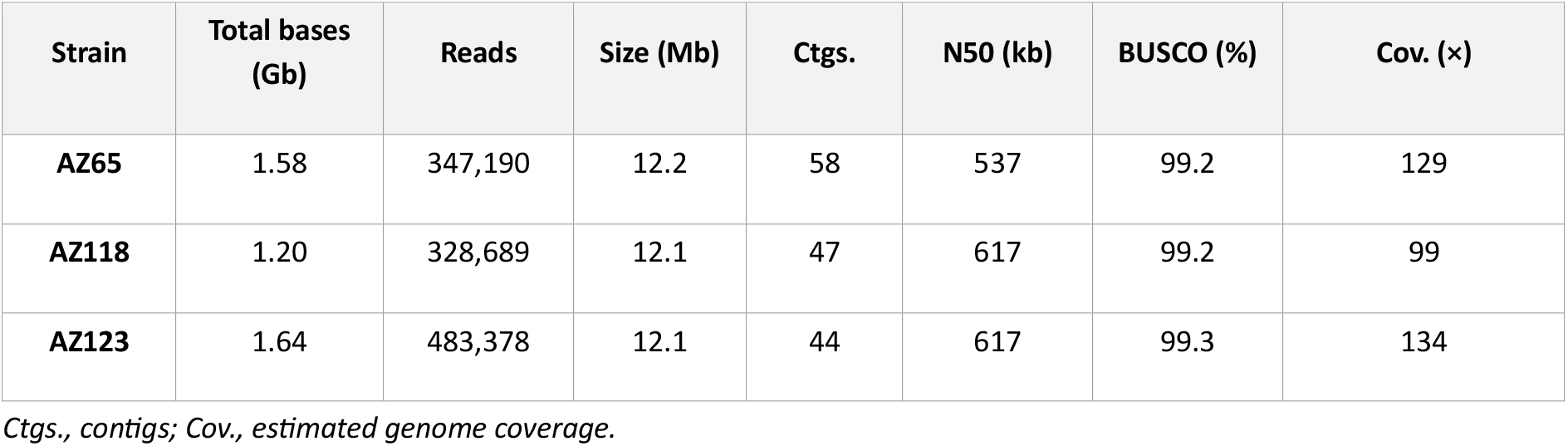
Sequencing output and genome assembly statistics for three marine *S. cerevisiae* strains.

### 3.3 Analysis of the *ENA* locus

Contigs bearing genes matching the sequences of *ENA1, ENA2, ENA5, RSM10* or *KRS1* were located using standard sequence comparisons. Results showed that AZ65 and AZ118 appeared to possess two different versions of this locus (Figure 3), possibly indicating that these strains are heterozygous diploids, whereas strain AZ123 showed only one version of this locus. The assembly for AZ65 showed a long locus with 4 copies of *ENA* (designated AZ65A1, AZ65A2, AZ65A3 and AZ65A4) located adjacent to *KRS1*, but upstream DNA including the *ENA1* promoter and *RSM10* was not present on this contig. The alternative contig had a single *ENA* copy (AZ65B1) between *RSM10* and *KRS1*. One simple interpretation is that this strain is heterozygous at this locus, with the two copies identical at *RSM10* but different in the *ENA* and *KRS1* regions. The assembly for AZ118 showed a long locus with two copies (AZ118A1 and AZ118A2), flanked by *RSM10* and *KRS1*, and a second copy (AZ118B1) in a contig on its own, lacking flanking sequences; this may represent a second allele of one of the two copies (most likely the second; see below for discussion) but identical in all other regions. The assembly for AZ123 showed a single locus similar to that of AZ118, with two copies of *ENA* flanked by *RSM10* and *KRS1*.

**Figure 3:**
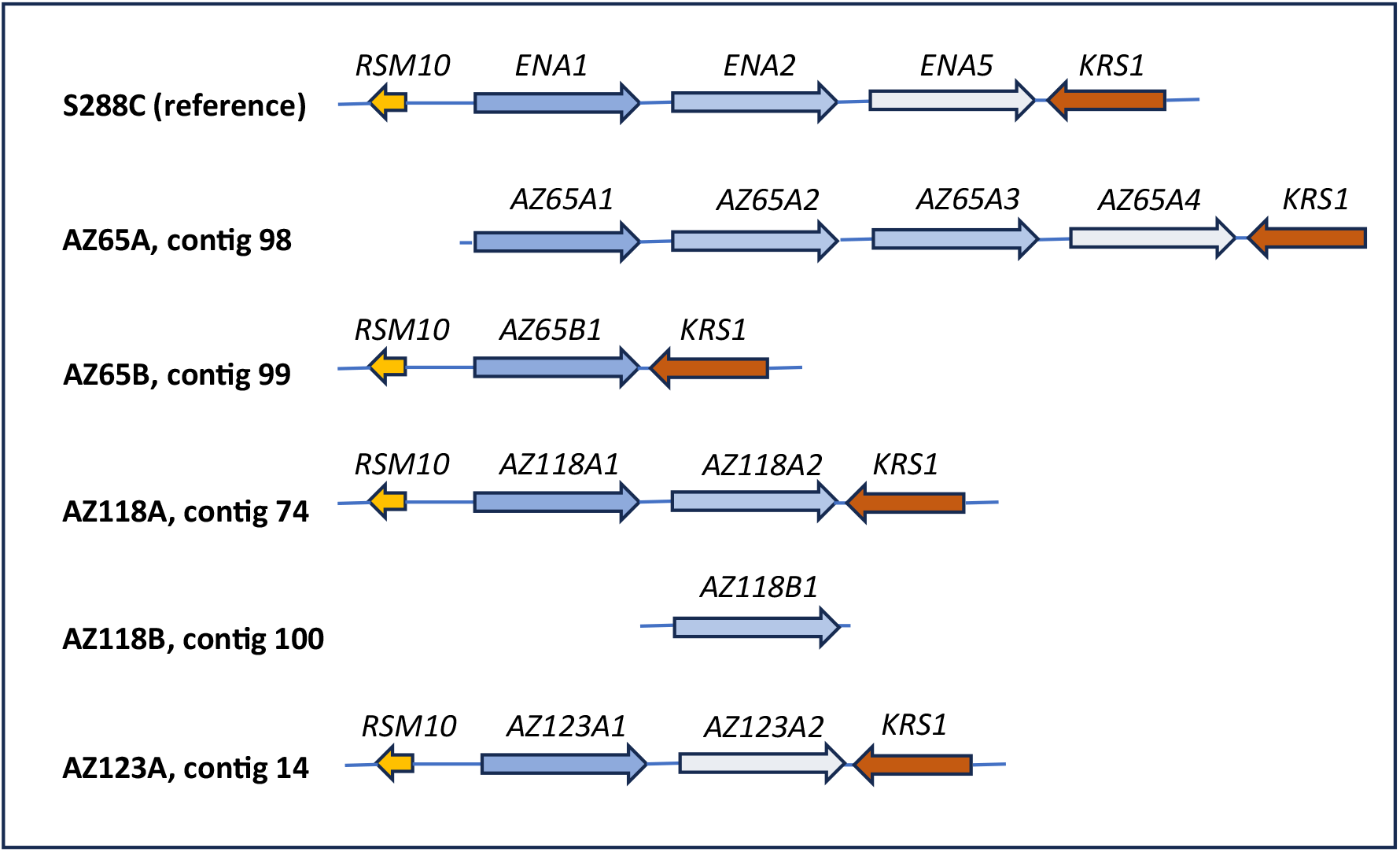
Arrangement of *ENA* loci detected in genome sequences. The typical flanking genes of the *ENA* locus on chromosome IV, *RMS10* and *KRS1*, are shown in orange and red where they were present on a given contig. *ENA* copies are shown in shades of blue, with *ENA1*-like copies darkest, *ENA2*-like copies medium blue, and *ENA5*-like copies in lightest blue. Copies are named according to strain (AZ65, AZ118 or AZ123), variant (A or B) and position within the locus (1, 2, 3 or 4). Where flanking genes are not present on a given contig, it is presumed that these were present in the genome but that the sequence was identical to that of an alternative locus, and no reads spanned this join; see Discussion for details.

**Figure 4:**
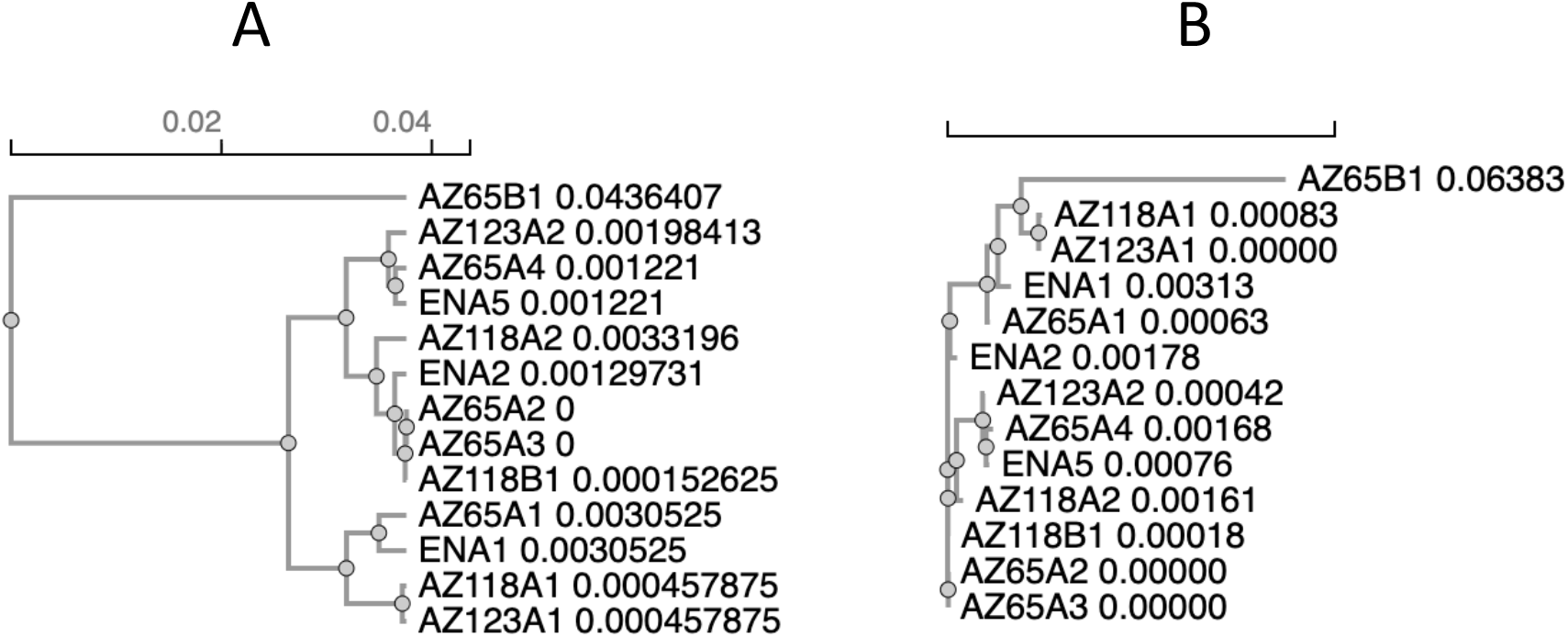
Similarity among *ENA* copies. **A: sequence similarity tree**. The sequences fall into three clusters representing (from top) *ENA5*-like, *ENA2*-like and *ENA1*-like sequences, with AZ65B1 as a divergent outlier. **B: inferred evolutionary tree**. The topology is generally similar, but AZ65B1 now groups with the *ENA1*-like sequences, suggesting that it is an unusually divergent *ENA1*-like gene.

Analysis of the coding sequences of all *ENA* copies, and comparison to the sequences of *ENA1, ENA2* and *ENA5* from S288C, showed a very high degree of sequence conservation (Supplementary Figure S1). Based on sequence similarity, the genes fall into three groups. The first gene in each locus (AZ65A1, AZ118A1 and AZ123A1) was most similar to *ENA1*. The single *ENA* copy in the second AZ65 locus, AZ65B1, was substantially divergent and represented an outgroup to all other *ENA* copies in simple sequence-comparison trees, but grouped with *ENA1* in evolutionary trees, suggesting that it is a highly derived version of *ENA1*, or a pseudogene derived from it. Of the other copies, AZ65A2, AZ65A3 and AZ118A2 resembled *ENA2*, and AZ65A4 and AZ123A2 resembled *ENA5. ENA2* and *ENA5* were also more similar to each other than to *ENA1*. These similarities are consistent with a simplified evolutionary hypothesis in which an initial duplication of *ENA1* gave rise to *ENA2*, including a novel promoter sequence, and a subsequent duplication of *ENA2* gave rise to *ENA5*, with an identical promoter sequence. Under this hypothesis, further duplication of *ENA2* could have produced the four-copy locus of AZ65, whereas the two-copy loci of AZ118 and AZ123 could reflect loss of *ENA2* and *ENA5*, respectively.

It should be noted that several of the sequences (AZ65B1, AZ118A2 and AZ123A2) contain single base insertions at different points which would result in reading frame errors (Supplementary Figure S1) and, presumably, non-functional protein products. These apparent frameshifts may reflect genuine sequence changes, or sequencing or assembly errors. To clarify this point, locus AZ65B1, which showed two single-base insertions leading to frameshift errors, was amplified by PCR from genomic DNA, cloned in a plasmid vector and sequenced. Both of the single base insertions in the genome sequence were absent from the amplified and cloned sequence (data not shown). This suggests that AZ65B1 is in fact a potentially functional ENA copy, and that the insertions seen in this and other copies are sequencing artefacts. Based on this assumption, the five apparent single base insertions in the 13 sequences were manually corrected to give the maximum possible sequence conservation, and the sequences translated to give putative protein sequences (Supplementary Figure S2). As expected, these proteins all had high similarity to ENA1, ENA2 and ENA5 of S288C, and similarity trees generated from the protein sequences were essentially identical in topology to those generated from the original DNA sequences.

Examination of the protein sequences (Supplementary Figure S2) revealed that the great majority of the variation occurred in the first 110 amino acids, in which the *ENA1*-like sequences differed from the *ENA2/ENA5*-like sequences, and in the last 33 amino acids, in which the *ENA5*-like sequences differed from the *ENA1/ENA2*-like sequences. In between these two regions there were no systematic differences. AZ65B1 had the largest number of unique variants, accounting for its divergent position, but was *ENA1*-like in the diagnostic N-terminal region. Thus we see that *ENA1*-like sequences, generally the first in the locus, have a unique N-terminal signature, whereas *ENA5*-like sequences, generally coming last in the locus, have a unique C-terminal signature. In all other regions, the sequences show very high identity.

Analysis of the upstream promoter, corresponding to the *ENA1* promoter in S288C (that is to say, the region between the start codons of *RSM10* and the first copy of *ENA*), showed very high sequence conservation among all the strains, with only a few nucleotide changes (Supplementary Figure S3). The only substantial difference was a decrease in the length of a TA repeat located at position -285 to -266 relative to the start of *ENA1*. While it is conceivable that this might alter the strength of the promoter or the interactions of the various transcription factors which control expression of this gene, there seem to be no substantial sequence differences affecting the known transcription factor binding sites of the *ENA1* promoter. It should be noted that the long version of the AZ65 locus (AZ65A) and the short version of the AZ118 locus (AZ118B) could not be included in this analysis since upstream DNA was not present on those contigs. However, the most likely explanation for this is that the upstream sequence is identical to that of the other locus; in this case the promoter of AZ65A1 would be identical to that of AZ65B1, and that of AZ118B1 to that of AZ118A1.

Analysis of the intergenic regions which presumably represent the promoters of the second and subsequent *ENA* copies in each locus revealed that the 609 bp intergenic sequence upstream of each secondary copy was completely conserved in all strains and all *ENA* copies examined (Supplementary Figure S4). While we have been unable to find any published analysis of these promoter regions in any strain of *S. cerevisiae*, the fact that they are identical in all strains examined suggests that variation in this region cannot be responsible for alterations in salt tolerance.

## 4. Discussion

The increasing interest in seawater-based bioprocesses, together with the commonly seen increase in stress tolerance seen in marine organisms compared to terrestrial ones, make examination of marine strains of the dominant industrial and research organism *Saccharomyces cerevisiae* a matter of considerable interest. Here we present an analysis of salt and inhibitor tolerance in three marine *S. cerevisiae* strains, AZ65, AZ118 and AZ123, together with their genome sequences and a preliminary analysis of the *ENA* locus, the most obvious potential source of changes in salt tolerance.

The ability of the marine isolates to sustain measurable growth at 1.5 M NaCl is notable in the context of natural variation across the species. In a survey of 1,011 *S. cerevisiae* isolates, Peter et al. (2018) found that measurable growth at this concentration is rare, with the great majority of strains severely inhibited. Against that background, the marine isolates characterised here (OD600 = 3.5–4.5 at 1.5 M NaCl) rank among the more halotolerant natural isolates reported. In inhibitor tolerance experiments, the marine strains showed increased tolerance to acetate, furfural and HMF stress, consistent with results previously reported by Greetham et al. (2019). We have also generated a *URA3* mutant of AZ123 and demonstrated that it can be transformed with plasmids leading to production of violacein and beta-carotene (data not shown), supporting its potential usefulness in bioprocesses beyond ethanol production.

The genomes of the three marine strains were sequenced using a combination of long read (Nanopore) and short read (Sanger) sequencing to generate high-quality genome sequences. YGAP analysis (Proux-Wéra et al, 2012) showed a high degree of synteny with other *S. cerevisiae* strains (data not shown), and ITS sequence analysis confirmed the previous finding that these strains are all *S. cerevisiae*. In all respects, these genomes conformed to expectations for new *S. cerevisiae* isolates.

Peter et al (2018) reported a correlation between copy number of *ENA* genes and salt tolerance in 1,011 *S. cerevisiae* strains; in fact, they reported that this was one of the very few clear associations between genome sequence and any physiological characteristic. We therefore predicted that our marine strains would have increased copy number at this locus. However, contrary to our expectation, this did not appear to be the case. Whereas the reference sequence from S288C shows three copies of *ENA* at this locus, our sequences showed 2 to 4 copies, with the strain showing highest salt tolerance, AZ123, clearly having two copies. AZ65 appeared to have two versions of the locus, one having 4 copies, apparently due to a duplication of *ENA2*, and one having a single copy. Both versions were adjacent to *KRS1* and showed the expected downstream sequence; the contig bearing the long version was lacking upstream sequence. This suggests that the strain may be a diploid which is heterozygous at this locus, thus having 5 copies of *ENA* in its diploid genome. If AZ123 is also diploid, it would have 4 copies of *ENA*, and in this case, AZ118 would most likely also have 4 copies, with the second, single-copy version representing a variant allele of one of the two copies in the other locus, most likely *ENA2*. Considering that wild terrestrial strains can have up to 6 copies per haploid genome, increased *ENA* copy number is not clearly supported as the basis for increased salt tolerance in marine strains.

The promoter region of *ENA1* is well studied. The corresponding sequences in the marine genomes aligned well with the reference sequence, showing only a few small differences which did not seem to relate to the known transcription factor-binding sites in this region. A TA-repeat was shorter in the marine strains than in the reference genome, but it is not clear what effect, if any, this might have on expression. Thus, alterations to this promoter do not provide an obvious explanation of increased salt tolerance in our strains. As for the promoters of the second and subsequent *ENA* copies at each locus, these were all identical to each other and to those in the reference genome; while the regulation of this promoter seems to be little-studied, there is no observed variation that explains changes in salt tolerance.

Overall, we found no clear evidence that the *ENA* locus is a major source of differences in salt or inhibitor tolerance in our marine-derived isolates compared with terrestrial and industrial comparator strains. Thus, the genetic and physiological basis of salt and inhibitor tolerance requires further analysis. We hope that further study of these genome sequences, together with other marine and terrestrial genome sequences, will yield insights that may allow us to increase salt and inhibitor tolerance in valuable industrial strains of *S. cerevisiae*.

## Supporting information

AZ65 contigs in fasta format

AZ118 contigs in fasta format

AZ123 contigs in fasta format

Data for Figures 1 and 2

