## Supplemental Figure 1 for "Analysis of genome sequences of marine isolates of *Saccharomyces cerevisiae*"

**Supplementary Figure S1: alignment of sequences of all *ENA* copies detected in marine strains AZ65, AZ118 and AZ123, together with *ENA1*, *ENA2* and *ENA5* of the reference strain S288C.** Start and stop codons are highlighted in green. Apparent indels which could cause frame shifts are highlighted in cyan.

```
AZ65B1      ATGGGCGAAGGAACTACTAAGGAAAACAATAATGCAGAATTCAATGCTTATCACACGCTG	60
AZ123A2     ATGAGCGAGGGAACTGTCAAAGAAAACAATAATGAAGAATTCAATGCTTATCACACATTG	60
AZ65A4      ATGAGCGAGGGAACTGTCAAAGAAAACAATAATGAAGAATTCAATGCTTATCACACATTG	60
ENA5        ATGAGCGAGGGAACTGTCAAAGAAAACAATAATGAAGAATTCAATGCTTATCACACATTG	60
AZ118A2     ATGAGCGAGGGAACTGTCAAAGAAAACAATAATGAAGAATTCAATGCTTATCACACATTG	60
ENA2        ATGAGCGAGGGAACTGTCAAAGAAAACAATAATGAAGAATTCAATGCTTATCACACATTG	60
AZ65A2      ATGAGCGAGGGAACTGTCAAAGAAAACAATAATGAAGAATTCAATGCTTATCACACATTG	60
AZ65A3      ATGAGCGAGGGAACTGTCAAAGAAAACAATAATGAAGAATTCAATGCTTATCACACATTG	60
AZ118B1     ATGAGCGAGGGAACTGTCAAAGAAAACAATAATGAAGAATTCAATGCTTATCACACATTG	60
AZ65A1      ATGGGCGAAGGAACTACTAAGGAAAACAATAATGCAGAATTCAATGCTTATCACACGCTG	60
ENA1        ATGGGCGAAGGAACTACTAAGGAAAACAATAATGCAGAATTCAATGCTTATCACACGCTG	60
AZ118A1     ATGGGCGAAGGAACTACTAAGGAAAACAATAATGCAGAATTCAATGCTTATCACACGCTG	60
AZ123A1     ATGGGCGAAGGAACTACTAAGGAAAACAATAATGCAGAATTCAATGCTTATCACACGCTG	60
            *** **** ******   ** ************* *********************  **

AZ65B1      ACTGCAGAAGAAGCCGCTGAATTCATAGGCACAAGTCTAACTGAAGGTTTGACCCAAGAT	120
AZ123A2     ACTACAGAAGAAGCCGCTGAATTTATAGGCACAAGTCTGACTGAAGGTCTTACTCAAGAT	120
AZ65A4      ACTACAGAAGAAGCCGCTGAATTTATAGGCACAAGTCTGACTGAAGGTCTTACTCAAGAT	120
ENA5        ACTACAGAAGAAGCCGCTGAATTTATAGGCACAAGTCTGACTGAAGGTCTTACTCAAGAT	120
AZ118A2     ACTACAGAAGAAGCCGCTGAATTTATAGGCACAAGTCTGACTGAAGGTCTTACTCAAGAT	120
ENA2        ACTACAGAAGAAGCCGCTGAATTTATAGGCACAAGTCTGACTGAAGGTCTTACTCAAGAT	120
AZ65A2      ACTACAGAAGAAGCCGCTGAATTTATAGGCACAAGTCTGACTGAAGGTCTTACTCAAGAT	120
AZ65A3      ACTACAGAAGAAGCCGCTGAATTTATAGGCACAAGTCTGACTGAAGGTCTTACTCAAGAT	120
AZ118B1     ACTACAGAAGAAGCCGCTGAATTTATAGGCACAAGTCTGACTGAAGGTCTTACTCAAGAT	120
AZ65A1      ACTGCAGAAGAAGCCGCTGAATTCATAGGCACAAGTCTAACTGAAGGTTTGACCCAAGAT	120
ENA1        ACTGCAGAAGAAGCCGCTGAATTCATAGGCACAAGTCTAACTGAAGGTTTGACCCAAGAT	120
AZ118A1     ACTGCAGAAGAAGCCGCTGAATTCATAGGCACAAGTCTAACTGAAGGTTTGACCCAAGAT	120
AZ123A1     ACTGCAGAAGAAGCCGCTGAATTCATAGGCACAAGTCTAACTGAAGGTTTGACCCAAGAT	120
            *** ******************* ************** ********* * ** ******

AZ65B1      GAGTTCGTCCACAGATTGAAAACAGTGGGTGAGAACACATTGGGTGATGACACTAAAATT	180
AZ123A2     GAGTCCCTCCGCAGACTGAAAGCAGTGGGAGAGAACACATTGGGTGATGATACCAAAATT	180
AZ65A4      GAGTCCCTCCGCAGACTGAAAGCAGTGGGAGAGAACACATTGGGTGATGATACCAAAATT	180
ENA5        GAGTCCCTCCGCAGACTGAAAGCAGTGGGAGAGAACACATTGGGTGATGATACCAAAATT	180
AZ118A2     GAGTCCCTCCGCAGACTGAAAGCAGTGGGAGAGAACACATTGGGTGATGATACCAAAATT	180
ENA2        GAGTCCCTCCGCAGACTGAAAGCAGTGGGAGAGAACACATTGGGTGATGATACCAAAATT	180
AZ65A2      GAGTCCCTCCGCAGACTGAAAGCAGTGGGAGAGAACACATTGGGTGATGATACCAAAATT	180
AZ65A3      GAGTCCCTCCGCAGACTGAAAGCAGTGGGAGAGAACACATTGGGTGATGATACCAAAATT	180
AZ118B1     GAGTCCCTCCGCAGACTGAAAGCAGTGGGAGAGAACACATTGGGTGATGATACCAAAATT	180
AZ65A1      GAGTTCGTCCACAGATTGAAAACAGTGGGTGAGAACACATTGGGTGATGACACTAAAATT	180
ENA1        GAGTTCGTCCACAGATTGAAAACAGTGGGTGAGAACACATTGGGTGATGACACTAAAATT	180
AZ118A1     GAGTTCGTCCACAGATTGAAAACAGTGGGTGAGAACACATTGGGTGATGACACTAAAATT	180
AZ123A1     GAGTTCGTCCACAGATTGAAAACAGTGGGTGAGAACACATTGGGTGATGACACTAAAATT	180
            **** * *** **** ***** ******* ******************** ** ******
```

```
AZ65B1      GATTACAAGGCAATGGTCCTCCATCAGGTATGTAATGCCATGATCATGGTCCTTCTAATA	240
AZ123A2     GACTATAAAGCAATGGTCCTCCATCAGGTATGTAATGCAATGATTATGGTCTTGGTAATA	240
AZ65A4      GACTATAAAGCAATGGTCCTCCATCAGGTATGTAATGCAATGATTATGGTCTTGGTAATA	240
ENA5        GACTATAAAGCAATGGTCCTCCATCAGGTATGTAATGCAATGATTATGGTCTTGGTAATA	240
AZ118A2     GACTATAAAGCAATGGTCCTCCATCAGGTATGTAATGCAATGATTATGGTCTTGGTAATA	240
ENA2        GACTATAAAGCAATGGTCCTCCATCAGGTATGTAATGCAATGATTATGGTCTTGGTAATA	240
AZ65A2      GACTATAAAGCAATGGTCCTCCATCAGGTATGTAATGCAATGATTATGGTCTTGGTAATA	240
AZ65A3      GACTATAAAGCAATGGTCCTCCATCAGGTATGTAATGCAATGATTATGGTCTTGGTAATA	240
AZ118B1     GACTATAAAGCAATGGTCCTCCATCAGGTATGTAATGCAATGATTATGGTCTTGGTAATA	240
AZ65A1      GATTACAAGGCAATGGTCCTCCATCAGGTATGTAATGCCATGATCATGGTCCTTCTAATA	240
ENA1        GATTACAAGGCAATGGTCCTCCATCAGGTATGTAATGCCATGATCATGGTCCTTCTAATA	240
AZ118A1     GATTACAAGGCAATGGTCCTCCATCAGGTATGTAATGCCATGATCATGGTCCTTCTAATA	240
AZ123A1     GATTACAAGGCAATGGTCCTCCATCAGGTATGTAATGCCATGATCATGGTCCTTCTAATA	240
            ** ** ** ***************************** ***** ****** *  *****

AZ65B1      TCCATGATAATCTCGTTTGCCATGCATGATTGGATTACTGGTGGTGTTATTTCCTTTGTC	300
AZ123A2     TCCATGGCGATCTCTTTCGCTGTGCGCGACTGGATTACTGGTGGCGTTATTTCTTTTGTT	300
AZ65A4      TCCATGGCGATCTCTTTCGCTGTGCGCGACTGGATTACTGGTGGCGTTATTTCTTTTGTT	300
ENA5        TCCATGGCGATCTCTTTCGCTGTGCGCGACTGGATTACTGGTGGCGTTATTTCTTTTGTT	300
AZ118A2     TCCATGGCGATCTCTTTCGCTGTGCGCGACTGGATTACTGGTGGCGTTATTTCTTTTGTT	300
ENA2        TCCATGGCGATCTCTTTCGCTGTGCGCGACTGGATTACTGGTGGCGTTATTTCTTTTGTT	300
AZ65A2      TCCATGGCGATCTCTTTCGCTGTGCGCGACTGGATTACTGGTGGCGTTATTTCTTTTGTT	300
AZ65A3      TCCATGGCGATCTCTTTCGCTGTGCGCGACTGGATTACTGGTGGCGTTATTTCTTTTGTT	300
AZ118B1     TCCATGGCGATCTCTTTCGCTGTGCGCGACTGGATTACTGGTGGCGTTATTTCTTTTGTT	300
AZ65A1      TCCATGGCGATCTCTTTCGCTGTGCGCGACTGGATTACTGGTGGCGTTATTTCTTTTGTT	300
ENA1        TCCATGATAATCTCGTTTGCCATGCATGATTGGATTACTGGTGGCGTTATTTCTTTTGTT	300
AZ118A1     TCCATGATAATCTCGTTTGCCATGCATGATTGGATTACTGGTGGTGTTATTTCCTTTGTC	300
AZ123A1     TCCATGATAATCTCGTTTGCCATGCATGATTGGATTACTGGTGGTGTTATTTCCTTTGTC	300
            ******   ***** ** **  ***  ** ************** ******** *****

AZ65B1      GTGGGTGTTAACGTCGGTATTGGCCTTGTTCAAGAATATAAGGCTACAAAGACAATGAAC	360
AZ123A2     ATCGCGGTCAATGTGCTCATTGGCCTAGTTCAAGAATATAAGGCTACCAAGACAATGAAC	360
AZ65A4      ATCGCGGTCAATGTGCTCATTGGCCTAGTTCAAGAATATAAGGCTACCAAGACAATGAAC	360
ENA5        ATCGCGGTCAATGTGCTCATTGGCCTAGTTCAAGAATATAAGGCTACCAAGACAATGAAC	360
AZ118A2     ATCGCGGTCAATGTGCTCATTGGCCTAGTTCAAGAATATAAGGCTACCAAGACAATGAAC	360
ENA2        ATCGCGGTCAATGTGCTCATTGGCCTAGTTCAAGAATATAAGGCTACCAAGACAATGAAC	360
AZ65A2      ATCGCGGTCAATGTGCTCATTGGCCTAGTTCAAGAATATAAGGCTACCAAGACAATGAAC	360
AZ65A3      ATCGCGGTCAATGTGCTCATTGGCCTAGTTCAAGAATATAAGGCTACCAAGACAATGAAC	360
AZ118B1     ATCGCGGTCAATGTGCTCATTGGCCTAGTTCAAGAATATAAGGCTACCAAGACAATGAAC	360
AZ65A1      ATCGCGGTCAATGTGCTCATTGGCCTAGTTCAAGAATATAAGGCTACCAAGACAATGAAC	360
ENA1        ATCGCGGTCAATGTGCTCATTGGCCTAGTTCAAGAATATAAGGCTACCAAGACAATGAAC	360
AZ118A1     GTGGGTGTTAACGTCGGTATTGGCCTTGTTCAAGAATATAAGGCTACAAAGACAATGAAC	360
AZ123A1     GTGGGTGTTAACGTCGGTATTGGCCTTGTTCAAGAATATAAGGCTACAAAGACAATGAAC	360
             * *  ** ** **    ******** ******************** ************

AZ65B1      TCTTTGAAAAACTTGAGCTCTCCCAATGCTCATGTTATTAGGAACGGGAAAAGTGAGACT	420
AZ123A2     TCTTTGAAAAACTTGAGCTCTCCCAATGCTCATGTTATTAGGAACGGGAAAAGTGAGACT	420
AZ65A4      TCTTTGAAAAACTTGAGCTCTCCCAATGCTCATGTTATTAGGAACGGGAAAAGTGAGACT	420
ENA5        TCTTTGAAAAACTTGAGCTCTCCCAATGCTCATGTTATTAGGAACGGGAAAAGTGAGACT	420
AZ118A2     TCTTTGAAAAACTTGAGCTCTCCCAATGCTCATGTTATTAGGAACGGGAAAAGTGAGACT	420
ENA2        TCTTTGAAAAACTTGAGCTCTCCCAATGCTCATGTTATTAGGAACGGGAAAAGTGAGACT	420
AZ65A2      TCTTTGAAAAACTTGAGCTCTCCCAATGCTCATGTTATTAGGAACGGGAAAAGTGAGACT	420
AZ65A3      TCTTTGAAAAACTTGAGCTCTCCCAATGCTCATGTTATTAGGAACGGGAAAAGTGAGACT	420
AZ118B1     TCTTTGAAAAACTTGAGCTCTCCCAATGCTCATGTTATTAGGAACGGGAAAAGTGAGACT	420
AZ65A1      TCTTTGAAAAACTTGAGCTCTCCCAATGCTCATGTTATTAGGAACGGGAAAAGTGAGACT	420
ENA1        TCTTTGAAAAACTTGAGCTCTCCCAATGCTCATGTTATTAGGAACGGGAAAAGTGAGACT	420
AZ118A1     TCTTTGAAAAACTTGAGCTCTCCCAATGCTCATGTTATCAGAAACGGGAAGAGCGAAACT	420
AZ123A1     TCTTTGAAAAACTTGAGCTCTCCCAATGCTCATGTTATCAGAAACGGGAAGAGCGAAACT	420
            ************************************** ** ******** ** ** ***
AZ65B1      ATAAACTCCAAAGATGTGGTTCCAGGTGATATTTGTCTGGTGAAGGTCGGTGATACTATT	480
AZ123A2     ATAAACTCAAAAGATGTGGTTCCAGGTGATATTTGTCTGGTGAAGGTCGGTGATACTATT	480
AZ65A4      ATAAACTCAAAAGATGTGGTTCCAGGTGATATTTGTCTGGTGAAGGTCGGTGATACTATT	480
ENA5        ATAAACTCAAAAGATGTGGTTCCAGGTGATATTTGTCTGGTGAAGGTCGGTGATACTATT	480
AZ118A2     ATAAACTCAAAAGATGTGGTTCCAGGTGATATTTGTCTGGTGAAGGTCGGTGATACTATT	480
ENA2        ATAAACTCAAAAGATGTGGTTCCAGGTGATATTTGTCTGGTGAAGGTCGGTGATACTATT	480
AZ65A2      ATAAACTCAAAAGATGTGGTTCCAGGTGATATTTGTCTGGTGAAGGTCGGTGATACTATT	480
AZ65A3      ATAAACTCAAAAGATGTGGTTCCAGGTGATATTTGTCTGGTGAAGGTCGGTGATACTATT	480
AZ118B1     ATAAACTCAAAAGATGTGGTTCCAGGTGATATTTGTCTGGTGAAGGTCGGTGATACTATT	480
AZ65A1      ATAAACTCAAAAGATGTGGTTCCAGGTGATATTTGTCTGGTGAAGGTCGGTGATACTATT	480
ENA1        ATAAACTCAAAAGATGTGGTTCCAGGTGATATTTGTCTGGTGAAGGTCGGTGATACTATT	480
AZ118A1     ATAAATTCCAAAGATGTCGTTCCAGGTGATATTTGTCTAGTGAAGGTCGGTGACACTATT	480
AZ123A1     ATAAATTCCAAAGATGTCGTTCCAGGTGATATTTGTCTAGTGAAGGTCGGTGACACTATT	480
            ***** ** ******** ******************** ************** ******

AZ65B1      CCTGCTGATTTGCGTTTGATTGAAACTAAGAATTTTGATACTGATGAATCACTGCTGACC	540
AZ123A2     CCTGCTGATTTGCGTTTAATTGAAACTAAGAATTTTGATACTGATGAATCACTACTGACC	540
AZ65A4      CCTGCTGATTTGCGTTTAATTGAAACTAAGAATTTTGATACTGATGAATCACTACTGACC	540
ENA5        CCTGCTGATTTGCGTTTAATTGAAACTAAGAATTTTGATACTGATGAATCACTACTGACC	540
AZ118A2     CCTGCTGATTTGCGTTTAATTGAAACTAAGAATTTTGATACTGATGAATCACTACTGACC	540
ENA2        CCTGCTGATTTGCGTTTAATTGAAACTAAGAATTTTGATACTGATGAATCACTACTGACC	540
AZ65A2      CCTGCTGATTTGCGTTTAATTGAAACTAAGAATTTTGATACTGATGAATCACTACTGACC	540
AZ65A3      CCTGCTGATTTGCGTTTAATTGAAACTAAGAATTTTGATACTGATGAATCACTACTGACC	540
AZ118B1     CCTGCTGATTTGCGTTTAATTGAAACTAAGAATTTTGATACTGATGAATCACTACTGACC	540
AZ65A1      CCTGCTGATTTGCGTTTAATTGAAACTAAGAATTTTGATACTGATGAATCACTACTGACC	540
ENA1        CCTGCTGATTTGCGTTTAATTGAAACTAAGAATTTTGATACTGATGAATCACTACTGACC	540
AZ118A1     CCTGCTGATTTACGTTTGATTGAAACCAAGAATTTTGATACTGATGAATCACTGCTGACC	540
AZ123A1     CCTGCTGATTTACGTTTGATTGAAACCAAGAATTTTGATACTGATGAATCACTGCTGACC	540
            *********** ***** ******** ************************** ******

AZ65B1      GGTGAATCTTTGCCTGTCTCCAAAGACGCCGACTTGGTATTTGGAAAAGAAGAAGAAACC	600
AZ123A2     GGTGAATCTTTGCCTGTCTCCAAAGACGCCAACTTAGTGTTTGGAAAAGAAGAAGAAACC	600
AZ65A4      GGTGAATCTTTGCCTGTCTCCAAAGACGCCAACTTAGTGTTTGGAAAAGAAGAAGAAACC	600
ENA5        GGTGAATCTTTGCCTGTCTCCAAAGACGCCAACTTAGTGTTTGGAAAAGAAGAAGAAACC	600
AZ118A2     GGTGAATCTTTGCCTGTCTCCAAAGACGCCAACTTAGTGTTTGGAAAAGAAGAAGAAACC	600
ENA2        GGTGAATCTTTGCCTGTCTCCAAAGACGCCAACTTAGTGTTTGGAAAAGAAGAAGAAACC	600
AZ65A2      GGTGAATCTTTGCCTGTCTCCAAAGACGCCAACTTAGTGTTTGGAAAAGAAGAAGAAACC	600
AZ65A3      GGTGAATCTTTGCCTGTCTCCAAAGACGCCAACTTAGTGTTTGGAAAAGAAGAAGAAACC	600
AZ118B1     GGTGAATCTTTGCCTGTCTCCAAAGACGCCAACTTAGTGTTTGGAAAAGAAGAAGAAACC	600
AZ65A1      GGTGAATCTTTGCCTGTCTCCAAAGACGCCAACTTAGTGTTTGGAAAAGAAGAAGAAACC	600
ENA1        GGTGAATCTTTGCCTGTCTCCAAAGACGCCAACTTAGTGTTTGGAAAAGAAGAAGAAACC	600
AZ118A1     GGTGAATCTTTGCCTGTCTCCAAAGACGCCGACTTGGTGTTTGGAAAAGAAGAAGAAACC	600
AZ123A1     GGTGAATCTTTGCCTGTCTCCAAAGACGCCAACTTAGTGTTTGGAAAAGAAGAAGAAACC	600
            ****************************** **** ** *********************

AZ65B1      TCTGTGGGTGATCGTTTGAATTTAGCATTCTCTTCGTCTACCGTTGTCAAAGGAAGGGCG	660
AZ123A2     TCCGTGGGTGATCGTTTGAATTTAGCATTTTCTTCATCCACCGTTGTCAAGGGAAGAGCC	660
AZ65A4      TCCGTGGGTGATCGTTTGAATTTAGCATTTTCTTCATCCACCGTTGTCAAGGGAAGAGCC	660
ENA5        TCCGTGGGTGATCGTTTGAATTTAGCATTTTCTTCATCCGCCGTTGTCAAGGGAAGAGCC	660
AZ118A2     TCCGTGGGTGATCGTTTGAATTTAGCATTTTCTTCATCCACCGTTGTCAAGGGAAGAGCC	660
ENA2        TCCGTGGGTGATCGTTTGAATTTAGCATTTTCTTCATCCGCCGTTGTCAAGGGAAGAGCC	660
AZ65A2      TCCGTGGGTGATCGTTTGAATTTAGCATTTTCTTCATCCACCGTTGTCAAGGGAAGAGCC	660
AZ65A3      TCCGTGGGTGATCGTTTGAATTTAGCATTTTCTTCATCCACCGTTGTCAAGGGAAGAGCC	660
AZ118B1     TCCGTGGGTGATCGTTTGAATTTAGCATTTTCTTCATCCACCGTTGTCAAGGGAAGAGCC	660
AZ65A1      TCCGTGGGTGATCGTTTGAATTTAGCATTTTCTTCATCCACCGTTGTCAAGGGAAGAGCC	660
ENA1        TCCGTGGGTGATCGTTTGAATTTAGCATTTTCTTCATCCGCCGTTGTCAAGGGAAGAGCC	660
AZ118A1     TCCGTGGGTGATCGTTTGAATTTAGCATTTTCTTCATCCACCGTTGTCAAGGGAAGAGCC	660
AZ123A1     TCCGTGGGTGATCGTTTGAATTTAGCATTTTCTTCATCCACCGTTGTCAAGGGAAGAGCC	660
            ** ************************** ***** **  ********** ***** **
AZ65B1      AAGGGTATTGTCATCAAGACAGCTTTGAACAGTGAAATTGGTAAAATTGCAAAATCTCTA	720
AZ123A2     AAGGGTATTGTCATCAAGACAGCTTTGAATAGTGAAATTGGTAAAATTGCAAAATCTCTA	720
AZ65A4      AAGGGTATTGTCATCAAGACAGCTTTGAATAGTGAAATTGGTAAAATTGCAAAATCTCTA	720
ENA5        AAGGGTATTGTCATCAAGACAGCTTTGAATAGTGAAATTGGTAAAATTGCAAAATCTCTA	720
AZ118A2     AAGGGTATTGTCATCAAGACAGCTTTGAATAGTGAAATTGGTAAAATTGCAAAATCTCTA	720
ENA2        AAGGGTATTGTCATCAAGACAGCTTTGAATAGTGAAATTGGTAAAATTGCAAAATCTCTA	720
AZ65A2      AAGGGTATTGTCATCAAGACAGCTTTGAATAGTGAAATTGGTAAAATTGCAAAATCTCTA	720
AZ65A3      AAGGGTATTGTCATCAAGACAGCTTTGAATAGTGAAATTGGTAAAATTGCAAAATCTCTA	720
AZ118B1     AAGGGTATTGTCATCAAGACAGCTTTGAATAGTGAAATTGGTAAAATTGCAAAATCTCTA	720
AZ65A1      AAGGGTATTGTCATCAAGACAGCTTTGAATAGTGAAATTGGTAAAATTGCAAAATCTCTA	720
ENA1        AAGGGTATTGTCATCAAGACAGCTTTGAATAGTGAAATTGGTAAAATTGCAAAATCTCTA	720
AZ118A1     AAGGGTATTGTCATCAAGACAGCTTTGAACAGTGAAATTGGTAAAATTGCAAAATCTCTA	720
AZ123A1     AAGGGTATTGTCATCAAGACAGCTTTGAATAGTGAAATTGGTAAAATTGCAAAATCTCTA	720
            ***************************** ******************************

AZ65B1      CAAGGTGATTCAGGTCTCATTTCCCGCGATCCTAGTAAGTCGTGGTTACAGAATACATGG	780
AZ123A2     CAAGGTGATTCAGGTCTCATTTCCCGCGATCCTAGTAAGTCGTGGTTACAGAATACATGG	780
AZ65A4      CAAGGTGATTCAGGTCTCATTTCCCGCGATCCTAGTAAGTCGTGGTTACAGAATACATGG	780
ENA5        CAAGGTGATTCAGGTCTCATTTCCCGCGATCCTAGTAAGTCGTGGTTACAGAATACATGG	780
AZ118A2     CAAGGTGATTCAGGTCTCATTTCCCGCGATCCTAGTAAGTCGTGGTTACAGAATACATGG	780
ENA2        CAAGGTGATTCAGGTCTCATTTCCCGCGATCCTAGTAAGTCGTGGTTACAGAATACATGG	780
AZ65A2      CAAGGTGATTCAGGTCTCATTTCCCGCGATCCTAGTAAGTCGTGGTTACAGAATACATGG	780
AZ65A3      CAAGGTGATTCAGGTCTCATTTCCCGCGATCCTAGTAAGTCGTGGTTACAGAATACATGG	780
AZ118B1     CAAGGTGATTCAGGTCTCATTTCCCGCGATCCTAGTAAGTCGTGGTTACAGAATACATGG	780
AZ65A1      CAAGGTGATTCAGGTCTCATTTCCCGCGATCCTAGTAAGTCGTGGTTACAGAATACATGG	780
ENA1        CAAGGTGATTCAGGTCTCATTTCCCGCGATCCTAGTAAGTCGTGGTTACAGAATACATGG	780
AZ118A1     CAAGGTGATTCAGGTCTCATTTCCCGCGATCCTAGTAAGTCGTGGTTACAGAATACATGG	780
AZ123A1     CAAGGTGATTCAGGTCTCATTTCCCGCGATCCTAGTAAGTCGTGGTTACAGAATACATGG	780
            ************************************************************

AZ65B1      ATATCTACAAAGAAAGTTACTGGTGCATTTTTAGGTACAAATGTCGGTACGCCCCTGCAC	840
AZ123A2     ATATCTACAAAGAAAGTTACTGGTGCATTTTTAGGTACAAATGTCGGTACGCCCCTGCAC	840
AZ65A4      ATATCTACAAAGAAAGTTACTGGTGCATTTTTAGGTACAAATGTCGGTACGCCCCTGCAC	840
ENA5        ATATCTACAAAGAAAGTTACTGGTGCATTTTTAGGTACAAATGTCGGTACGCCCCTGCAC	840
AZ118A2     ATATCTACAAAGAAAGTTACTGGTGCATTTTTAGGTACAAATGTCGGTACGCCCCTGCAC	840
ENA2        ATATCTACAAAGAAAGTTACTGGTGCATTTTTAGGTACAAATGTCGGTACGCCCCTGCAC	840
AZ65A2      ATATCTACAAAGAAAGTTACTGGTGCATTTTTAGGTACAAATGTCGGTACGCCCCTGCAC	840
AZ65A3      ATATCTACAAAGAAAGTTACTGGTGCATTTTTAGGTACAAATGTCGGTACGCCCCTGCAC	840
AZ118B1     ATATCTACAAAGAAAGTTACTGGTGCATTTTTAGGTACAAATGTCGGTACGCCCCTGCAC	840
AZ65A1      ATATCTACAAAGAAAGTTACTGGTGCATTTTTAGGTACAAATGTCGGTACGCCCCTGCAC	840
ENA1        ATATCTACAAAGAAAGTTACTGGTGCATTTTTAGGTACAAATGTCGGTACGCCCCTGCAC	840
AZ118A1     ATATCTACAAAGAAAGTTACTGGTGCATTTTTAGGTACAAATGTCGGTACGCCCCTGCAC	840
AZ123A1     ATATCTACAAAGAAAGTTACTGGTGCATTTTTAGGTACAAATGTCGGTACGCCCCTGCAC	840
            ************************************************************

AZ65B1      AGGAAACTGTCTAAGTTGGCGGTATTGCTTTTCTGGATAGCTGTATTTGTTTGCTATAAT	900
AZ123A2     AGGAAACTGTCTAAGTTGGCGGTATTGCTTTTCTGGATTGCCGTACTT-TTTGCTATCAT	899
AZ65A4      AGGAAACTGTCTAAGTTGGCGGTATTGCTTTTCTGGATTGCCGTACTT-TTTGCTATCAT	899
ENA5        AGGAAACTGTCTAAGTTGGCGGTATTGCTTTTCTGGATTGCCGTACTT-TTTGCTATCAT	899
AZ118A2     AGGAAACTGTCTAAGTTGGCGGTATTGCTTTTCTGGATTGCCGTACTT-TTTGCTATCAT	899
ENA2        AGGAAACTGTCTAAGTTGGCGGTATTGCTTTTCTGGATTGCCGTACTT-TTTGCTATCAT	899
AZ65A2      AGGAAACTGTCTAAGTTGGCGGTATTGCTTTTCTGGATTGCCGTACTT-TTTGCTATCAT	899
AZ65A3      AGGAAACTGTCTAAGTTGGCGGTATTGCTTTTCTGGATTGCCGTACTT-TTTGCTATCAT	899
AZ118B1     AGGAAACTGTCTAAGTTGGCGGTATTGCTTTTCTGGATTGCCGTACTT-TTTGCTATCAT	899
AZ65A1      AGGAAACTGTCTAAGTTGGCGGTATTGCTTTTCTGGATTGCCGTACTT-TTTGCTATCAT	899
ENA1        AGGAAACTGTCTAAGTTGGCGGTATTGCTTTTCTGGATTGCCGTACTT-TTTGCTATCAT	899
AZ118A1     AGGAAACTGTCTAAGTTGGCGGTATTGCTTTTCTGGATTGCCGTACTT-TTTGCTATCAT	899
AZ123A1     AGGAAACTGTCTAAGTTGGCGGTATTGCTTTTCTGGATTGCCGTACTT-TTTGCTATCAT	899
            ************************************** ** *** ** ******** **
AZ65B1      TGTTATGGCCTCTCAAAAGTTCAGTGTAGATAAGGGTATAGCTATCTATGCAATTTGCGT	960
AZ123A2     CGTCATGGCCTCTCAAAAGTTTGACGTAGATAAAAGGGTAGCTATCTATGCCATTTGTGT	959
AZ65A4      CGTCATGGCCTCTCAAAAGTTTGACGTAGATAAAAGGGTAGCTATCTATGCCATTTGTGT	959
ENA5        CGTCATGGCCTCTCAAAAGTTTGACGTAGATAAAAGGGTAGCTATCTATGCCATTTGTGT	959
AZ118A2     CGTCATGGCCTCTCAAAAGTTTGACGTAGATAAAAGGGTAGCTATCTATGCCATTTGTGT	959
ENA2        CGTCATGGCCTCTCAAAAGTTTGACGTAGATAAAAGGGTAGCTATCTATGCCATTTGTGT	959
AZ65A2      CGTCATGGCCTCTCAAAAGTTTGACGTAGATAAAAGGGTAGCTATCTATGCCATTTGTGT	959
AZ65A3      CGTCATGGCCTCTCAAAAGTTTGACGTAGATAAAAGGGTAGCTATCTATGCCATTTGTGT	959
AZ118B1     CGTCATGGCCTCTCAAAAGTTTGACGTAGATAAAAGGGTAGCTATCTATGCCATTTGTGT	959
AZ65A1      CGTCATGGCCTCTCAAAAGTTTGACGTAGATAAAAGGGTAGCTATCTATGCCATTTGTGT	959
ENA1        CGTCATGGCCTCTCAAAAGTTTGACGTAGATAAAAGGGTAGCTATCTATGCCATTTGTGT	959
AZ118A1     CGTCATGGCCTCTCAAAAGTTTGACGTAGATAAAAGGGTAGCTATCTATGCCATTTGTGT	959
AZ123A1     CGTCATGGCCTCTCAAAAGTTTGACGTAGATAAAAGGGTAGCTATCTATGCCATTTGTGT	959
             ** *****************    ********  *  ************* ***** **

AZ65B1      TGCCCTATCTATGATTCCCTCTTCATTAGTCGTTGTCTTGACTATTACTATGTCTGTAGG	1020
AZ123A2     GGCCCTATCCATGATTCCCTCTTCATTAGTCGTTGTCTTGACCATCACCATGTCTGTTGG	1019
AZ65A4      GGCCCTATCCATGATTCCCTCTTCATTAGTCGTTGTCTTGACCATCACCATGTCTGTTGG	1019
ENA5        GGCCCTATCCATGATTCCCTCTTCATTAGTCGTTGTCTTGACCATCACCATGTCTGTTGG	1019
AZ118A2     GGCCCTATCCATGATTCCCTCTTCATTAGTCGTTGTCTTGACCATCACCATGTCTGTTGG	1019
ENA2        GGCCCTATCCATGATTCCCTCTTCATTAGTCGTTGTCTTGACCATCACCATGTCTGTTGG	1019
AZ65A2      GGCCCTATCCATGATTCCCTCTTCATTAGTCGTTGTCTTGACCATCACCATGTCTGTTGG	1019
AZ65A3      GGCCCTATCCATGATTCCCTCTTCATTAGTCGTTGTCTTGACCATCACCATGTCTGTTGG	1019
AZ118B1     GGCCCTATCCATGATTCCCTCTTCATTAGTCGTTGTCTTGACCATCACCATGTCTGTTGG	1019
AZ65A1      GGCCCTATCCATGATTCCCTCTTCATTAGTCGTTGTCTTGACCATCACCATGTCTGTTGG	1019
ENA1        GGCCCTATCCATGATTCCCTCTTCATTAGTCGTTGTCTTGACCATCACCATGTCTGTTGG	1019
AZ118A1     GGCCCTATCCATGATTCCCTCTTCATTAGTCGTTGTCTTGACCATCACCATGTCTGTTGG	1019
AZ123A1     GGCCCTATCCATGATTCCCTCTTCATTAGTCGTTGTCTTGACCATCACCATGTCTGTTGG	1019
             ******** ******************************** ** ** ******** **

AZ65B1      AGCTGCCGTTATGGTTTCTAGACATGTTATTGTAAGAAAATTAGATTCTTTGGAGGCCCT	1080
AZ123A2     GGCTGCTGTTATGGTTTCTAGAAACGTTATTGTAAGAAAATTAGATTCTCTGGAAGCTTT	1079
AZ65A4      GGCTGCTGTTATGGTTTCTAGAAACGTTATTGTAAGAAAATTAGATTCTCTGGAAGCTTT	1079
ENA5        GGCTGCTGTTATGGTTTCTAGAAACGTTATTGTAAGAAAATTAGATTCTCTGGAAGCTTT	1079
AZ118A2     GGCTGCTGTTATGGTTTCTAGAAACGTTATTGTAAGAAAATTAGATTCTCTGGAAGCTTT	1079
ENA2        GGCTGCTGTTATGGTTTCTAGAAACGTTATTGTAAGAAAATTAGATTCTCTGGAAGCTTT	1079
AZ65A2      GGCTGCTGTTATGGTTTCTAGAAACGTTATTGTAAGAAAATTAGATTCTCTGGAAGCTTT	1079
AZ65A3      GGCTGCTGTTATGGTTTCTAGAAACGTTATTGTAAGAAAATTAGATTCTCTGGAAGCTTT	1079
AZ118B1     GGCTGCTGTTATGGTTTCTAGAAACGTTATTGTAAGAAAATTAGATTCTCTGGAAGCTTT	1079
AZ65A1      GGCTGCTGTTATGGTTTCTAGAAACGTTATTGTAAGAAAATTAGATTCTCTGGAAGCTTT	1079
ENA1        GGCTGCTGTTATGGTTTCTAGAAACGTTATTGTAAGAAAATTAGATTCTCTGGAAGCTTT	1079
AZ118A1     GGCTGCTGTTATGGTTTCTAGAAACGTTATTGTAAGAAAATTAGATTCTCTGGAAGCTTT	1079
AZ123A1     GGCTGCTGTTATGGTTTCTAGAAACGTTATTGTAAGAAAATTAGATTCTCTGGAAGCTTT	1079
             ***** *************** * ************************ **** **  *

AZ65B1      AGGTGCAGTTAACGACATTTGTTCTGACAAGACCGGTACTCTTACACAGGGAAAAATGTT	1140
AZ123A2     AGGTGCCGTTAACGATATCTGTTCTGACAAGACCGGTACTCTTACACAGGGTAAAATGTT	1139
AZ65A4      AGGTGCCGTTAACGATATCTGTTCTGACAAGACCGGTACTCTTACACAGGGTAAAATGTT	1139
ENA5        AGGTGCCGTTAACGATATCTGTTCTGACAAGACCGGTACTCTTACACAGGGTAAAATGTT	1139
AZ118A2     AGGTGCCGTTAACGATATCTGTTCTGACAAGACCGGTACTCTTACACAGGGTAAAATGTT	1139
ENA2        AGGTGCCGTTAACGATATCTGTTCTGACAAGACCGGTACTCTTACACAGGGTAAAATGTT	1139
AZ65A2      AGGTGCCGTTAACGATATCTGTTCTGACAAGACCGGTACTCTTACACAGGGTAAAATGTT	1139
AZ65A3      AGGTGCCGTTAACGATATCTGTTCTGACAAGACCGGTACTCTTACACAGGGTAAAATGTT	1139
AZ118B1     AGGTGCCGTTAACGATATCTGTTCTGACAAGACCGGTACTCTTACACAGGGTAAAATGTT	1139
AZ65A1      AGGTGCCGTTAACGATATCTGTTCTGACAAGACCGGTACTCTTACACAGGGTAAAATGTT	1139
ENA1        AGGTGCCGTTAACGATATCTGTTCTGACAAGACCGGTACTCTTACACAGGGTAAAATGTT	1139
AZ118A1     AGGTGCCGTTAACGATATCTGTTCTGACAAGACCGGTACTCTTACACAGGGTAAAATGTT	1139
AZ123A1     AGGTGCCGTTAACGATATCTGTTCTGACAAGACCGGTACTCTTACACAGGGTAAAATGTT	1139
            ****** ******** ** ******************************** ********
AZ65B1      AGCAAGGCAGATTTGGATCCCTCGCTTTGGTACTATAACTATCTCGAATTCTGATGACCC	1200
AZ123A2     AGCGAGGCAAATTTGGATCCCTCGCTTTGGTACCATAACTATCTCGAATTCTGATGACCC	1199
AZ65A4      AGCGAGGCAAATTTGGATCCCTCGCTTTGGTACCATAACTATCTCGAATTCTGATGACCC	1199
ENA5        AGCGAGGCAAATTTGGATCCCTCGCTTTGGTACCATAACTATCTCGAATTCTGATGACCC	1199
AZ118A2     AGCGAGGCAAATTTGGATCCCTCGCTTTGGTACCATAACTATCTCGAATTCTGATGACCC	1199
ENA2        AGCGAGGCAAATTTGGATCCCTCGCTTTGGTACCATAACTATCTCGAATTCTGATGACCC	1199
AZ65A2      AGCGAGGCAAATTTGGATCCCTCGCTTTGGTACCATAACTATCTCGAATTCTGATGACCC	1199
AZ65A3      AGCGAGGCAAATTTGGATCCCTCGCTTTGGTACCATAACTATCTCGAATTCTGATGACCC	1199
AZ118B1     AGCGAGGCAAATTTGGATCCCTCGCTTTGGTACCATAACTATCTCGAATTCTGATGACCC	1199
AZ65A1      AGCGAGGCAAATTTGGATCCCTCGCTTTGGTACCATAACTATCTCGAATTCTGATGACCC	1199
ENA1        AGCGAGGCAAATTTGGATCCCTCGCTTTGGTACCATAACTATCTCGAATTCTGATGACCC	1199
AZ118A1     AGCGAGGCAAATTTGGATCCCTCGCTTTGGTACCATAACTATCTCGAATTCTGATGACCC	1199
AZ123A1     AGCGAGGCAAATTTGGATCCCTCGCTTTGGTACCATAACTATCTCGAATTCTGATGACCC	1199
            *** ***** *********************** **************************

AZ65B1      CTTTAATCCCAATGAGGGCAACGTGAGTTTGATTCCAAGGTTTTCACCTTACGAGTACTC	1260
AZ123A2     CTTTAATCCCAATGAGGGCAACGTGAGTTTGATTCCAAGGTTTTCACCTTACGAATATTC	1259
AZ65A4      CTTTAATCCCAATGAGGGCAACGTGAGTTTGATTCCAAGGTTTTCACCTTACGAATATTC	1259
ENA5        CTTTAATCCCAATGAGGGCAACGTGAGTTTGATTCCAAGGTTTTCACCTTACGAATATTC	1259
AZ118A2     CTTTAATCCCAATGAGGGCAACGTGAGTTTGATTCCAAGGTTTTCACCTTACGAATATTC	1259
ENA2        CTTTAATCCCAATGAGGGCAACGTGAGTTTGATTCCAAGGTTTTCACCTTACGAATATTC	1259
AZ65A2      CTTTAATCCCAATGAGGGCAACGTGAGTTTGATTCCAAGGTTTTCACCTTACGAATATTC	1259
AZ65A3      CTTTAATCCCAATGAGGGCAACGTGAGTTTGATTCCAAGGTTTTCACCTTACGAATATTC	1259
AZ118B1     CTTTAATCCCAATGAGGGCAACGTGAGTTTGATTCCAAGGTTTTCACCTTACGAATATTC	1259
AZ65A1      CTTTAATCCCAATGAGGGCAACGTGAGTTTGATTCCAAGGTTTTCACCTTACGAATATTC	1259
ENA1        CTTTAATCCCAATGAGGGCAACGTGAGTTTGATTCCAAGGTTTTCACCTTACGAATATTC	1259
AZ118A1     CTTTAATCCCAATGAGGGCAACGTGAGTTTGATTCCAAGGTTTTCACCTTACGAATATTC	1259
AZ123A1     CTTTAATCCCAATGAGGGCAACGTGAGTTTGATTCCAAGGTTTTCACCTTACGAATATTC	1259
            ****************************************************** ** **

AZ65B1      CCACAATGAGGATGGTGACGTCGGTATTCTCCAAAACTTTCAAGGATCGCCTATACGAAA	1320
AZ123A2     TCATAATGAGGATGGTGACGTTGGTATTCTCCAGAA-TTTCAAGGATCGCCTATACGAAA	1318
AZ65A4      TCATAATGAGGATGGTGACGTTGGTATTCTCCAGAA-TTTCAAGGATCGCCTATACGAAA	1318
ENA5        TCATAATGAGGATGGTGACGTTGGTATTCTCCAGAA-TTTCAAGGATCGCCTATACGAAA	1318
AZ118A2     TCATAATGAGGATGGTGACGTTGGTATTCTCCAGAACTTTCAAGGATCGCCTATACGAAA	1319
ENA2        TCATAATGAGGATGGTGACGTTGGTATTCTCCAGAA-TTTCAAGGATCGCCTATACGAAA	1318
AZ65A2      TCATAATGAGGATGGTGACGTTGGTATTCTCCAGAA-TTTCAAGGATCGCCTATACGAAA	1318
AZ65A3      TCATAATGAGGATGGTGACGTTGGTATTCTCCAGAA-TTTCAAGGATCGCCTATACGAAA	1318
AZ118B1     TCATAATGAGGATGGTGACGTTGGTATTCTCCAGAA-TTTCAAGGATCGCCTATACGAAA	1318
AZ65A1      TCATAATGAGGATGGTGACGTTGGTATTCTCCAGAA-TTTCAAGGATCGCCTATACGAAA	1318
ENA1        TCATAATGAGGATGGTGACGTTGGTATTCTCCAGAA-TTTCAAGGATCGCCTATACGAAA	1318
AZ118A1     TCATAATGAGGATGGTGACGTTGGTATTCTCCAGAA-TTTCAAGGATCGCCTATACGAAA	1318
AZ123A1     TCATAATGAGGATGGTGACGTTGGTATTCTCCAGAA-TTTCAAGGATCGCCTATACGAAA	1318
             ** ***************** *********** ** ***********************

AZ65B1      AGGATTTACCAGAAGATATTGACATGGATCTATTTCAAAAATGGCTCGAAACCGCCACTT	1380
AZ123A2     AAGATTTACCAGAAGATATTGACATGGATCTATTTCAAAAATGGCTCGAAACCGCCACTT	1378
AZ65A4      AAGATTTACCAGAAGATATTGACATGGATCTATTTCAAAAATGGCTCGAAACCGCCACTT	1378
ENA5        AAGATTTACCAGAAGATATTGACATGGATCTATTTCAAAAATGGCTCGAAACCGCCACTT	1378
AZ118A2     AAGATTTACCAGAAGATATTGACATGGATCTATTTCAAAAATGGCTCGAAACCGCCACTT	1379
ENA2        AAGATTTACCAGAAGATATTGACATGGATCTATTTCAAAAATGGCTCGAAACCGCCACTT	1378
AZ65A2      AAGATTTACCAGAAGATATTGACATGGATCTATTTCAAAAATGGCTCGAAACCGCCACTT	1378
AZ65A3      AAGATTTACCAGAAGATATTGACATGGATCTATTTCAAAAATGGCTCGAAACCGCCACTT	1378
AZ118B1     AAGATTTACCAGAAGATATTGACATGGATCTATTTCAAAAATGGCTCGAAACCGCCACTT	1378
AZ65A1      AAGATTTACCAGAAGATATTGACATGGATCTATTTCAAAAATGGCTCGAAACCGCCACTT	1378
ENA1        AAGATTTACCAGAAGATATTGACATGGATCTATTTCAAAAATGGCTCGAAACCGCCACTT	1378
AZ118A1     AAGATTTACCAGAAGATATTGACATGGATCTATTTCAAAAATGGCTCGAAACCGCCACTT	1378
AZ123A1     AAGATTTACCAGAAGATATTGACATGGATCTATTTCAAAAATGGCTCGAAACCGCCACTT	1378
            * **********************************************************
AZ65B1      TGGCTAACATTGCTACTGTTTTCAAAGATGACGCAACTGACTGTTGGAAAGCTCACGGTG	1440
AZ123A2     TGGCTAACATTGCTACTGTTTTCAAAGATGACGCAACTGACTGTTGGAAAGCTCATGGTG	1438
AZ65A4      TGGCTAACATTGCTACTGTTTTCAAAGATGACGCAACTGACTGTTGGAAAGCTCATGGTG	1438
ENA5        TGGCTAACATTGCTACTGTTTTCAAAGATGACGCAACTGACTGTTGGAAAGCTCATGGTG	1438
AZ118A2     TGGCTAACATTGCTACTGTTTTCAAAGATGACGCAACTGACTGTTGGAAAGCTCATGGTG	1439
ENA2        TGGCTAACATTGCTACTGTTTTCAAAGATGACGCAACTGACTGTTGGAAAGCTCATGGTG	1438
AZ65A2      TGGCTAACATTGCTACTGTTTTCAAAGATGACGCAACTGACTGTTGGAAAGCTCATGGTG	1438
AZ65A3      TGGCTAACATTGCTACTGTTTTCAAAGATGACGCAACTGACTGTTGGAAAGCTCATGGTG	1438
AZ118B1     TGGCTAACATTGCTACTGTTTTCAAAGATGACGCAACTGACTGTTGGAAAGCTCATGGTG	1438
AZ65A1      TGGCTAACATTGCTACTGTTTTCAAAGATGACGCAACTGACTGTTGGAAAGCTCATGGTG	1438
ENA1        TGGCTAACATTGCTACTGTTTTCAAAGATGACGCAACTGACTGTTGGAAAGCTCATGGTG	1438
AZ118A1     TGGCTAACATTGCTACTGTTTTCAAAGATGACGCAACTGACTGTTGGAAAGCTCATGGTG	1438
AZ123A1     TGGCTAACATTGCTACTGTTTTCAAAGATGACGCAACTGACTGTTGGAAAGCTCATGGTG	1438
            ******************************************************* ****

AZ65B1      ATCCAACAGAAATTGCCATTCAGGTATTTGCTACCAAGATGGATTTACCTCGCAATGCCC	1500
AZ123A2     ACCCAACAGAAATTGCGATTCAAGTGTTTGCTACTAAGATGGACTTGCCTCGCAATGCCC	1498
AZ65A4      ACCCAACAGAAATTGCGATTCAAGTGTTTGCTACTAAGATGGACTTGCCTCGCAATGCCC	1498
ENA5        ACCCAACAGAAATTGCGATTCAAGTGTTTGCTACTAAGATGGACTTGCCTCACAATGCCC	1498
AZ118A2     ACCCAACAGAAATTGCGATTCAAGTGTTTGCTACTAAGATGGACTTGCCTCGCAATGCCC	1499
ENA2        ACCCAACAGAAATTGCGATTCAAGTGTTTGCTACTAAGATGGACTTGCCTCACAATGCCC	1498
AZ65A2      ACCCAACAGAAATTGCGATTCAAGTGTTTGCTACTAAGATGGACTTGCCTCGCAATGCCC	1498
AZ65A3      ACCCAACAGAAATTGCGATTCAAGTGTTTGCTACTAAGATGGACTTGCCTCGCAATGCCC	1498
AZ118B1     ACCCAACAGAAATTGCGATTCAAGTGTTTGCTACTAAGATGGACTTGCCTCGCAATGCCC	1498
AZ65A1      ACCCAACAGAAATTGCGATTCAAGTGTTTGCTACTAAGATGGACTTGCCTCGCAATGCCC	1498
ENA1        ACCCAACAGAAATTGCGATTCAAGTGTTTGCTACTAAGATGGACTTGCCTCACAATGCCC	1498
AZ118A1     ACCCAACAGAAATTGCGATTCAAGTGTTTGCTACTAAGATGGACTTGCCTCGCAATGCCC	1498
AZ123A1     ACCCAACAGAAATTGCGATTCAAGTGTTTGCTACTAAGATGGACTTGCCTCGCAATGCCC	1498
            * ************** ***** ** ******** ******** ** **** ********

AZ65B1      TTACCGGTGAGAAATCGAATAATCAAAGTAATGAAAGCGATAAATCTTCTCTACTGGGTC	1560
AZ123A2     TTACCGGTGAGAAATCGACTAATCAAAGTAATGAGAATGATCAATCCTCTCTTTCACAAC	1558
AZ65A4      TTACCGGTGAGAAATCGACTAATCAAAGTAATGAGAATGATCAATCCTCTCTTTCACAAC	1558
ENA5        TTACCGGTGAGAAATCGACTAATCAAAGTAATGAGAATGACCAATCCTCTCTTTCACAAC	1558
AZ118A2     TTACCGGTGAGAAATCGACTAATCAAAGTAATGAGAATGATCAATCCTCTCTTTCACAAC	1559
ENA2        TTACCGGTGAGAAATCGACTAATCAAAGTAATGAGAATGACCAATCCTCTCTTTCACAAC	1558
AZ65A2      TTACCGGTGAGAAATCGACTAATCAAAGTAATGAGAATGATCAATCCTCTCTTTCACAAC	1558
AZ65A3      TTACCGGTGAGAAATCGACTAATCAAAGTAATGAGAATGATCAATCCTCTCTTTCACAAC	1558
AZ118B1     TTACCGGTGAGAAATCGACTAATCAAAGTAATGAGAATGATCAATCCTCTCTTTCACAAC	1558
AZ65A1      TTACCGGTGAGAAATCGACTAATCAAAGTAATGAGAATGATCAATCCTCTCTTTCACAAC	1558
ENA1        TTACCGGTGAGAAATCGACTAATCAAAGTAATGAGAATGACCAATCCTCTCTTTCACAAC	1558
AZ118A1     TTACCGGTGAGAAATCGACTAATCAAAGTAATGAGAATGATCAATCCTCTCTTTCACAAC	1558
AZ123A1     TTACCGGTGAGAAATCGACTAATCAAAGTAATGAGAATGATCAATCCTCTCTTTCACAAC	1558
            ****************** *************** *  **  **** *****       *

AZ65B1      ACAATGAGAAACCCAGTAATGCCCAATTCCAGCATATCGCTGAATTTCCATTTGATTCGA	1620
AZ123A2     ACAATGAGAAGCCTGGCAGTGCACAATTCGAACATATTGCTGAATTCCCATTCGACTCAA	1618
AZ65A4      ACAATGAGAAGCCTGGCAGTGCACAATTCGAACATATTGCTGAATTCCCATTCGACTCAA	1618
ENA5        ACAATGAGAAGCCTGGCAGTGCACAATTCGAACATATTGCTGAATTCCCATTCGACTCAA	1618
AZ118A2     ACAATGAGAAGCCTGGCAGTGCACAATTCGAACATATTGCTGAATTCCCATTCGACTCAA	1619
ENA2        ACAATGAGAAGCCTGGCAGTGCACAATTCGAACATATTGCTGAATTCCCATTCGACTCAA	1618
AZ65A2      ACAATGAGAAGCCTGGCAGTGCACAATTCGAACATATTGCTGAATTCCCATTCGACTCAA	1618
AZ65A3      ACAATGAGAAGCCTGGCAGTGCACAATTCGAACATATTGCTGAATTCCCATTCGACTCAA	1618
AZ118B1     ACAATGAGAAGCCTGGCAGTGCACAATTCGAACATATTGCTGAATTCCCATTCGACTCAA	1618
AZ65A1      ACAATGAGAAGCCTGGCAGTGCACAATTCGAACATATTGCTGAATTCCCATTCGACTCAA	1618
ENA1        ACAATGAGAAGCCTGGCAGTGCACAATTCGAACATATTGCTGAATTCCCATTCGACTCAA	1618
AZ118A1     ACAATGAGAAGCCTGGCAGTGCACAATTCGAACATATTGCTGAATTCCCATTCGACTCAA	1618
AZ123A1     ACAATGAGAAGCCTGGCAGTGCACAATTCGAACATATTGCTGAATTCCCATTCGACTCAA	1618
            ********** **  * * *** ****** * ***** ******** ***** ** ** *
AZ65B1      CAGTGAAGAGAATGTCATCTGTATACTACAACAATCACAACGAAACATACAACGTTTATG	1680
AZ123A2     CTGTGAAGCGAATGTCATCTGTCTACTACAACAATCACAACGAAACATATAATATTTATG	1678
AZ65A4      CTGTGAAGCGAATGTCATCTGTCTACTACAACAATCACAACGAAACATATAATATTTATG	1678
ENA5        CTGTGAAGCGAATGTCATCTGTCTACTACAACAATCACAACGAAACATATAATATTTATG	1678
AZ118A2     CTGTGAAGCGAATGTCATCTGTCTACTACAACAATCACAACGAAACATATAATATTTATG	1679
ENA2        CTGTGAAGCGAATGTCATCTGTCTACTACAACAATCACAACGAAACATATAATATTTATG	1678
AZ65A2      CTGTGAAGCGAATGTCATCTGTCTACTACAACAATCACAACGAAACATATAATATTTATG	1678
AZ65A3      CTGTGAAGCGAATGTCATCTGTCTACTACAACAATCACAACGAAACATATAATATTTATG	1678
AZ118B1     CTGTGAAGCGAATGTCATCTGTCTACTACAACAATCACAACGAAACATATAATATTTATG	1678
AZ65A1      CTGTGAAGCGAATGTCATCTGTCTACTACAACAATCACAACGAAACATATAATATTTATG	1678
ENA1        CTGTGAAGCGAATGTCATCTGTCTACTACAACAATCACAACGAAACATATAATATTTATG	1678
AZ118A1     CTGTGAAGCGAATGTCATCTGTCTACTACAACAATCACAACGAAACATATAATATTTATG	1678
AZ123A1     CTGTGAAGCGAATGTCATCTGTCTACTACAACAATCACAACGAAACATATAATATTTATG	1678
            * ****** ************* ************************** **  ******

AZ65B1      GAAAAGGTGCTTTTGAAAGCATTATCAGCTGTTGCAGCTCTTGGTATGGCAAGGATGGTA	1740
AZ123A2     GCAAGGGTGCTTTCGAAAGCATCATCAGTTGTTGCAGTTCTTGGTATGGTAAGGATGGTG	1738
AZ65A4      GCAAGGGTGCTTTCGAAAGCATCATCAGTTGTTGCAGTTCTTGGTATGGTAAGGATGGTG	1738
ENA5        GCAAGGGTGCTTTCGAAAGCATCATCAGTTGTTGCAGTTCTTGGTATGGTAAGGATGGTG	1738
AZ118A2     GCAAGGGTGCTTTCGAAAGCATCATCAGTTGTTGCAGTTCTTGGTATGGTAAGGATGGTG	1739
ENA2        GCAAGGGTGCTTTCGAAAGCATCATCAGTTGTTGCAGTTCTTGGTATGGTAAGGATGGTG	1738
AZ65A2      GCAAGGGTGCTTTCGAAAGCATCATCAGTTGTTGCAGTTCTTGGTATGGTAAGGATGGTG	1738
AZ65A3      GCAAGGGTGCTTTCGAAAGCATCATCAGTTGTTGCAGTTCTTGGTATGGTAAGGATGGTG	1738
AZ118B1     GCAAGGGTGCTTTCGAAAGCATCATCAGTTGTTGCAGTTCTTGGTATGGTAAGGATGGTG	1738
AZ65A1      GCAAGGGTGCTTTCGAAAGCATCATCAGTTGTTGCAGTTCTTGGTATGGTAAGGATGGTG	1738
ENA1        GCAAGGGTGCTTTCGAAAGCATCATCAGTTGTTGCAGTTCTTGGTATGGTAAGGATGGTG	1738
AZ118A1     GCAAGGGTGCTTTCGAAAGCATCATCAGTTGTTGCAGTTCTTGGTATGGTAAGGATGGTG	1738
AZ123A1     GCAAGGGTGCTTTCGAAAGCATCATCAGTTGTTGCAGTTCTTGGTATGGTAAGGATGGTG	1738
            * ** ******** ******** ***** ******** *********** *********

AZ65B1      CAAAAATCACGCCTTTGACCGATTGTGATGTCGAAACTATAAGGAAAAATGTTTACAGTC	1800
AZ123A2     TAAAAATCACACCATTGACCGATTGTGATGTCGAAACGATAAGGAAAAATGTTTACAGTC	1798
AZ65A4      TAAAAATCACACCATTGACCGATTGTGATGTCGAAACGATAAGGAAAAATGTTTACAGTC	1798
ENA5        TAAAAATCACACCATTGACCGATTGTGATGTCGAAACGATAAGGAAAAATGTTTACAGTC	1798
AZ118A2     TAAAAATCACACCATTGACCGATTGTGATGTCGAAACGATAAGGAAAAATGTTTACAGTC	1799
ENA2        TAAAAATCACACCATTGACCGATTGTGATGTCGAAACGATAAGGAAAAATGTTTACAGTC	1798
AZ65A2      TAAAAATCACACCATTGACCGATTGTGATGTCGAAACGATAAGGAAAAATGTTTACAGTC	1798
AZ65A3      TAAAAATCACACCATTGACCGATTGTGATGTCGAAACGATAAGGAAAAATGTTTACAGTC	1798
AZ118B1     TAAAAATCACACCATTGACCGATTGTGATGTCGAAACGATAAGGAAAAATGTTTACAGTC	1798
AZ65A1      TAAAAATCACACCATTGACCGATTGTGATGTCGAAACGATAAGGAAAAATGTTTACAGTC	1798
ENA1        TAAAAATCACACCATTGACCGATTGTGATGTCGAAACGATAAGGAAAAATGTTTACAGTC	1798
AZ118A1     TAAAAATCACACCATTGACCGATTGTGATGTCGAAACGATAAGGAAAAATGTTTACAGTC	1798
AZ123A1     TAAAAATCACACCATTGACCGATTGTGATGTCGAAACGATAAGGAAAAATGTTTACAGTC	1798
             ********* ** *********************** **********************

AZ65B1      TATCAAATGAAGGTTTAAGAGTCTTGGGGTTTGCTTCCAAATCTTTTACCAAGGATGAAC	1860
AZ123A2     TATCAAATGAGGGTTTAAGAGTCTTGGGTTTTGCCTCCAAATCTTTCACTAAAGATCAAG	1858
AZ65A4      TATCAAATGAGGGTTTAAGAGTCTTGGGTTTTGCCTCCAAATCTTTCACTAAAGATCAAG	1858
ENA5        TATCAAATGAGGGTTTAAGAGTCTTGGGTTTTGCCTCCAAATCTTTCACTAAAGATCAAG	1858
AZ118A2     TATCAAATGAGGGTTTAAGAGTCTTGGGTTTTGCCTCCAAATCTTTCACTAAAGATCAAG	1859
ENA2        TATCAAATGAGGGTTTAAGAGTCTTGGGTTTTGCCTCCAAATCTTTCACTAAAGATCAAG	1858
AZ65A2      TATCAAATGAGGGTTTAAGAGTCTTGGGTTTTGCCTCCAAATCTTTCACTAAAGATCAAG	1858
AZ65A3      TATCAAATGAGGGTTTAAGAGTCTTGGGTTTTGCCTCCAAATCTTTCACTAAAGATCAAG	1858
AZ118B1     TATCAAATGAGGGTTTAAGAGTCTTGGGTTTTGCCTCCAAATCTTTCACTAAAGATCAAG	1858
AZ65A1      TATCAAATGAGGGTTTAAGAGTCTTGGGTTTTGCCTCCAAATCTTTCACTAAAGATCAAG	1858
ENA1        TATCAAATGAGGGTTTAAGAGTCTTGGGTTTTGCCTCCAAATCTTTCACTAAAGATCAAG	1858
AZ118A1     TATCAAATGAGGGTTTAAGAGTCTTGGGTTTTGCCTCCAAATCTTTCACTAAAGATCAAG	1858
AZ123A1     TATCAAATGAGGGTTTAAGAGTCTTGGGTTTTGCCTCCAAATCTTTCACTAAAGATCAAG	1858
            ********** ***************** ***** *********** ** ** *** **
AZ65B1      AAAGTGACGAGCAATTGAAGAATAATACTTCGAACAGGGCTGCCGCAGAAAGCGATTTAG	1920
AZ123A2     TGAATGACGATCAATTGAAAAACATTACTTCAAACAGGGCCACCGCAGAAAGTGATTTAG	1918
AZ65A4      TGAATGACGATCAATTGAAAAACATTACTTCAAACAGGGCCACCGCAGAAAGTGATTTAG	1918
ENA5        TGAATGACGATCAATTGAAAAACATTACTTCAAACAGGGCCACCGCAGAAAGTGATTTAG	1918
AZ118A2     TGAATGACGATCAATTGAAAAACATTACTTCAAACAGGGCCACCGCAGAAAGTGATTTAG	1919
ENA2        TGAATGACGATCAATTGAAAAACATTACTTCAAACAGGGCCACCGCAGAAAGTGATTTAG	1918
AZ65A2      TGAATGACGATCAATTGAAAAACATTACTTCAAACAGGGCCACCGCAGAAAGTGATTTAG	1918
AZ65A3      TGAATGACGATCAATTGAAAAACATTACTTCAAACAGGGCCACCGCAGAAAGTGATTTAG	1918
AZ118B1     TGAATGACGATCAATTGAAAAACATTACTTCAAACAGGGCCACCGCAGAAAGTGATTTAG	1918
AZ65A1      TGAATGACGATCAATTGAAAAACATTACTTCAAACAGGGCCACCGCAGAAAGTGATTTAG	1918
ENA1        TGAATGACGATCAATTGAAAAACATTACTTCAAACAGGGCCACCGCAGAAAGTGATTTAG	1918
AZ118A1     TGAATGACGATCAATTGAAAAACATTACTTCAAACAGGGCCACCGCAGAAAGTGATTTAG	1918
AZ123A1     TGAATGACGATCAATTGAAAAACATTACTTCAAACAGGGCCACCGCAGAAAGTGATTTAG	1918
              * ****** ******** ** * ****** ********  ********** *******

AZ65B1      TTTTCCTAGGTTTGATTGGTATTTACGATCCACCTAGAAACGAGACTGCCGGTGCAGTCA	1980
AZ123A2     TTTTCCTAGGGTTGATTGGTATTTACGATCCACCCAGAAATGAGACTGCCGGTGCAGTCA	1978
AZ65A4      TTTTCCTAGGGTTGATTGGTATTTACGATCCACCCAGAAATGAGACTGCCGGTGCAGTCA	1978
ENA5        TTTTCCTAGGGTTGATTGGTATTTACGATCCACCCAGAAATGAGACTGCCGGTGCAGTCA	1978
AZ118A2     TTTTCCTAGGGTTGATTGGTATTTACGATCCACCCAGAAATGAGACTGCCGGTGCAGTCA	1979
ENA2        TTTTCCTAGGGTTGATTGGTATTTACGATCCACCCAGAAATGAGACTGCCGGTGCAGTCA	1978
AZ65A2      TTTTCCTAGGGTTGATTGGTATTTACGATCCACCCAGAAATGAGACTGCCGGTGCAGTCA	1978
AZ65A3      TTTTCCTAGGGTTGATTGGTATTTACGATCCACCCAGAAATGAGACTGCCGGTGCAGTCA	1978
AZ118B1     TTTTCCTAGGGTTGATTGGTATTTACGATCCACCCAGAAATGAGACTGCCGGTGCAGTCA	1978
AZ65A1      TTTTCCTAGGGTTGATTGGTATTTACGATCCACCCAGAAATGAGACTGCCGGTGCAGTCA	1978
ENA1        TTTTCCTAGGGTTGATTGGTATTTACGATCCACCCAGAAATGAGACTGCCGGTGCAGTCA	1978
AZ118A1     TTTTCCTAGGGTTGATTGGTATTTACGATCCACCCAGAAATGAGACTGCCGGTGCAGTCA	1978
AZ123A1     TTTTCCTAGGGTTGATTGGTATTTACGATCCACCCAGAAATGAGACTGCCGGTGCAGTCA	1978
            ********** *********************** ***** *******************

AZ65B1      AGAAGTTTCATCAGGCCGGTATTAACGTCCATATGTTAACTGGGGACTTTGTGGGTACAG	2040
AZ123A2     AGAAGTTTCACCAAGCTGGTATTAACGTTCATATGTTAACTGGGGACTTTGTGGGTACAG	2038
AZ65A4      AGAAGTTTCACCAAGCTGGTATTAACGTTCATATGTTAACTGGGGACTTTGTGGGTACAG	2038
ENA5        AGAAGTTTCACCAAGCTGGTATTAACGTTCATATGTTAACTGGGGACTTTGTGGGTACAG	2038
AZ118A2     AGAAGTTTCACCAAGCTGGTATTAACGTTCATATGTTAACTGGGGACTTTGTGGGTACAG	2039
ENA2        AGAAGTTTCACCAAGCTGGTATTAACGTTCATATGTTAACTGGGGACTTTGTGGGTACAG	2038
AZ65A2      AGAAGTTTCACCAAGCTGGTATTAACGTTCATATGTTAACTGGGGACTTTGTGGGTACAG	2038
AZ65A3      AGAAGTTTCACCAAGCTGGTATTAACGTTCATATGTTAACTGGGGACTTTGTGGGTACAG	2038
AZ118B1     AGAAGTTTCACCAAGCTGGTATTAACGTTCATATGTTAACTGGGGACTTTGTGGGTACAG	2038
AZ65A1      AGAAGTTTCACCAAGCTGGTATTAACGTTCATATGTTAACTGGGGACTTTGTGGGTACAG	2038
ENA1        AGAAGTTTCACCAAGCTGGTATTAACGTTCATATGTTAACTGGGGACTTTGTGGGTACAG	2038
AZ118A1     AGAAGTTTCACCAAGCTGGTATTAACGTTCATATGTTAACTGGGGACTTTGTGGGTACAG	2038
AZ123A1     AGAAGTTTCACCAAGCTGGTATTAACGTTCATATGTTAACTGGGGACTTTGTGGGTACAG	2038
            ********** ** ** *********** *******************************

AZ65B1      CAAAGGCTATTGCTCAAGAAGTTGGTATCTTACCCACCAATTTGTACCATTACTCCCAAG	2100
AZ123A2     CAAAGGCTATCGCTCAGGAGGTTGGCATCTTACCCACCAATTTGTACCATTACTCCCAAG	2098
AZ65A4      CAAAGGCTATCGCTCAGGAGGTTGGTATCTTACCCACCAATTTGTACCATTACTCCCAAG	2098
ENA5        CAAAGGCTATCGCTCAGGAGGTTGGCATCTTACCCACCAATTTGTACCATTACTCCCAAG	2098
AZ118A2     CAAAGGCTATCGCTCAGGAGGTTGGCATCTTACCCACCAATTTGTACCATTACTCCCAAG	2099
ENA2        CAAAGGCTATCGCTCAGGAGGTTGGCATCTTACCCACCAATTTGTACCATTACTCCCAAG	2098
AZ65A2      CAAAGGCTATCGCTCAGGAGGTTGGCATCTTACCCACCAATTTGTACCATTACTCCCAAG	2098
AZ65A3      CAAAGGCTATCGCTCAGGAGGTTGGCATCTTACCCACCAATTTGTACCATTACTCCCAAG	2098
AZ118B1     CAAAGGCTATCGCTCAGGAGGTTGGCATCTTACCCACCAATTTGTACCATTACTCCCAAG	2098
AZ65A1      CAAAGGCTATCGCTCAGGAGGTTGGCATCTTACCCACCAATTTGTACCATTACTCCCAAG	2098
ENA1        CAAAGGCTATCGCTCAGGAGGTTGGCATCTTACCCACCAATTTGTACCATTACTCCCAAG	2098
AZ118A1     CAAAGGCTATCGCTCAGGAGGTTGGCATCTTACCCACCAATTTGTACCATTACTCCCAAG	2098
AZ123A1     CAAAGGCTATCGCTCAGGAGGTTGGCATCTTACCCACCAATTTGTACCATTACTCCCAAG	2098
            ********** ***** ** ***** **********************************
AZ65B1      AGATTGTTGACAGTATGGTCATGACCGGATCTCAGTTTGACGGACTAAGTGAGGAGGAAG	2160
AZ123A2     AGATTGTTGACAGTATGGTCATGACCGGATCTCAGTTTGACGGACTAAGTGAGGAGGAAG	2158
AZ65A4      AGATTGTTGACAGTATGGTCATGACCGGATCTCAGTTTGACGGACTAAGTGAGGAGGAAG	2158
ENA5        AGATTGTTGACAGTATGGTCATGACCGGATCTCAGTTTGACGGACTAAGTGAGGAGGAAG	2158
AZ118A2     AGATTGTTGACAGTATGGTCATGACCGGATCTCAGTTTGACGGACTAAGTGAGGAGGAAG	2159
ENA2        AGATTGTTGACAGTATGGTCATGACCGGATCTCAGTTTGACGGACTAAGTGAGGAGGAAG	2158
AZ65A2      AGATTGTTGACAGTATGGTCATGACCGGATCTCAGTTTGACGGACTAAGTGAGGAGGAAG	2158
AZ65A3      AGATTGTTGACAGTATGGTCATGACCGGATCTCAGTTTGACGGACTAAGTGAGGAGGAAG	2158
AZ118B1     AGATTGTTGACAGTATGGTCATGACCGGATCTCAGTTTGACGGACTAAGTGAGGAGGAAG	2158
AZ65A1      AGATTGTTGACAGTATGGTCATGACCGGATCTCAGTTTGACGGACTAAGTGAGGAGGAAG	2158
ENA1        AGATTGTTGACAGTATGGTCATGACCGGATCTCAGTTTGACGGACTAAGTGAGGAGGAAG	2158
AZ118A1     AGATTGTTGACAGTATGGTCATGACCGGATCTCAGTTTGACGGACTAAGTGAGGAGGAAG	2158
AZ123A1     AGATTGTTGACAGTATGGTCATGACCGGATCTCAGTTTGACGGACTAAGTGAGGAGGAAG	2158
            ************************************************************

AZ65B1      TGGACGATTTGCCCGTCTTACCTTTAGTTATTGCCCGTTGCTCTCCGCAGACTAAGGTCA	2220
AZ123A2     TGGACGATTTGCCCGTCTTACCTTTAGTTATTGCACGTTGCTCTCCGCAGACTAAGGTGA	2218
AZ65A4      TGGACGATTTGCCCGTCTTACCTTTAGTTATTGCACGTTGCTCTCCGCAGACTAAGGTGA	2218
ENA5        TGGACGATTTGCCCGTCTTACCTTTAGTTATTGCACGTTGCTCTCCGCAGACTAAGGTGA	2218
AZ118A2     TGGACGATTTGCCCGTCTTACCTTTAGTTATTGCACGTTGCTCTCCGCAGACTAAGGTGA	2219
ENA2        TGGACGATTTGCCCGTCTTACCTTTAGTTATTGCACGTTGCTCTCCGCAGACTAAGGTGA	2218
AZ65A2      TGGACGATTTGCCCGTCTTACCTTTAGTTATTGCACGTTGCTCTCCGCAGACTAAGGTGA	2218
AZ65A3      TGGACGATTTGCCCGTCTTACCTTTAGTTATTGCACGTTGCTCTCCGCAGACTAAGGTGA	2218
AZ118B1     TGGACGATTTGCCCGTCTTACCTTTAGTTATTGCACGTTGCTCTCCGCAGACTAAGGTGA	2218
AZ65A1      TGGACGATTTGCCCGTCTTACCTTTAGTTATTGCACGTTGCTCTCCGCAGACTAAGGTGA	2218
ENA1        TGGACGATTTGCCCGTCTTACCTTTAGTTATTGCACGTTGCTCTCCGCAGACTAAGGTGA	2218
AZ118A1     TGGACGATTTGCCCGTCTTACCTTTAGTTATTGCACGTTGCTCTCCGCAGACTAAGGTGA	2218
AZ123A1     TGGACGATTTGCCCGTCTTACCTTTAGTTATTGCACGTTGCTCTCCGCAGACTAAGGTGA	2218
            ********************************** *********************** *

AZ65B1      GAATGATCGAAGCTTTACACCGTAGGAAGAAGTTCTGCGCAATGACAGGTGACGGTGTCA	2280
AZ123A2     GAATGATCGAAGCTTTACACCGTAGGAAGAAGTTCTGCGCAATGACAGGTGACGGTGTTA	2278
AZ65A4      GAATGATCGAAGCTTTACACCGTAGGAAAAAGTTCTGCGCAATGACGGGTGATGGTGTTA	2278
ENA5        GAATGATCGAAGCTTTACACCGTAGGAAGAAGTTCTGCGCAATGACAGGTGACGGTGTTA	2278
AZ118A2     GAATGATCGAAGCTTTACACCGTAGGAAGAAGTTCTGCGCAATGACAGGTGACGGTGTTA	2279
ENA2        GAATGATCGAAGCTTTACACCGTAGGAAGAAGTTCTGCGCAATGACAGGTGACGGTGTTA	2278
AZ65A2      GAATGATCGAAGCTTTACACCGTAGGAAGAAGTTCTGCGCAATGACAGGTGACGGTGTTA	2278
AZ65A3      GAATGATCGAAGCTTTACACCGTAGGAAGAAGTTCTGCGCAATGACAGGTGACGGTGTTA	2278
AZ118B1     GAATGATCGAAGCTTTACACCGTAGGAAGAAGTTCTGCGCAATGACAGGTGACGGTGTTA	2278
AZ65A1      GAATGATCGAAGCTTTACACCGTAGGAAGAAGTTCTGCGCAATGACAGGTGACGGTGTTA	2278
ENA1        GAATGATCGAAGCTTTACACCGTAGGAAGAAGTTCTGCACAATGACAGGTGACGGTGTTA	2278
AZ118A1     GAATGATCGAAGCTTTACACCGTAGGAAGAAGTTCTGCGCAATGACAGGTGACGGTGTTA	2278
AZ123A1     GAATGATCGAAGCTTTACACCGTAGGAAGAAGTTCTGCGCAATGACAGGTGACGGTGTTA	2278
            **************************** ********* ******* ***** ***** *

AZ65B1      ACGATTCTCCATCTCTAAAAATGGCCAATGTTGGTATTGCAATGGGTATTAATGGTTCAG	2340
AZ123A2     ATGATTCTCCATCTCTAAAAATGGCCAATGTTGGTATTGCAATGGGTATTAATGGTTCAG	2338
AZ65A4      ACGATTCTCCATCTCTAAAAATGGCCAATGTTGGTATTGCAATGGGTATTAATGGTTCAG	2338
ENA5        ACGATTCTCCATCTCTAAAAATGGCCAATGTTGGTATTGCAATGGGTATTAATGGTTCAG	2338
AZ118A2     ATGATTCTCCATCTCTAAAAATGGCCAATGTTGGTATTGCAATGGGTATTAATGGTTCAG	2339
ENA2        ACGATTCTCCATCTCTAAAAATGGCCAATGTTGGTATTGCAATGGGTATTAATGGTTCAG	2338
AZ65A2      ACGATTCTCCATCTCTAAAAATGGCCAATGTTGGTATTGCAATGGGTATTAATGGTTCAG	2338
AZ65A3      ACGATTCTCCATCTCTAAAAATGGCCAATGTTGGTATTGCAATGGGTATTAATGGTTCAG	2338
AZ118B1     ATGATTCTCCATCTCTAAAAATGGCCAATGTTGGTATTGCAATGGGTATTAATGGTTCAG	2338
AZ65A1      ACGATTCTCCATCTCTAAAAATGGCCAATGTTGGTATTGCAATGGGTATTAATGGTTCAG	2338
ENA1        ACGATTCTCCATCTCTAAAAATGGCCAATGTTGGTATTGCAATGGGTATTAATGGTTCAG	2338
AZ118A1     ATGATTCTCCATCTCTAAAAATGGCCAATGTTGGTATTGCAATGGGTATTAATGGTTCAG	2338
AZ123A1     ATGATTCTCCATCTCTAAAAATGGCCAATGTTGGTATTGCAATGGGTATTAATGGTTCAG	2338
            * **********************************************************
AZ65B1      ATGTTTCCAAAGAAGCGTCTGATATTGTTCTAAGCGATGACAATTTTGCTTCTATTTTGA	2400
AZ123A2     ATGTTTCCAAAGAAGCGTCTGATATTGTTCTAAGCGATGACAATTTTGCTTCTATTTTGA	2398
AZ65A4      ATGTTTCCAAAGAAGCGTCTGATATTGTTCTAAGCGATGACAATTTTGCTTCTATTTTGA	2398
ENA5        ATGTTTCCAAAGAAGCGTCTGATATTGTTCTAAGCGATGACAATTTTGCTTCTATTTTGA	2398
AZ118A2     ATGTTTCCAAAGAAGCGTCTGATATTGTTCTAAGCGATGACAATTTTGCTTCTATTTTGA	2399
ENA2        ATGTTTCCAAAGAAGCGTCTGATATTGTTCTAAGCGATGACAATTTTGCTTCTATTTTGA	2398
AZ65A2      ATGTTTCCAAAGAAGCGTCTGATATTGTTCTAAGCGATGACAATTTTGCTTCTATTTTGA	2398
AZ65A3      ATGTTTCCAAAGAAGCGTCTGATATTGTTCTAAGCGATGACAATTTTGCTTCTATTTTGA	2398
AZ118B1     ATGTTTCCAAAGAAGCGTCTGATATTGTTCTAAGCGATGACAATTTTGCTTCTATTTTGA	2398
AZ65A1      ATGTTTCCAAAGAAGCGTCTGATATTGTTCTAAGCGATGACAATTTTGCTTCTATTTTGA	2398
ENA1        ATGTTTCCAAAGAAGCGTCTGATATTGTTCTAAGCGATGACAATTTTGCTTCTATTTTGA	2398
AZ118A1     ATGTTTCCAAAGAAGCGTCTGATATTGTTCTAAGCGATGACAATTTTGCTTCTATTTTGA	2398
AZ123A1     ATGTTTCCAAAGAAGCGTCTGATATTGTTCTAAGCGATGACAATTTTGCTTCTATTTTGA	2398
            ************************************************************

AZ65B1      ATGCTGTCGAAGAAGGTCGTAGGATGACGGATAACATTCAGAAGTTTGTCCTACAATTAT	2460
AZ123A2     ATGCTGTCGAAGAAGGTCGTAGGATGACGGATAACATTCAGAAGTTTGTCCTACAATTAT	2458
AZ65A4      ATGCTGTCGAAGAAGGTCGTAGGATGACGGATAACATTCAGAAGTTTGTCCTACAATTAT	2458
ENA5        ATGCTGTCGAAGAAGGTCGTAGGATGACGGATAACATTCAGAAGTTTGTCCTACAATTAT	2458
AZ118A2     ATGCTGTCGAAGAAGGTCGTAGGATGACGGATAACATTCAGAAGTTTGTCCTACAATTAT	2459
ENA2        ATGCTGTCGAAGAAGGTCGTAGGATGACGGATAACATTCAGAAGTTTGTCCTACAATTAT	2458
AZ65A2      ATGCTGTCGAAGAAGGTCGTAGGATGACGGATAACATTCAGAAGTTTGTCCTACAATTAT	2458
AZ65A3      ATGCTGTCGAAGAAGGTCGTAGGATGACGGATAACATTCAGAAGTTTGTCCTACAATTAT	2458
AZ118B1     ATGCTGTCGAAGAAGGTCGTAGGATGACGGATAACATTCAGAAGTTTGTCCTACAATTAT	2458
AZ65A1      ATGCTGTCGAAGAAGGTCGTAGGATGACGGATAACATTCAGAAGTTTGTCCTACAATTAT	2458
ENA1        ATGCTGTCGAAGAAGGTCGTAGGATGACGGATAACATTCAGAAGTTTGTCCTACAATTAT	2458
AZ118A1     ATGCTGTCGAAGAAGGTCGTAGGATGACGGATAACATTCAGAAGTTTGTCCTACAATTAT	2458
AZ123A1     ATGCTGTCGAAGAAGGTCGTAGGATGACGGATAACATTCAGAAGTTTGTCCTACAATTAT	2458
            ************************************************************

AZ65B1      TGGCAGAAAATGTTGCTCAGGCTTTGTATTTGATCATTGGTTTAGTATTCAGAGATGAGA	2520
AZ123A2     TGGCAGAAAATGTTGCTCAGGCTTTGTATTTGATCATTGGTTTAGTATTCAGAGATGAGA	2518
AZ65A4      TGGCAGAAAATGTTGCTCAGGCTTTGTATTTGATCATTGGTTTAGTATTCAGAGATGAGA	2518
ENA5        TGGCAGAAAATGTTGCTCAGGCTTTGTATTTGATCATTGGTTTAGTATTCAGAGATGAGA	2518
AZ118A2     TGGCAGAAAATGTTGCTCAGGCTTTGTATTTGATCATTGGTTTAGTATTCAGAGATGAGA	2519
ENA2        TGGCAGAAAATGTTGCTCAGGCTTTGTATTTGATCATTGGTTTAGTATTCAGAGATGAGA	2518
AZ65A2      TGGCAGAAAATGTTGCTCAGGCTTTGTATTTGATCATTGGTTTAGTATTCAGAGATGAGA	2518
AZ65A3      TGGCAGAAAATGTTGCTCAGGCTTTGTATTTGATCATTGGTTTAGTATTCAGAGATGAGA	2518
AZ118B1     TGGCAGAAAATGTTGCTCAGGCTTTGTATTTGATCATTGGTTTAGTATTCAGAGATGAGA	2518
AZ65A1      TGGCAGAAAATGTTGCTCAGGCTTTGTATTTGATCATTGGTTTAGTATTCAGAGATGAGA	2518
ENA1        TGGCAGAAAATGTTGCTCAGGCTTTGTATTTGATCATTGGTTTAGTATTCAGAGATGAGA	2518
AZ118A1     TGGCAGAAAATGTTGCTCAGGCTTTGTATTTGATCATTGGTTTAGTATTCAGAGATGAGA	2518
AZ123A1     TGGCAGAAAATGTTGCTCAGGCTTTGTATTTGATCATTGGTTTAGTATTCAGAGATGAGA	2518
            ************************************************************

AZ65B1      ACGGAAAATCAGTGTTTCCCTTATCGCCTGTAGAAGTTTTATGGATTATCGTTGTCACCT	2580
AZ123A2     ACGGAAAATCAGTATTTCCCTTATCACCAGTGGAAGTATTGTGGATTATTGTCGTCACCT	2578
AZ65A4      ACGGAAAATCAGTATTTCCCTTATCACCAGTGGAAGTATTGTGGATTATTGTCGTCACCT	2578
ENA5        ACGGAAAATCAGTATTTCCCTTATCACCAGTGGAAGTATTGTGGATTATTGTCGTCACCT	2578
AZ118A2     ACGGAAAATCAGTATTTCCCTTATCACCAGTGGAAGTATTGTGGATTATTGTCGTCACCT	2579
ENA2        ACGGAAAATCAGTATTTCCCTTATCACCAGTGGAAGTATTGTGGATTATTGTCGTCACCT	2578
AZ65A2      ACGGAAAATCAGTATTTCCCTTATCACCAGTGGAAGTATTGTGGATTATTGTCGTCACCT	2578
AZ65A3      ACGGAAAATCAGTATTTCCCTTATCACCAGTGGAAGTATTGTGGATTATTGTCGTCACCT	2578
AZ118B1     ACGGAAAATCAGTATTTCCCTTATCACCAGTGGAAGTATTGTGGATTATTGTCGTCACCT	2578
AZ65A1      ACGGAAAATCAGTATTTCCCTTATCACCAGTGGAAGTATTGTGGATTATTGTCGTCACCT	2578
ENA1        ACGGAAAATCAGTATTTCCCTTATCACCAGTGGAAGTATTGTGGATTATTGTCGTCACCT	2578
AZ118A1     ACGGAAAATCAGTATTTCCCTTATCACCAGTGGAAGTATTGTGGATTATTGTCGTCACCT	2578
AZ123A1     ACGGAAAATCAGTATTTCCCTTATCACCAGTGGAAGTATTGTGGATTATTGTCGTCACCT	2578
            ************* *********** ** ** ***** ** ******** ** *******
AZ65B1      CTTGTTTCCCTGCTATGGGGCTAGGTCTAGAAAAGGCTGCTCCAGATTTAATGGATAGAC	2640
AZ123A2     CTTGTTTTCCTGCTATGGGGCTAGGTCTAGAAAAGGCTGCTCCAGATTTGATGGATAGAC	2638
AZ65A4      CTTGTTTTCCTGCTATGGGGCTAGGTCTAGAAAAGGCTGCTCCAGATTTGATGGATAGAC	2638
ENA5        CTTGTTTTCCTGCTATGGGGCTAGGTCTAGAAAAGGCTGCTCCAGATTTGATGGATAGAC	2638
AZ118A2     CTTGTTTTCCTGCTATGGGGCTAGGTCTAGAAAAGGCTGCTCCAGATTTGATGGATAGAC	2639
ENA2        CTTGTTTTCCTGCTATGGGGCTAGGTCTAGAAAAGGCTGCTCCAGATTTGATGGATAGAC	2638
AZ65A2      CTTGTTTTCCTGCTATGGGGCTAGGTCTAGAAAAGGCTGCTCCAGATTTGATGGATAGAC	2638
AZ65A3      CTTGTTTTCCTGCTATGGGGCTAGGTCTAGAAAAGGCTGCTCCAGATTTGATGGATAGAC	2638
AZ118B1     CTTGTTTTCCTGCTATGGGGCTAGGTCTAGAAAAGGCTGCTCCAGATTTGATGGATAGAC	2638
AZ65A1      CTTGTTTTCCTGCTATGGGGCTAGGTCTAGAAAAGGCTGCTCCAGATTTGATGGATAGAC	2638
ENA1        CTTGTTTTCCTGCTATGGGGCTAGGTCTAGAAAAGGCTGCTCCAGATTTGATGGATAGAC	2638
AZ118A1     CTTGTTTTCCTGCTATGGGGCTAGGTCTAGAAAAGGCTGCTCCAGATTTGATGGATAGAC	2638
AZ123A1     CTTGTTTTCCTGCTATGGGGCTAGGTCTAGAAAAGGCTGCTCCAGATTTGATGGATAGAC	2638
            ******* ***************************************** **********

AZ65B1      CTCCTAACGATTCAGAAGTTGGTATTTTCACCTGGGAAGTTATTATCGATACATTTGCAT	2700
AZ123A2     CTCCTAATGATTCAGAGGTTGGTATTTTCACATGGGAGGTTATTATAGATACATTTGCAT	2698
AZ65A4      CTCCTAATGATTCAGAGGTTGGTATTTTCACATGGGAGGTTATTATAGATACATTTGCAT	2698
ENA5        CTCCTAATGATTCAGAGGTTGGTATTTTCACATGGGAGGTTATTATAGATACATTTGCAT	2698
AZ118A2     CTCCTAATGATTCAGAGGTTGGTATTTTCACATGGGAGGTTATTATAGATACATTTGCAT	2699
ENA2        CTCCTCATGATTCAGAGGTTGGTATTTTCACGTGGGAGGTTATTATAGATACATTTGCAT	2698
AZ65A2      CTCCTAATGATTCAGAGGTTGGTATTTTCACATGGGAGGTTATTATAGATACATTTGCAT	2698
AZ65A3      CTCCTAATGATTCAGAGGTTGGTATTTTCACATGGGAGGTTATTATAGATACATTTGCAT	2698
AZ118B1     CTCCTAATGATTCAGAGGTTGGTATTTTCACATGGGAGGTTATTATAGATACATTTGCAT	2698
AZ65A1      CTCCTAATGATTCAGAGGTTGGTATTTTCACATGGGAGGTTATTATAGATACATTTGCAT	2698
ENA1        CTCCTCATGATTCAGAGGTTGGTATTTTCACGTGGGAGGTTATTATAGATACATTTGCAT	2698
AZ118A1     CTCCTAATGATTCAGAGGTTGGTATTTTCACATGGGAGGTTATTATAGATACATTTGCAT	2698
AZ123A1     CTCCTAATGATTCAGAGGTTGGTATTTTCACATGGGAGGTTATTATAGATACATTTGCAT	2698
            ***** * ******** ************** ***** ******** *************

AZ65B1      ACGGGTTTTTGATGGCCGGTTCTTGTATGGCCTCCTTCACTGGGTCGCTCTACGGAATAA	2760
AZ123A2     ATGGGATTATAATGACAGGGTCCTGTATGGCTTCATTTACTGGATCACTGTATGGAATAA	2758
AZ65A4      ATGGGATTATAATGACAGGGTCCTGTATGGCTTCATTTACTGGATCACTGTATGGAATAA	2758
ENA5        ATGGGATTATAATGACAGGGTCCTGTATGGCTTCATTTACTGGATCACTGTATGGAATAA	2758
AZ118A2     ATGGGATTATAATGACAGGGTCCTGTATGGCTTCATTTACTGGATCACTGTATGGAATAA	2759
ENA2        ATGGGATTATAATGACAGGGTCCTGTATGGCTTCATTTACTGGATCACTGTATGGAATAA	2758
AZ65A2      ATGGGATTATAATGACAGGGTCCTGTATGGCTTCATTTACTGGATCACTGTATGGAATAA	2758
AZ65A3      ATGGGATTATAATGACAGGGTCCTGTATGGCTTCATTTACTGGATCACTGTATGGAATAA	2758
AZ118B1     ATGGGATTATAATGACAGGGTCCTGTATGGCTTCATTTACTGGATCACTGTATGGAATAA	2758
AZ65A1      ATGGGATTATAATGACAGGGTCCTGTATGGCTTCATTTACTGGATCACTGTATGGAATAA	2758
ENA1        ATGGGATTATAATGACAGGGTCCTGTATGGCTTCATTTACTGGATCACTGTATGGAATAA	2758
AZ118A1     ATGGGATTATAATGACAGGGTCCTGTATGGCTTCATTTACTGGATCACTGTATGGAATAA	2758
AZ123A1     ATGGGATTATAATGACAGGGTCCTGTATGGCTTCATTTACTGGATCACTGTATGGAATAA	2758
            * *** ** * *** * ** ** ******** ** ** ***** ** ** ** *******

AZ65B1      ATAATGGTGAATTGGGAGTTAATTGTGATAGATCTTATAATGGTAGCTGTCATGATGTTT	2820
AZ123A2     ATAGTGGTAGATTGGGGCACGATTGTGATGGCACCTATAACAGCAGTTGTCGTGATGTTT	2818
AZ65A4      ATAGTGGTAGATTGGGGCACGATTGTGATGGCACCTATAACAGCAGTTGTCGTGATGTTT	2818
ENA5        ATAGTGGTAGATTGGGGCACGATTGTGATGGCACCTATAACAGCAGTTGTCGTGATGTTT	2818
AZ118A2     ATAGTGGTAGATTGGGGCACGATTGTGATGGCACCTATAACAGCAGTTGTCGTGATGTTT	2819
ENA2        ATAGTGGTAGATTGGGGCACGATTGTGATGGCACCTATAACAGCAGTTGTCGTGATGTTT	2818
AZ65A2      ATAGTGGTAGATTGGGGCACGATTGTGATGGCACCTATAACAGCAGTTGTCGTGATGTTT	2818
AZ65A3      ATAGTGGTAGATTGGGGCACGATTGTGATGGCACCTATAACAGCAGTTGTCGTGATGTTT	2818
AZ118B1     ATAGTGGTAGATTGGGGCACGATTGTGATGGCACCTATAACAGCAGTTGTCGTGATGTTT	2818
AZ65A1      ATAGTGGTAGATTGGGGCACGATTGTGATGGCACCTATAACAGCAGTTGTCGTGATGTTT	2818
ENA1        ATAGTGGTAGATTGGGGCACGATTGTGATGGCACCTATAACAGCAGTTGTCGTGATGTTT	2818
AZ118A1     ATAGTGGTAGATTGGGGCACGATTGTGATGGCACCTATAACAGCAGTTGTCGTGATGTTT	2818
AZ123A1     ATAGTGGTAGATTGGGGCACGATTGTGATGGCACCTATAACAGCAGTTGTCGTGATGTTT	2818
            *** ****  ******     ******** *  * *****  * ** **** ********
AZ65B1      ATAGATCACGTTCTGCAGCCTTCGCAACTATGACATGGTGTGCTTTGATTTTGGCTTGGG	2880
AZ123A2     ATAGATCACGTTCTGCGGCTTTCGCAACTATGACGTGGTGCGCTTTGATTCTGGCTTGGG	2878
AZ65A4      ATAGATCACGTTCTGCGGCTTTCGCAACTATGACGTGGTGCGCTTTGATTCTGGCTTGGG	2878
ENA5        ATAGATCACGTTCTGCGGCTTTCGCAACTATGACGTGGTGCGCTTTGATTCTGGCTTGGG	2878
AZ118A2     ATAGATCACGTTCTGCGGCTTTCGCAACTATGACGTGGTGCGCTTTGATTCTGGCTTGGG	2879
ENA2        ATAGATCACGTTCTGCGGCTTTCGCAACCATGACGTGGTGCGCTTTGATTCTGGCTTGGG	2878
AZ65A2      ATAGATCACGTTCTGCGGCTTTCGCAACTATGACGTGGTGCGCTTTGATTCTGGCTTGGG	2878
AZ65A3      ATAGATCACGTTCTGCGGCTTTCGCAACTATGACGTGGTGCGCTTTGATTCTGGCTTGGG	2878
AZ118B1     ATAGATCACGTTCTGCGGCTTTCGCAACTATGACGTGGTGCGCTTTGATTCTGGCTTGGG	2878
AZ65A1      ATAGATCACGTTCTGCGGCTTTCGCAACTATGACGTGGTGCGCTTTGATTCTGGCTTGGG	2878
ENA1        ATAGATCACGTTCTGCGGCTTTCGCAACCATGACGTGGTGCGCTTTGATTCTGGCTTGGG	2878
AZ118A1     ATAGATCACGTTCTGCGGCTTTCGCAACTATGACGTGGTGCGCTTTGATTCTGGCTTGGG	2878
AZ123A1     ATAGATCACGTTCTGCGGCTTTCGCAACTATGACGTGGTGCGCTTTGATTCTGGCTTGGG	2878
            **************** ** ******** ***** ***** ********* *********

AZ65B1      AAGTGGTTGATATGAGAAGATCTTTTTTCAGAATGCATCCAGACACTGACAGCCCAGTTA	2940
AZ123A2     AAGTGGTTGACATGAGAAGATCTTTTTTCAGAATGCATCCAGACACTGACAGCCCAGTCA	2938
AZ65A4      AAGTGGTTGACATGAGAAGATCTTTTTTCAGAATGCATCCAGACACTGACAGCCCAGTCA	2938
ENA5        AAGTGGTTGACATGAGAAGATCTTTTTTCAGAATGCATCCAGACACTGACAGCCCAGTCA	2938
AZ118A2     AAGTGGTTGACATGAGAAGATCTTTTTTCAGAATGCATCCAGACACTGACAGCCCAGTCA	2939
ENA2        AAGTGGTTGACATGAGAAGATCCTTTTTCAGAATGCATCCAGACACTGACAGCCCAGTCA	2938
AZ65A2      AAGTGGTTGACATGAGAAGATCTTTTTTCAGAATGCATCCAGACACTGACAGCCCAGTCA	2938
AZ65A3      AAGTGGTTGACATGAGAAGATCTTTTTTCAGAATGCATCCAGACACTGACAGCCCAGTCA	2938
AZ118B1     AAGTGGTTGACATGAGAAGATCTTTTTTCAGAATGCATCCAGACACTGACAGCCCAGTCA	2938
AZ65A1      AAGTGGTTGACATGAGAAGATCTTTTTTCAGAATGCATCCAGACACTGACAGCCCAGTCA	2938
ENA1        AAGTGGTTGACATGAGAAGATCCTTTTTCAGAATGCATCCAGACACTGACAGCCCAGTCA	2938
AZ118A1     AAGTGGTTGACATGAGAAGATCTTTTTTCAGAATGCATCCAGACACTGACAGCCCAGTCA	2938
AZ123A1     AAGTGGTTGACATGAGAAGATCTTTTTTCAGAATGCATCCAGACACTGACAGCCCAGTCA	2938
            ********** *********** *********************************** *

AZ65B1      AGGAATTTTTTAAGAGTATTTGGGGAAACCAGTTTTTGTTTTGGTCAATCATTTTTGGAT	3000
AZ123A2     AGGAATTTTTCAGAAGTATTTGGGGAAACCAGTTTTTGTTCTGGTCAATCATTTTTGGAT	2998
AZ65A4      AGGAATTTTTCAGAAGTATTTGGGGAAACCAGTTTTTGTTCTGGTCAATCATTTTTGGAT	2998
ENA5        AGGAATTTTTCAGAAGTATTTGGGGAAACCAGTTTTTGTTCTGGTCAATCATTTTTGGAT	2998
AZ118A2     AGGAATTTTTCAGAAGTATTTGGGGAAACCAGTTTTTGTTCTGGTCAATCATTTTTGGAT	2999
ENA2        AGGAATTTTTCAGAAGTATTTGGGGAAACCAGTTTTTGTTCTGGTCAATCATTTTTGGAT	2998
AZ65A2      AGGAATTTTTCAGAAGTATTTGGGGAAACCAGTTTTTGTTCTGGTCAATCATTTTTGGAT	2998
AZ65A3      AGGAATTTTTCAGAAGTATTTGGGGAAACCAGTTTTTGTTCTGGTCAATCATTTTTGGAT	2998
AZ118B1     AGGAATTTTTCAGAAGTATTTGGGGAAACCAGTTTTTGTTCTGGTCAATCATTTTTGGAT	2998
AZ65A1      AGGAATTTTTCAGAAGTATTTGGGGAAACCAGTTTTTGTTCTGGTCAATCATTTTTGGAT	2998
ENA1        AGGAATTTTTCAGAAGCATTTGGGGAAACCAGTTTTTGTTCTGGTCAATCATTTTTGGAT	2998
AZ118A1     AGGAATTTTTCAGAAGTATTTGGGGAAACCAGTTTTTGTTCTGGTCAATCATTTTTGGAT	2998
AZ123A1     AGGAATTTTTCAGAAGTATTTGGGGAAACCAGTTTTTGTTCTGGTCAATCATTTTTGGAT	2998
            ********** *  ** *********************** *******************

AZ65B1      TTGTGTCTGCCTTCCCTGTCGTCTATATTCCGGTTATTAATGATAAAGTGTTTTTGCATA	3060
AZ123A2     TTGTGTCAGCCTTCCCCGTCGTCTATATTCCGGTTATTAATGATAAAGTGTTTTTGCATA	3058
AZ65A4      TTGTGTCAGCCTTCCCCGTCGTCTATATTCCGGTTATTAATGATAAAGTGTTTTTGCATA	3058
ENA5        TTGTGTCAGCCTTCCCCGTCGTCTATATTCCGGTTATTAATGATAAAGTGTTTTTGCATA	3058
AZ118A2     TTGTGTCAGCCTTCCCCGTCGTCTATATTCCGGTTATTAATGATAAAGTGTTTTTGCATA	3059
ENA2        TTGTGTCAGCCTTCCCCGTCGTCTATATTCCGGTTATTAATGATAAAGTGTTTTTGCATA	3058
AZ65A2      TTGTGTCAGCCTTCCCCGTCGTCTATATTCCGGTTATTAATGATAAAGTGTTTTTGCATA	3058
AZ65A3      TTGTGTCAGCCTTCCCCGTCGTCTATATTCCGGTTATTAATGATAAAGTGTTTTTGCATA	3058
AZ118B1     TTGTGTCAGCCTTCCCCGTCGTCTATATTCCGGTTATTAATGATAAAGTGTTTTTGCATA	3058
AZ65A1      TTGTGTCAGCCTTCCCCGTCGTCTATATTCCGGTTATTAATGATAAAGTGTTTTTGCATA	3058
ENA1        TTGTGTCAGCCTTCCCCGTCGTCTATATTCCGGTTATTAATGATAAAGTGTTTTTGCATA	3058
AZ118A1     TTGTGTCAGCCTTCCCCGTCGTCTATATTCCGGTTATTAATGATAAAGTGTTTTTGCATA	3058
AZ123A1     TTGTGTCAGCCTTCCCCGTCGTCTATATTCCGGTTATTAATGATAAAGTGTTTTTGCATA	3058
            ******* ******** *******************************************
AZ65B1      AACCAATTGGTCCTGAATGGGGTCTCGCCATTGCATTCACAATTGCATTCTGGATTGGTG	3120
AZ123A2     AACCAATTGGTGCTGAATGGGGTCTCGCCATTGCATTCACAATTGCATTCTGGATAGGTG	3118
AZ65A4      AACCAATTGGTGCTGAATGGGGTCTCGCCATTGCATTCACAATTGCATTCTGGGTAGGTG	3118
ENA5        AACCAATTGGTGCTGAATGGGGTCTCGCCATTGCATTCACAATTGCATTCTGGATAGGTG	3118
AZ118A2     AACCAATTGGTGCTGAATGGGGTCTCGCCATTGCATTCACAATTGCATTCTGGATAGGTG	3119
ENA2        AACCAATTGGTGCTGAATGGGGTCTCGCCATTGCATTCACAATTGCATTCTGGATAGGTG	3118
AZ65A2      AACCAATTGGTGCTGAATGGGGTCTCGCCATTGCATTCACAATTGCATTCTGGATAGGTG	3118
AZ65A3      AACCAATTGGTGCTGAATGGGGTCTCGCCATTGCATTCACAATTGCATTCTGGATAGGTG	3118
AZ118B1     AACCAATTGGTGCTGAATGGGGTCTCGCCATTGCATTCACAATTGCATTCTGGATAGGTG	3118
AZ65A1      AACCAATTGGTGCTGAATGGGGTCTCGCCATTGCATTCACAATTGCATTCTGGATAGGTG	3118
ENA1        AACCAATTGGTGCTGAATGGGGTCTCGCCATTGCATTCACAATTGCATTCTGGATAGGTG	3118
AZ118A1     AACCAATTGGTGCTGAATGGGGTCTCGCCATTGCATTCACAATTGCATTCTGGATAGGTG	3118
AZ123A1     AACCAATTGGTGCTGAATGGGGTCTCGCCATTGCATTCACAATTGCATTCTGGATAGGTG	3118
            *********** ***************************************** * ****

AZ65B1      CTGAACTTTACAAGTTAGGGAAGAGGCGCTACTTCAAAACTCAGAGAGCACACAACCCGG	3180
AZ123A2     CTGAACTTTACAAGTGTGGAAAGAGGCGCTATTTCAAAACTCAGAGAGCACACAACCCGG	3178
AZ65A4      CTGAACTTTACAAGTGTGGAAAGAGGCGCTATTTCAAAACTCAGAGAGCTCACAATTCGG	3178
ENA5        CTGAACTTTACAAGTGTGGAAAGAGGCGCTATTTCAAAACTCAGAGAGCTCACAATTCGG	3178
AZ118A2     CTGAACTTTACAAGTGTGGAAAGAGGCGCTATTTCAAAACTCAGAGAGCACACAACCCGG	3179
ENA2        CTGAACTTTACAAGTGTGGAAAGAGGCGCTATTTCAAAACTCAGAGAGCGCACAACCCGG	3178
AZ65A2      CTGAACTTTACAAGTGTGGAAAGAGGCGCTATTTCAAAACTCAGAGAGCGCACAACCCGG	3178
AZ65A3      CTGAACTTTACAAGTGTGGAAAGAGGCGCTATTTCAAAACTCAGAGAGCGCACAACCCGG	3178
AZ118B1     CTGAACTTTACAAGTGTGGAAAGAGGCGCTATTTCAAAACTCAGAGAGCGCACAACCCGG	3178
AZ65A1      CTGAACTTTACAAGTGTGGAAAGAGGCGCTATTTCAAAACTCAGAGAGCGCACAACCCGG	3178
ENA1        CTGAACTTTACAAGTGTGGAAAGAGGCGCTATTTCAAAACTCAGAGAGCGCACAACCCGG	3178
AZ118A1     CTGAACTTTACAAGTGTGGAAAGAGGCGCTATTTCAAAACTCAGAGAGCGCACAACCCGG	3178
AZ123A1     CTGAACTTTACAAGTGTGGAAAGAGGCGCTATTTCAAAACTCAGAGAGCGCACAACCCGG	3178
            ***************  ** *********** ***************** *****  ***

AZ65B1      AAAACGATTTGGAGAGTAA-CAATAAGCGCGATCCATTCGAAGCGTACAGTACTTCTACT	3239
AZ123A2     AAAACGATTTGGAGCGTAAGCAGTAAACACGATCCATTTGAAGCGTATAGCACATCCACA	3238
AZ65A4      AAAACGATTTGGAGCG-AAGCAGTAAACACGATCCATTTGAAGCGTATAGCACATCCACA	3237
ENA5        AAAACGATTTGGAGCG-AAGCAGTAAACACGATCCATTTGAAGCGTATAGCACATCCACA	3237
AZ118A2     AAAACGATTTGGAGAG-TAACAATAAGCGCGGTCCATTCGAAGCGTACAGTACTTCTACT	3238
ENA2        AGAATGATTTGGAGAG-TAACAATAAGCGCGATCCATTCGAAGCGTATAGTACTTCTACT	3237
AZ65A2      AGAATGATTTGGAGAG-TAACAATAAGCGCGATCCATTCGAAGCGTATAGTACTTCTACT	3237
AZ65A3      AGAATGATTTGGAGAG-TAACAATAAGCGCGATCCATTCGAAGCGTATAGTACTTCTACT	3237
AZ118B1     AGAATGATTTGGAGAG-TAACAATAAGCGCGATCCATTCGAAGCGTATAGTACTTCTACT	3237
AZ65A1      AGAATGATTTGGAGAG-TAACAATAAGCGCGATCCATTCGAAGCGTATAGTACTTCTACT	3237
ENA1        AGAATGATTTGGAGAG-TAACAATAAGCGCGATCCATTCGAAGCGTATAGTACTTCTACT	3237
AZ118A1     AGAATGATTTGGAGAG-TAACAATAAGCGCGATCCATTCGAAGCGTATAGTACTTCTACT	3237
AZ123A1     AGAATGATTTGGAGAG-TAACAATAAGCGCGATCCATTCGAAGCGTATAGTACTTCTACT	3237
            * ** ********* *  * ** *** * ** ****** ******** ** ** ** **

AZ65B1      ACAATCCATACA-GAAATTAATATTGGTATCAAACAATGA	3278
AZ123A2     ACCCTTCAAAGC-GAAATTAATATCAGTGTCAAGCATTAA	3277
AZ65A4      ACCCTTCAAAGC-GAAATTAATATCAGTGTCAAGCATTAA	3276
ENA5        ACCCTTCAAAGC-GAAATTAATATCAGTGTCAAGCATTAA	3276
AZ118A2     ACAATCCATAGCAGAAATTAATATTGGTATCAAACAATGA	3278
ENA2        ACAATCCATA-CAGAAGTTAATATTGGTATTAAACAATGA	3276
AZ65A2      ACAATCCATA-CAGAAGCTAATATTGGTATTAAACAATGA	3276
AZ65A3      ACAATCCATA-CAGAAGCTAATATTGGTATTAAACAATGA	3276
AZ118B1     ACAATCCATA-CAGAAGCTAATATTGGTATTAAACAATGA	3276
AZ65A1      ACAATCCATA-CAGAAGCTAATATTGGTATTAAACAATGA	3276
ENA1        ACAATCCATA-CAGAAGTTAATATTGGTATTAAACAATGA	3276
AZ118A1     ACAATCCATA-CAGAAGCTAATATTGGTATTAAACAATGA	3276
AZ123A1     ACAATCCATA-CAGAAGCTAATATTGGTATTAAACAATGA	3276
            **  * ** *   ***  ******  ** * ** ** * *
```
