## Supplemental Figure 2 for "Analysis of genome sequences of marine isolates of *Saccharomyces cerevisiae*"

**Supplementary Figure S2: Alignment of ENA protein sequences generated from DNA sequences with possible indels manually corrected**. Sequences are colour coded as follows: red = *ENA1*-like; green = *ENA2*-like; blue = *ENA5*-like. Variant residues are highlighted as follows: red = *ENA1*-like; yellow = *ENA1/ENA2*-like; green = *ENA2*-like; cyan = *ENA2/ENA5*-like; blue = *ENA5*-like. Unique variants or those which do not follow this pattern are highlighted magenta. It can be seen that the *ENA1*-like sequences have a unique N-terminal region (red) which changes to the *ENA2/ENA5*-like pattern (cyan) at a variable location around residues 80 to 107, after which all variants are non-systematic (magenta) until the C-terminal region, where the *ENA1*-like and *ENA2*-like sequences remain similar, but the *ENA5*-like sequences diverge after residues 1059 to 1065. Thus it appears that *ENA2*-like sequences are chimeric with *ENA5*-like N-terminus and *ENA1*-like C-terminus or, equivalently, *ENA2* and *ENA5*-like sequences are similar except that *ENA5*-like sequences have a unique C-terminus. AZ65B1 is seen to have the largest number of unique variants but is *ENA1*-like in the diagnostic N-terminal region.

```
AZ65B1       MGEGTTKENNNAEFNAYHTLTAEEAAEFIGTSLTEGLTQDEFVHRLKTVGENTLGDDTKI	60
ENA5         MSEGTVKENNNEEFNAYHTLTTEEAAEFIGTSLTEGLTQDESLRRLKAVGENTLGDDTKI	60
AZ65A4       MSEGTVKENNNEEFNAYHTLTTEEAAEFIGTSLTEGLTQDESLRRLKAVGENTLGDDTKI	60
AZ123A2      MSEGTVKENNNEEFNAYHTLTTEEAAEFIGTSLTEGLTQDESLRRLKAVGENTLGDDTKI	60
ENA2         MSEGTVKENNNEEFNAYHTLTTEEAAEFIGTSLTEGLTQDESLRRLKAVGENTLGDDTKI	60
AZ65A2       MSEGTVKENNNEEFNAYHTLTTEEAAEFIGTSLTEGLTQDESLRRLKAVGENTLGDDTKI	60
AZ65A3       MSEGTVKENNNEEFNAYHTLTTEEAAEFIGTSLTEGLTQDESLRRLKAVGENTLGDDTKI	60
AZ118B1      MSEGTVKENNNEEFNAYHTLTTEEAAEFIGTSLTEGLTQDESLRRLKAVGENTLGDDTKI	60
AZ118A2      MSEGTVKENNNEEFNAYHTLTTEEAAEFIGTSLTEGLTQDESLRRLKAVGENTLGDDTKI	60
ENA1         MGEGTTKENNNAEFNAYHTLTAEEAAEFIGTSLTEGLTQDEFVHRLKTVGENTLGDDTKI	60
AZ65A1       MGEGTTKENNNAEFNAYHTLTAEEAAEFIGTSLTEGLTQDEFVHRLKTVGENTLGDDTKI	60
AZ118A1      MGEGTTKENNNAEFNAYHTLTAEEAAEFIGTSLTEGLTQDEFVHRLKTVGENTLGDDTKI	60
AZ123A1      MGEGTTKENNNAEFNAYHTLTAEEAAEFIGTSLTEGLTQDEFVHRLKTVGENTLGDDTKI	60
             *.***.***** *********:******************* ::***:************

AZ65B1       DYKAMVLHQVCNAMIMVLLISMIISFAMHDWITGGVISFVVGVNVGIGLVQEYKATKTMN	120
ENA5         DYKAMVLHQVCNAMIMVLVISMAISFAVRDWITGGVISFVIAVNVLIGLVQEYKATKTMN	120
AZ65A4       DYKAMVLHQVCNAMIMVLVISMAISFAVRDWITGGVISFVIAVNVLIGLVQEYKATKTMN	120
AZ123A2      DYKAMVLHQVCNAMIMVLVISMAISFAVRDWITGGVISFVIAVNVLIGLVQEYKATKTMN	120
ENA2         DYKAMVLHQVCNAMIMVLVISMAISFAVRDWITGGVISFVIAVNVLIGLVQEYKATKTMN	120
AZ65A2       DYKAMVLHQVCNAMIMVLVISMAISFAVRDWITGGVISFVIAVNVLIGLVQEYKATKTMN	120
AZ65A3       DYKAMVLHQVCNAMIMVLVISMAISFAVRDWITGGVISFVIAVNVLIGLVQEYKATKTMN	120
AZ118B1      DYKAMVLHQVCNAMIMVLVISMAISFAVRDWITGGVISFVIAVNVLIGLVQEYKATKTMN	120
AZ118A2      DYKAMVLHQVCNAMIMVLVISMAISFAVRDWITGGVISFVIAVNVLIGLVQEYKATKTMN	120
ENA1         DYKAMVLHQVCNAMIMVLLISMIISFAMHDWITGGVISFVIAVNVLIGLVQEYKATKTMN	120
AZ65A1       DYKAMVLHQVCNAMIMVLLISMAISFAVRDWITGGVISFVIAVNVLIGLVQEYKATKTMN	120
AZ118A1      DYKAMVLHQVCNAMIMVLLISMIISFAMHDWITGGVISFVVGVNVGIGLVQEYKATKTMN	120
AZ123A1      DYKAMVLHQVCNAMIMVLLISMIISFAMHDWITGGVISFVVGVNVGIGLVQEYKATKTMN	120
             ******************:*** ****::***********:.*** **************
```

```
AZ65B1       SLKNLSSPNAHVIRNGKSETINSKDVVPGDICLVKVGDTIPADLRLIETKNFDTDESLLT	180
ENA5         SLKNLSSPNAHVIRNGKSETINSKDVVPGDICLVKVGDTIPADLRLIETKNFDTDESLLT	180
AZ65A4       SLKNLSSPNAHVIRNGKSETINSKDVVPGDICLVKVGDTIPADLRLIETKNFDTDESLLT	180
AZ123A2      SLKNLSSPNAHVIRNGKSETINSKDVVPGDICLVKVGDTIPADLRLIETKNFDTDESLLT	180
ENA2         SLKNLSSPNAHVIRNGKSETINSKDVVPGDICLVKVGDTIPADLRLIETKNFDTDESLLT	180
AZ65A2       SLKNLSSPNAHVIRNGKSETINSKDVVPGDICLVKVGDTIPADLRLIETKNFDTDESLLT	180
AZ65A3       SLKNLSSPNAHVIRNGKSETINSKDVVPGDICLVKVGDTIPADLRLIETKNFDTDESLLT	180
AZ118B1      SLKNLSSPNAHVIRNGKSETINSKDVVPGDICLVKVGDTIPADLRLIETKNFDTDESLLT	180
AZ118A2      SLKNLSSPNAHVIRNGKSETINSKDVVPGDICLVKVGDTIPADLRLIETKNFDTDESLLT	180
ENA1         SLKNLSSPNAHVIRNGKSETINSKDVVPGDICLVKVGDTIPADLRLIETKNFDTDESLLT	180
AZ65A1       SLKNLSSPNAHVIRNGKSETINSKDVVPGDICLVKVGDTIPADLRLIETKNFDTDESLLT	180
AZ118A1      SLKNLSSPNAHVIRNGKSETINSKDVVPGDICLVKVGDTIPADLRLIETKNFDTDESLLT	180
AZ123A1      SLKNLSSPNAHVIRNGKSETINSKDVVPGDICLVKVGDTIPADLRLIETKNFDTDESLLT	180
             ************************************************************

AZ65B1       GESLPVSKDADLVFGKEEETSVGDRLNLAFSSSTVVKGRAKGIVIKTALNSEIGKIAKSL	240
ENA5         GESLPVSKDANLVFGKEEETSVGDRLNLAFSSSAVVKGRAKGIVIKTALNSEIGKIAKSL	240
AZ65A4       GESLPVSKDANLVFGKEEETSVGDRLNLAFSSSTVVKGRAKGIVIKTALNSEIGKIAKSL	240
AZ123A2      GESLPVSKDANLVFGKEEETSVGDRLNLAFSSSTVVKGRAKGIVIKTALNSEIGKIAKSL	240
ENA2         GESLPVSKDANLVFGKEEETSVGDRLNLAFSSSAVVKGRAKGIVIKTALNSEIGKIAKSL	240
AZ65A2       GESLPVSKDANLVFGKEEETSVGDRLNLAFSSSTVVKGRAKGIVIKTALNSEIGKIAKSL	240
AZ65A3       GESLPVSKDANLVFGKEEETSVGDRLNLAFSSSTVVKGRAKGIVIKTALNSEIGKIAKSL	240
AZ118B1      GESLPVSKDANLVFGKEEETSVGDRLNLAFSSSTVVKGRAKGIVIKTALNSEIGKIAKSL	240
AZ118A2      GESLPVSKDANLVFGKEEETSVGDRLNLAFSSSTVVKGRAKGIVIKTALNSEIGKIAKSL	240
ENA1         GESLPVSKDANLVFGKEEETSVGDRLNLAFSSSAVVKGRAKGIVIKTALNSEIGKIAKSL	240
AZ65A1       GESLPVSKDANLVFGKEEETSVGDRLNLAFSSSTVVKGRAKGIVIKTALNSEIGKIAKSL	240
AZ118A1      GESLPVSKDADLVFGKEEETSVGDRLNLAFSSSTVVKGRAKGIVIKTALNSEIGKIAKSL	240
AZ123A1      GESLPVSKDANLVFGKEEETSVGDRLNLAFSSSTVVKGRAKGIVIKTALNSEIGKIAKSL	240
             **********:**********************:**************************

AZ65B1       QGDSGLISRDPSKSWLQNTWISTKKVTGAFLGTNVGTPLHRKLSKLAVLLFWIAVFFAII	300
ENA5         QGDSGLISRDPSKSWLQNTWISTKKVTGAFLGTNVGTPLHRKLSKLAVLLFWIAVLFAII	300
AZ65A4       QGDSGLISRDPSKSWLQNTWISTKKVTGAFLGTNVGTPLHRKLSKLAVLLFWIAVLFAII	300
AZ123A2      QGDSGLISRDPSKSWLQNTWISTKKVTGAFLGTNVGTPLHRKLSKLAVLLFWIAVLFAII	300
ENA2         QGDSGLISRDPSKSWLQNTWISTKKVTGAFLGTNVGTPLHRKLSKLAVLLFWIAVLFAII	300
AZ65A2       QGDSGLISRDPSKSWLQNTWISTKKVTGAFLGTNVGTPLHRKLSKLAVLLFWIAVLFAII	300
AZ65A3       QGDSGLISRDPSKSWLQNTWISTKKVTGAFLGTNVGTPLHRKLSKLAVLLFWIAVLFAII	300
AZ118B1      QGDSGLISRDPSKSWLQNTWISTKKVTGAFLGTNVGTPLHRKLSKLAVLLFWIAVLFAII	300
AZ118A2      QGDSGLISRDPSKSWLQNTWISTKKVTGAFLGTNVGTPLHRKLSKLAVLLFWIAVLFAII	300
ENA1         QGDSGLISRDPSKSWLQNTWISTKKVTGAFLGTNVGTPLHRKLSKLAVLLFWIAVLFAII	300
AZ65A1       QGDSGLISRDPSKSWLQNTWISTKKVTGAFLGTNVGTPLHRKLSKLAVLLFWIAVLFAII	300
AZ118A1      QGDSGLISRDPSKSWLQNTWISTKKVTGAFLGTNVGTPLHRKLSKLAVLLFWIAVLFAII	300
AZ123A1      QGDSGLISRDPSKSWLQNTWISTKKVTGAFLGTNVGTPLHRKLSKLAVLLFWIAVLFAII	300
             *******************************************************:****

AZ65B1       VMASQKFSVDKGIAIYAICVALSMIPSSLVVVLTITMSVGAAVMVSRHVIVRKLDSLEAL	360
ENA5         VMASQKFDVDKRVAIYAICVALSMIPSSLVVVLTITMSVGAAVMVSRNVIVRKLDSLEAL	360
AZ65A4       VMASQKFDVDKRVAIYAICVALSMIPSSLVVVLTITMSVGAAVMVSRNVIVRKLDSLEAL	360
AZ123A2      VMASQKFDVDKRVAIYAICVALSMIPSSLVVVLTITMSVGAAVMVSRNVIVRKLDSLEAL	360
ENA2         VMASQKFDVDKRVAIYAICVALSMIPSSLVVVLTITMSVGAAVMVSRNVIVRKLDSLEAL	360
AZ65A2       VMASQKFDVDKRVAIYAICVALSMIPSSLVVVLTITMSVGAAVMVSRNVIVRKLDSLEAL	360
AZ65A3       VMASQKFDVDKRVAIYAICVALSMIPSSLVVVLTITMSVGAAVMVSRNVIVRKLDSLEAL	360
AZ118B1      VMASQKFDVDKRVAIYAICVALSMIPSSLVVVLTITMSVGAAVMVSRNVIVRKLDSLEAL	360
AZ118A2      VMASQKFDVDKRVAIYAICVALSMIPSSLVVVLTITMSVGAAVMVSRNVIVRKLDSLEAL	360
ENA1         VMASQKFDVDKRVAIYAICVALSMIPSSLVVVLTITMSVGAAVMVSRNVIVRKLDSLEAL	360
AZ65A1       VMASQKFDVDKRVAIYAICVALSMIPSSLVVVLTITMSVGAAVMVSRNVIVRKLDSLEAL	360
AZ118A1      VMASQKFDVDKRVAIYAICVALSMIPSSLVVVLTITMSVGAAVMVSRNVIVRKLDSLEAL	360
AZ123A1      VMASQKFDVDKRVAIYAICVALSMIPSSLVVVLTITMSVGAAVMVSRNVIVRKLDSLEAL	360
             *******.*** :**********************************:************
AZ65B1       GAVNDICSDKTGTLTQGKMLARQIWIPRFGTITISNSDDPFNPNEGNVSLIPRFSPYEYS	420
ENA5         GAVNDICSDKTGTLTQGKMLARQIWIPRFGTITISNSDDPFNPNEGNVSLIPRFSPYEYS	420
AZ65A4       GAVNDICSDKTGTLTQGKMLARQIWIPRFGTITISNSDDPFNPNEGNVSLIPRFSPYEYS	420
AZ123A2      GAVNDICSDKTGTLTQGKMLARQIWIPRFGTITISNSDDPFNPNEGNVSLIPRFSPYEYS	420
ENA2         GAVNDICSDKTGTLTQGKMLARQIWIPRFGTITISNSDDPFNPNEGNVSLIPRFSPYEYS	420
AZ65A2       GAVNDICSDKTGTLTQGKMLARQIWIPRFGTITISNSDDPFNPNEGNVSLIPRFSPYEYS	420
AZ65A3       GAVNDICSDKTGTLTQGKMLARQIWIPRFGTITISNSDDPFNPNEGNVSLIPRFSPYEYS	420
AZ118B1      GAVNDICSDKTGTLTQGKMLARQIWIPRFGTITISNSDDPFNPNEGNVSLIPRFSPYEYS	420
AZ118A2      GAVNDICSDKTGTLTQGKMLARQIWIPRFGTITISNSDDPFNPNEGNVSLIPRFSPYEYS	420
ENA1         GAVNDICSDKTGTLTQGKMLARQIWIPRFGTITISNSDDPFNPNEGNVSLIPRFSPYEYS	420
AZ65A1       GAVNDICSDKTGTLTQGKMLARQIWIPRFGTITISNSDDPFNPNEGNVSLIPRFSPYEYS	420
AZ118A1      GAVNDICSDKTGTLTQGKMLARQIWIPRFGTITISNSDDPFNPNEGNVSLIPRFSPYEYS	420
AZ123A1      GAVNDICSDKTGTLTQGKMLARQIWIPRFGTITISNSDDPFNPNEGNVSLIPRFSPYEYS	420
             ************************************************************

AZ65B1       HNEDGDVGILQNFKDRLYEKDLPEDIDMDLFQKWLETATLANIATVFKDDATDCWKAHGD	480
ENA5         HNEDGDVGILQNFKDRLYEKDLPEDIDMDLFQKWLETATLANIATVFKDDATDCWKAHGD	480
AZ65A4       HNEDGDVGILQNFKDRLYEKDLPEDIDMDLFQKWLETATLANIATVFKDDATDCWKAHGD	480
AZ123A2      HNEDGDVGILQNFKDRLYEKDLPEDIDMDLFQKWLETATLANIATVFKDDATDCWKAHGD	480
ENA2         HNEDGDVGILQNFKDRLYEKDLPEDIDMDLFQKWLETATLANIATVFKDDATDCWKAHGD	480
AZ65A2       HNEDGDVGILQNFKDRLYEKDLPEDIDMDLFQKWLETATLANIATVFKDDATDCWKAHGD	480
AZ65A3       HNEDGDVGILQNFKDRLYEKDLPEDIDMDLFQKWLETATLANIATVFKDDATDCWKAHGD	480
AZ118B1      HNEDGDVGILQNFKDRLYEKDLPEDIDMDLFQKWLETATLANIATVFKDDATDCWKAHGD	480
AZ118A2      HNEDGDVGILQNFKDRLYEKDLPEDIDMDLFQKWLETATLANIATVFKDDATDCWKAHGD	480
ENA1         HNEDGDVGILQNFKDRLYEKDLPEDIDMDLFQKWLETATLANIATVFKDDATDCWKAHGD	480
AZ65A1       HNEDGDVGILQNFKDRLYEKDLPEDIDMDLFQKWLETATLANIATVFKDDATDCWKAHGD	480
AZ118A1      HNEDGDVGILQNFKDRLYEKDLPEDIDMDLFQKWLETATLANIATVFKDDATDCWKAHGD	480
AZ123A1      HNEDGDVGILQNFKDRLYEKDLPEDIDMDLFQKWLETATLANIATVFKDDATDCWKAHGD	480
             ************************************************************

AZ65B1       PTEIAIQVFATKMDLPRNALTGEKSNNQSNESDKSSLLGHNEKPSNAQFQHIAEFPFDST	540
ENA5         PTEIAIQVFATKMDLPHNALTGEKSTNQSNENDQSSLSQHNEKPGSAQFEHIAEFPFDST	540
AZ65A4       PTEIAIQVFATKMDLPRNALTGEKSTNQSNENDQSSLSQHNEKPGSAQFEHIAEFPFDST	540
AZ123A2      PTEIAIQVFATKMDLPRNALTGEKSTNQSNENDQSSLSQHNEKPGSAQFEHIAEFPFDST	540
ENA2         PTEIAIQVFATKMDLPHNALTGEKSTNQSNENDQSSLSQHNEKPGSAQFEHIAEFPFDST	540
AZ65A2       PTEIAIQVFATKMDLPRNALTGEKSTNQSNENDQSSLSQHNEKPGSAQFEHIAEFPFDST	540
AZ65A3       PTEIAIQVFATKMDLPRNALTGEKSTNQSNENDQSSLSQHNEKPGSAQFEHIAEFPFDST	540
AZ118B1      PTEIAIQVFATKMDLPRNALTGEKSTNQSNENDQSSLSQHNEKPGSAQFEHIAEFPFDST	540
AZ118A2      PTEIAIQVFATKMDLPRNALTGEKSTNQSNENDQSSLSQHNEKPGSAQFEHIAEFPFDST	540
ENA1         PTEIAIQVFATKMDLPHNALTGEKSTNQSNENDQSSLSQHNEKPGSAQFEHIAEFPFDST	540
AZ65A1       PTEIAIQVFATKMDLPRNALTGEKSTNQSNENDQSSLSQHNEKPGSAQFEHIAEFPFDST	540
AZ118A1      PTEIAIQVFATKMDLPRNALTGEKSTNQSNENDQSSLSQHNEKPGSAQFEHIAEFPFDST	540
AZ123A1      PTEIAIQVFATKMDLPRNALTGEKSTNQSNENDQSSLSQHNEKPGSAQFEHIAEFPFDST	540
             ****************:********.*****.*:***  *****..***:**********

AZ65B1       VKRMSSVYYNNHNETYNVYGKGAFESIISCCSSWYGKDGTKITPLTDCDVETIRKNVYSL	600
ENA5         VKRMSSVYYNNHNETYNIYGKGAFESIISCCSSWYGKDGVKITPLTDCDVETIRKNVYSL	600
AZ65A4       VKRMSSVYYNNHNETYNIYGKGAFESIISCCSSWYGKDGVKITPLTDCDVETIRKNVYSL	600
AZ123A2      VKRMSSVYYNNHNETYNIYGKGAFESIISCCSSWYGKDGVKITPLTDCDVETIRKNVYSL	600
ENA2         VKRMSSVYYNNHNETYNIYGKGAFESIISCCSSWYGKDGVKITPLTDCDVETIRKNVYSL	600
AZ65A2       VKRMSSVYYNNHNETYNIYGKGAFESIISCCSSWYGKDGVKITPLTDCDVETIRKNVYSL	600
AZ65A3       VKRMSSVYYNNHNETYNIYGKGAFESIISCCSSWYGKDGVKITPLTDCDVETIRKNVYSL	600
AZ118B1      VKRMSSVYYNNHNETYNIYGKGAFESIISCCSSWYGKDGVKITPLTDCDVETIRKNVYSL	600
AZ118A2      VKRMSSVYYNNHNETYNIYGKGAFESIISCCSSWYGKDGVKITPLTDCDVETIRKNVYSL	600
ENA1         VKRMSSVYYNNHNETYNIYGKGAFESIISCCSSWYGKDGVKITPLTDCDVETIRKNVYSL	600
AZ65A1       VKRMSSVYYNNHNETYNIYGKGAFESIISCCSSWYGKDGVKITPLTDCDVETIRKNVYSL	600
AZ118A1      VKRMSSVYYNNHNETYNIYGKGAFESIISCCSSWYGKDGVKITPLTDCDVETIRKNVYSL	600
AZ123A1      VKRMSSVYYNNHNETYNIYGKGAFESIISCCSSWYGKDGVKITPLTDCDVETIRKNVYSL	600
             *****************:*********************.********************

AZ65B1       SNEGLRVLGFASKSFTKDEQSDEQLKNNTSNRAAAESDLVFLGLIGIYDPPRNETAGAVK	660
ENA5         SNEGLRVLGFASKSFTKDQVNDDQLKNITSNRATAESDLVFLGLIGIYDPPRNETAGAVK	660
AZ65A4       SNEGLRVLGFASKSFTKDQVNDDQLKNITSNRATAESDLVFLGLIGIYDPPRNETAGAVK	660
AZ123A2      SNEGLRVLGFASKSFTKDQVNDDQLKNITSNRATAESDLVFLGLIGIYDPPRNETAGAVK	660
ENA2         SNEGLRVLGFASKSFTKDQVNDDQLKNITSNRATAESDLVFLGLIGIYDPPRNETAGAVK	660
AZ65A2       SNEGLRVLGFASKSFTKDQVNDDQLKNITSNRATAESDLVFLGLIGIYDPPRNETAGAVK	660
AZ65A3       SNEGLRVLGFASKSFTKDQVNDDQLKNITSNRATAESDLVFLGLIGIYDPPRNETAGAVK	660
AZ118B1      SNEGLRVLGFASKSFTKDQVNDDQLKNITSNRATAESDLVFLGLIGIYDPPRNETAGAVK	660
AZ118A2      SNEGLRVLGFASKSFTKDQVNDDQLKNITSNRATAESDLVFLGLIGIYDPPRNETAGAVK	660
ENA1         SNEGLRVLGFASKSFTKDQVNDDQLKNITSNRATAESDLVFLGLIGIYDPPRNETAGAVK	660
AZ65A1       SNEGLRVLGFASKSFTKDQVNDDQLKNITSNRATAESDLVFLGLIGIYDPPRNETAGAVK	660
AZ118A1      SNEGLRVLGFASKSFTKDQVNDDQLKNITSNRATAESDLVFLGLIGIYDPPRNETAGAVK	660
AZ123A1      SNEGLRVLGFASKSFTKDQVNDDQLKNITSNRATAESDLVFLGLIGIYDPPRNETAGAVK	660
             ******************: .*:**** *****:**************************
AZ65B1       KFHQAGINVHMLTGDFVGTAKAIAQEVGILPTNLYHYSQEIVDSMVMTGSQFDGLSEEEV	720
ENA5         KFHQAGINVHMLTGDFVGTAKAIAQEVGILPTNLYHYSQEIVDSMVMTGSQFDGLSEEEV	720
AZ65A4       KFHQAGINVHMLTGDFVGTAKAIAQEVGILPTNLYHYSQEIVDSMVMTGSQFDGLSEEEV	720
AZ123A2      KFHQAGINVHMLTGDFVGTAKAIAQEVGILPTNLYHYSQEIVDSMVMTGSQFDGLSEEEV	720
ENA2         KFHQAGINVHMLTGDFVGTAKAIAQEVGILPTNLYHYSQEIVDSMVMTGSQFDGLSEEEV	720
AZ65A2       KFHQAGINVHMLTGDFVGTAKAIAQEVGILPTNLYHYSQEIVDSMVMTGSQFDGLSEEEV	720
AZ65A3       KFHQAGINVHMLTGDFVGTAKAIAQEVGILPTNLYHYSQEIVDSMVMTGSQFDGLSEEEV	720
AZ118B1      KFHQAGINVHMLTGDFVGTAKAIAQEVGILPTNLYHYSQEIVDSMVMTGSQFDGLSEEEV	720
AZ118A2      KFHQAGINVHMLTGDFVGTAKAIAQEVGILPTNLYHYSQEIVDSMVMTGSQFDGLSEEEV	720
ENA1         KFHQAGINVHMLTGDFVGTAKAIAQEVGILPTNLYHYSQEIVDSMVMTGSQFDGLSEEEV	720
AZ65A1       KFHQAGINVHMLTGDFVGTAKAIAQEVGILPTNLYHYSQEIVDSMVMTGSQFDGLSEEEV	720
AZ118A1      KFHQAGINVHMLTGDFVGTAKAIAQEVGILPTNLYHYSQEIVDSMVMTGSQFDGLSEEEV	720
AZ123A1      KFHQAGINVHMLTGDFVGTAKAIAQEVGILPTNLYHYSQEIVDSMVMTGSQFDGLSEEEV	720
             ************************************************************

AZ65B1       DDLPVLPLVIARCSPQTKVRMIEALHRRKKFCAMTGDGVNDSPSLKMANVGIAMGINGSD	780
ENA5         DDLPVLPLVIARCSPQTKVRMIEALHRRKKFCAMTGDGVNDSPSLKMANVGIAMGINGSD	780
AZ65A4       DDLPVLPLVIARCSPQTKVRMIEALHRRKKFCAMTGDGVNDSPSLKMANVGIAMGINGSD	780
AZ123A2      DDLPVLPLVIARCSPQTKVRMIEALHRRKKFCAMTGDGVNDSPSLKMANVGIAMGINGSD	780
ENA2         DDLPVLPLVIARCSPQTKVRMIEALHRRKKFCAMTGDGVNDSPSLKMANVGIAMGINGSD	780
AZ65A2       DDLPVLPLVIARCSPQTKVRMIEALHRRKKFCAMTGDGVNDSPSLKMANVGIAMGINGSD	780
AZ65A3       DDLPVLPLVIARCSPQTKVRMIEALHRRKKFCAMTGDGVNDSPSLKMANVGIAMGINGSD	780
AZ118B1      DDLPVLPLVIARCSPQTKVRMIEALHRRKKFCAMTGDGVNDSPSLKMANVGIAMGINGSD	780
AZ118A2      DDLPVLPLVIARCSPQTKVRMIEALHRRKKFCAMTGDGVNDSPSLKMANVGIAMGINGSD	780
ENA1         DDLPVLPLVIARCSPQTKVRMIEALHRRKKFCTMTGDGVNDSPSLKMANVGIAMGINGSD	780
AZ65A1       DDLPVLPLVIARCSPQTKVRMIEALHRRKKFCAMTGDGVNDSPSLKMANVGIAMGINGSD	780
AZ118A1      DDLPVLPLVIARCSPQTKVRMIEALHRRKKFCAMTGDGVNDSPSLKMANVGIAMGINGSD	780
AZ123A1      DDLPVLPLVIARCSPQTKVRMIEALHRRKKFCAMTGDGVNDSPSLKMANVGIAMGINGSD	780
             ********************************:***************************

AZ65B1       VSKEASDIVLSDDNFASILNAVEEGRRMTDNIQKFVLQLLAENVAQALYLIIGLVFRDEN	840
ENA5         VSKEASDIVLSDDNFASILNAVEEGRRMTDNIQKFVLQLLAENVAQALYLIIGLVFRDEN	840
AZ65A4       VSKEASDIVLSDDNFASILNAVEEGRRMTDNIQKFVLQLLAENVAQALYLIIGLVFRDEN	840
AZ123A2      VSKEASDIVLSDDNFASILNAVEEGRRMTDNIQKFVLQLLAENVAQALYLIIGLVFRDEN	840
ENA2         VSKEASDIVLSDDNFASILNAVEEGRRMTDNIQKFVLQLLAENVAQALYLIIGLVFRDEN	840
AZ65A2       VSKEASDIVLSDDNFASILNAVEEGRRMTDNIQKFVLQLLAENVAQALYLIIGLVFRDEN	840
AZ65A3       VSKEASDIVLSDDNFASILNAVEEGRRMTDNIQKFVLQLLAENVAQALYLIIGLVFRDEN	840
AZ118B1      VSKEASDIVLSDDNFASILNAVEEGRRMTDNIQKFVLQLLAENVAQALYLIIGLVFRDEN	840
AZ118A2      VSKEASDIVLSDDNFASILNAVEEGRRMTDNIQKFVLQLLAENVAQALYLIIGLVFRDEN	840
ENA1         VSKEASDIVLSDDNFASILNAVEEGRRMTDNIQKFVLQLLAENVAQALYLIIGLVFRDEN	840
AZ65A1       VSKEASDIVLSDDNFASILNAVEEGRRMTDNIQKFVLQLLAENVAQALYLIIGLVFRDEN	840
AZ118A1      VSKEASDIVLSDDNFASILNAVEEGRRMTDNIQKFVLQLLAENVAQALYLIIGLVFRDEN	840
AZ123A1      VSKEASDIVLSDDNFASILNAVEEGRRMTDNIQKFVLQLLAENVAQALYLIIGLVFRDEN	840
             ************************************************************

AZ65B1       GKSVFPLSPVEVLWIIVVTSCFPAMGLGLEKAAPDLMDRPPNDSEVGIFTWEVIIDTFAY	900
ENA5         GKSVFPLSPVEVLWIIVVTSCFPAMGLGLEKAAPDLMDRPPNDSEVGIFTWEVIIDTFAY	900
AZ65A4       GKSVFPLSPVEVLWIIVVTSCFPAMGLGLEKAAPDLMDRPPNDSEVGIFTWEVIIDTFAY	900
AZ123A2      GKSVFPLSPVEVLWIIVVTSCFPAMGLGLEKAAPDLMDRPPNDSEVGIFTWEVIIDTFAY	900
ENA2         GKSVFPLSPVEVLWIIVVTSCFPAMGLGLEKAAPDLMDRPPHDSEVGIFTWEVIIDTFAY	900
AZ65A2       GKSVFPLSPVEVLWIIVVTSCFPAMGLGLEKAAPDLMDRPPNDSEVGIFTWEVIIDTFAY	900
AZ65A3       GKSVFPLSPVEVLWIIVVTSCFPAMGLGLEKAAPDLMDRPPNDSEVGIFTWEVIIDTFAY	900
AZ118B1      GKSVFPLSPVEVLWIIVVTSCFPAMGLGLEKAAPDLMDRPPNDSEVGIFTWEVIIDTFAY	900
AZ118A2      GKSVFPLSPVEVLWIIVVTSCFPAMGLGLEKAAPDLMDRPPNDSEVGIFTWEVIIDTFAY	900
ENA1         GKSVFPLSPVEVLWIIVVTSCFPAMGLGLEKAAPDLMDRPPHDSEVGIFTWEVIIDTFAY	900
AZ65A1       GKSVFPLSPVEVLWIIVVTSCFPAMGLGLEKAAPDLMDRPPNDSEVGIFTWEVIIDTFAY	900
AZ118A1      GKSVFPLSPVEVLWIIVVTSCFPAMGLGLEKAAPDLMDRPPNDSEVGIFTWEVIIDTFAY	900
AZ123A1      GKSVFPLSPVEVLWIIVVTSCFPAMGLGLEKAAPDLMDRPPNDSEVGIFTWEVIIDTFAY	900
             *****************************************:******************
AZ65B1       GFLMAGSCMASFTGSLYGINNGELGVNCDRSYNGSCHDVYRSRSAAFATMTWCALILAWE	960
ENA5         GIIMTGSCMASFTGSLYGINSGRLGHDCDGTYNSSCRDVYRSRSAAFATMTWCALILAWE	960
AZ65A4       GIIMTGSCMASFTGSLYGINSGRLGHDCDGTYNSSCRDVYRSRSAAFATMTWCALILAWE	960
AZ123A2      GIIMTGSCMASFTGSLYGINSGRLGHDCDGTYNSSCRDVYRSRSAAFATMTWCALILAWE	960
ENA2         GIIMTGSCMASFTGSLYGINSGRLGHDCDGTYNSSCRDVYRSRSAAFATMTWCALILAWE	960
AZ65A2       GIIMTGSCMASFTGSLYGINSGRLGHDCDGTYNSSCRDVYRSRSAAFATMTWCALILAWE	960
AZ65A3       GIIMTGSCMASFTGSLYGINSGRLGHDCDGTYNSSCRDVYRSRSAAFATMTWCALILAWE	960
AZ118B1      GIIMTGSCMASFTGSLYGINSGRLGHDCDGTYNSSCRDVYRSRSAAFATMTWCALILAWE	960
AZ118A2      GIIMTGSCMASFTGSLYGINSGRLGHDCDGTYNSSCRDVYRSRSAAFATMTWCALILAWE	960
ENA1         GIIMTGSCMASFTGSLYGINSGRLGHDCDGTYNSSCRDVYRSRSAAFATMTWCALILAWE	960
AZ65A1       GIIMTGSCMASFTGSLYGINSGRLGHDCDGTYNSSCRDVYRSRSAAFATMTWCALILAWE	960
AZ118A1      GIIMTGSCMASFTGSLYGINSGRLGHDCDGTYNSSCRDVYRSRSAAFATMTWCALILAWE	960
AZ123A1      GIIMTGSCMASFTGSLYGINSGRLGHDCDGTYNSSCRDVYRSRSAAFATMTWCALILAWE	960
             *::*:***************.*.** :** :**.**:***********************

AZ65B1       VVDMRRSFFRMHPDTDSPVKEFFKSIWGNQFLFWSIIFGFVSAFPVVYIPVINDKVFLHK	1020
ENA5         VVDMRRSFFRMHPDTDSPVKEFFRSIWGNQFLFWSIIFGFVSAFPVVYIPVINDKVFLHK	1020
AZ65A4       VVDMRRSFFRMHPDTDSPVKEFFRSIWGNQFLFWSIIFGFVSAFPVVYIPVINDKVFLHK	1020
AZ123A2      VVDMRRSFFRMHPDTDSPVKEFFRSIWGNQFLFWSIIFGFVSAFPVVYIPVINDKVFLHK	1020
ENA2         VVDMRRSFFRMHPDTDSPVKEFFRSIWGNQFLFWSIIFGFVSAFPVVYIPVINDKVFLHK	1020
AZ65A2       VVDMRRSFFRMHPDTDSPVKEFFRSIWGNQFLFWSIIFGFVSAFPVVYIPVINDKVFLHK	1020
AZ65A3       VVDMRRSFFRMHPDTDSPVKEFFRSIWGNQFLFWSIIFGFVSAFPVVYIPVINDKVFLHK	1020
AZ118B1      VVDMRRSFFRMHPDTDSPVKEFFRSIWGNQFLFWSIIFGFVSAFPVVYIPVINDKVFLHK	1020
AZ118A2      VVDMRRSFFRMHPDTDSPVKEFFRSIWGNQFLFWSIIFGFVSAFPVVYIPVINDKVFLHK	1020
ENA1         VVDMRRSFFRMHPDTDSPVKEFFRSIWGNQFLFWSIIFGFVSAFPVVYIPVINDKVFLHK	1020
AZ65A1       VVDMRRSFFRMHPDTDSPVKEFFRSIWGNQFLFWSIIFGFVSAFPVVYIPVINDKVFLHK	1020
AZ118A1      VVDMRRSFFRMHPDTDSPVKEFFRSIWGNQFLFWSIIFGFVSAFPVVYIPVINDKVFLHK	1020
AZ123A1      VVDMRRSFFRMHPDTDSPVKEFFRSIWGNQFLFWSIIFGFVSAFPVVYIPVINDKVFLHK	1020
             ***********************:************************************

AZ65B1       PIGPEWGLAIAFTIAFWIGAELYKLGKRRYFKTQRAHNPENDLESNNKRDPFEAYSTSTT	1080
ENA5         PIGAEWGLAIAFTIAFWIGAELYKCGKRRYFKTQRAHNSENDLERSSKHDPFEAYSTSTT	1080
AZ65A4       PIGAEWGLAIAFTIAFWVGAELYKCGKRRYFKTQRAHNSENDLERSSKHDPFEAYSTSTT	1080
AZ123A2      PIGAEWGLAIAFTIAFWIGAELYKCGKRRYFKTQRAHNPENDLELSSKHDPFEAYSTSTT	1080
ENA2         PIGAEWGLAIAFTIAFWIGAELYKCGKRRYFKTQRAHNPENDLESNNKRDPFEAYSTSTT	1080
AZ65A2       PIGAEWGLAIAFTIAFWIGAELYKCGKRRYFKTQRAHNPENDLESNNKRDPFEAYSTSTT	1080
AZ65A3       PIGAEWGLAIAFTIAFWIGAELYKCGKRRYFKTQRAHNPENDLESNNKRDPFEAYSTSTT	1080
AZ118B1      PIGAEWGLAIAFTIAFWIGAELYKCGKRRYFKTQRAHNPENDLESNNKRDPFEAYSTSTT	1080
AZ118A2      PIGAEWGLAIAFTIAFWIGAELYKCGKRRYFKTQRAHNPENDLESNNKRGPFEAYSTSTT	1080
ENA1         PIGAEWGLAIAFTIAFWIGAELYKCGKRRYFKTQRAHNPENDLESNNKRDPFEAYSTSTT	1080
AZ65A1       PIGAEWGLAIAFTIAFWIGAELYKCGKRRYFKTQRAHNPENDLESNNKRDPFEAYSTSTT	1080
AZ118A1      PIGAEWGLAIAFTIAFWIGAELYKCGKRRYFKTQRAHNPENDLESNNKRDPFEAYSTSTT	1080
AZ123A1      PIGAEWGLAIAFTIAFWIGAELYKCGKRRYFKTQRAHNPENDLESNNKRDPFEAYSTSTT	1080
             *** *************:****** ************* ***** ..*:.**********

AZ65B1       IHTEINIGIKQ	1091
ENA5         LQSEINISVKH	1091
AZ65A4       LQSEINISVKH	1091
AZ123A2      LQSEINISVKH	1091
ENA2         IHTEVNIGIKQ	1091
AZ65A2       IHTEANIGIKQ	1091
AZ65A3       IHTEANIGIKQ	1091
AZ118B1      IHTEANIGIKQ	1091
AZ118A2      IHTEINIGIKQ	1091
ENA1         IHTEVNIGIKQ	1091
AZ65A1       IHTEANIGIKQ	1091
AZ118A1      IHTEANIGIKQ	1091
AZ123A1      IHTEANIGIKQ	1091
             :::* **.:*:
```
