## Supplemental Figure 3 for "Analysis of genome sequences of marine isolates of *Saccharomyces cerevisiae*"

**Supplementary Figure S3: Clustal-Omega Alignment of *RSM10*, intergenic promoter region and first *ENA* copy in reference strain S288C and marine strains AZ65, AZ118 and AZ123.** The sequences begin with the stop codon of *RSM10* (in reverse orientation) and end with the stop codon of the first *ENA* copy. Start and stop codons are highlighted in green. Bases where the sequence differs from that of S288C are highlighted in red. Apparent indels which could cause frameshifts in the *ENA* sequence are highlighted in cyan. The intergenic region containing the promoters of *RSM10* and *ENA1* runs from 613 to 1947 in the S288C sequence.

```
AZ65       TTACTTCTTTTCCAAATGTTTTTTAAAATCGGGGCTATCCAATAGTTCCAATACTTTTTG	60
S288C      TTACTTCTTTTCCAAATGTTTTTTAAAATCGGGGCTATCCAATAGTTCCAATACTTTTTG	60
AZ118      TTACTTCTTTTCCAAATGTTTTTTAAAATCGGGGCTATCCAATAGTTCCAATACTTTTTG	60
AZ123      TTACTTCTTTTCCAAATGTTTTTTAAAATCGGGGCTATCCAATAGTTCCAATACTTTTTG	60
           ************************************************************

AZ65       TCCGACAGCACTAGTTTGAGCTTTATCATCGTTTCTCAGAGAATACAATTCATCGATATT	120
S288C      TCCGACAGCACTAGTTTGAGCTTTATCATCGTTTCTCAGAGAATACAATTCATCGATATT	120
AZ118      TCCGACAGCACTAGTTTGAGCTTTATCATCGTTTCTCAGAGAATACAATTCATCGATATT	120
AZ123      TCCGACAGCACTAGTTTGAGCTTTATCATCGTTTCTCAGAGAATACAATTCATCGATATT	120
           ************************************************************

AZ65       GCTTAGTGATTTTTCGAGGCCATTGGCGTCCGAACCCAAGTCCAGCGATATTTCGGACCT	180
S288C      GCTTAGTGATTTTTCGAGGCCATTGGCGTCCGAACCCAAGTCCAGCGATATTTCGGACCT	180
AZ118      GCTTAGTGATTTTTCGAGGCCATTGGCGTCCGAACCCAAGTCCAGGGATATTTCGGACCT	180
AZ123      GCTTAGTGATTTTTCGAGGCCATTGGCGTCCGAACCCAAGTCCAGCGATATTTCGGACCT	180
           ********************************************* **************

AZ65       TTGAAAAAAGTTACATTTCATTCCGACGCCTGCCATAGAGTGCTTCGTAATGTAAGCGAT	240
S288C      TTGAAAGAAGTTACATTTCATTCCGACGCCTGCCATAGAGTGCTTCGTAATGTAAGCTAT	240
AZ118      TTGAAAGAAGTTACATTTCATTCCGACGCCTGCCATAGAGTGCTTCGTAATGTAAGCTAT	240
AZ123      TTGAAAAAAGTTACATTTCATTCCGACGCCTGCCATAGAGTGCTTCGTAATGTAAGCTAT	240
           ****** ************************************************** **

AZ65       CAACATCTGTAGCACTTCGGGGTTCGTATCCCAGGCCCTAATGAGCCTCTTATGTGTATG	300
S288C      CAACATCTGTAGCACTTCGGGGTTCGTATCCCAGGCCCTAATGAGCCTCTTATGTGTATG	300
AZ118      CAACATCTGTAGCACTTCGGGGTTCGTATCCCAGGCCCTAATGAGCCTCTTATGTGTATG	300
AZ123      CAACATCTGTAGCACTTCGGGGTTCGTATCCCAGGCCCTAATGAGCCTCTTATGTGTATG	300
           ************************************************************

AZ65       TCTTTCAAAATTCTCTTTGGATTTAGCGTGCACGAAAGGTGATTTTATTACAGTCCACCT	360
S288C      TCTTTCAAAATTCTCTTTGGATTTAGCGTGCACGAAAGGTGATTTTATTACAGTCCACCT	360
AZ118      TCTTTCAAAATTCTCTTTGGATTTAGCGTGCACGAAAGGTGATTTTATTACAGTCCACCT	360
AZ123      TCTTTCAAAATTCTCTTTGGATTTAGCGTGCACGAAAGGTGATTTTATTACAGTCCACCT	360
           ************************************************************

AZ65       TTCTCTTCTCGTGGGCAAAGGTTTAGGACCTGTCAATGGGATACCCAAATAGTAACCTGT	420
S288C      TTCTCTTCTCGTGGGCAAAGGTTTAGGACCTGTCAATGGGATACCCAAATAGTAACCTGT	420
AZ118      TTCTCTTCTCGTGGGCAAAGGTTTAGGACCTGTCAATGGGATACCCAAATAGTAACCTGT	420
AZ123      TTCTCTTCTCGTGGGCAAAGGTTTAGGACCTGTCAATGGGATACCCAAATAGTAACCTGT	420
           ************************************************************
```

```
AZ65       TCTAAGTATGAAGTCTGAATAAAAATCGAGGTTTTCATTGTCATACGATCTTAACTGAAT	480
S288C      TCTAAGTATGAAGTCTGAATAAAAATCGAGGTTTTCATTGTCATACGATCTTAACTGAAT	480
AZ118      TCTAAGTATGAAGTCTGAATAAAAATCGAGGTTTTCATTGTCATACGATCTTAACTGAAT	480
AZ123      TCTAAGTATGAAGTCTGAATAAAAATCGAGGTTTTCATTGTCATACGATCTTAACTGAAT	480
           ************************************************************

AZ65       GTCTGCAACTAAATCACCATATTTGATAGGTAACTTTAAAGGAGCATAGTAGACTGCCTC	540
S288C      GTCTGCAACTAAATCACCATATTTGATAGGTAACTTTAAAGGAGCATAGTAGACTGCCTC	540
AZ118      GTCTGCAACTAAATCACCATATTTGATAGGTAACTTTAAAGGAGCATAGTAGACTGCCTC	540
AZ123      GTCTGCAACTAAATCACCATATTTGATAGGTAACTTTAAAGGAGCATAGTAGACTGCCTC	540
           ************************************************************

AZ65       TACATTTACAGGATATGGGCGCGTCGACTGTGTTCTGATAAATGATCTTAATGCAATGGT	600
S288C      TACATTTACAGGATATGGGCGCGTCGACTGTGTTCTGATAAATGATCTTAATGCAATGGT	600
AZ118      TACATTTACAGGATATGGGCGCGTCGACTGTGTTCTGATAAATGATCTTAATGCAATGGT	600
AZ123      TACATTTACAGGATATGGGCGCGTCGACTGTGTTCTGATAAATGATCTTAATGCAATGGT	600
           ************************************************************

AZ65       ATTTCTAAGCATTGGCCAATGTTTTTC-CCCTTTTCTGATTTTGCACGGTTCGCATAATT	659
S288C      ATTTCTAAGCATTGGCCAATGTTTTTCCCCCTTTTCTGATTTTGCACGGTTCGCATAATT	660
AZ118      ATTTCTAAGCATTGGCCAATGTTTTTCCCCCTTTTCTGATTTTGCACGGTTCGCATAATT	660
AZ123      ATTTCTAAGCATTGGCCAATGTTTTTCCCCCTTTTCTGATTTTGCACGGTTCGCATAATT	660
           *************************** ********************************

AZ65       CTAATTTGTACCAAATAGTTTCATTTCATGCCCTTTCTTGCATTTTTTCCTTTTCAAAAG	719
S288C      CTAATTTGTACCAAATAGTTTCATTTCATGCCCTTTCTTGCATTTTTTCCTTTTCAAAAG	720
AZ118      CTAATTTGTACCAAATAGTTTCATTTCATGCCCTTTCTTGCATTTTTTCCTTTTCAAAAG	720
AZ123      CTAATTTGTACCAAATAGTTTCATTTCATGCCCTTTCTTGCATTTTTTCCTTTTCAAAAG	720
           ************************************************************

AZ65       AAAATAGATGTACAGGTATTACCCGCCTGGTACATAGTTTGCTAACATTCTTGCCCAATC	779
S288C      AAAATAGATGTACAGGTATTACCCGCCTGGTACATAGTTTGCTAACATTCTTGCCCAATC	780
AZ118      AAAATAGATGTACAGGTATTACCCGCCTGGTACATAGTTTGCTAACATTCTTGCCCAATC	780
AZ123      AAAATAGATGTACAGGTATTACCCGCCTGGTACATAGTTTGCTAACATTCTTGCCCAATC	780
           ************************************************************

AZ65       TCTCTGCTGTTAATAAAGGATGTGTGGATACATGACTAATTTCCGGATTAAACAGTGACG	839
S288C      TCTCTGCTGTTAATAAAGGATGTGTGGATACATGACTAATTTCCGGATTAAACAGTGACG	840
AZ118      TCTCTGCTGTTAATAAAGGATGTGTGGATACATGACTAATTTCCGGATTAAACAGTGACG	840
AZ123      TCTCTGCTGTTAATAAAGGATGTGTGGATACATGACTAATTTCCGGATTAAACAGTGACG	840
           ************************************************************

AZ65       ATAAGAAACGGTATGGCAAGAGTCAGAATTTGATAAGACTTCAGTAATATTGATTATCGG	899
S288C      ATAAGAAACGGTATGGCAAGAGTCAGAATTTGATAAGACTTCAGTAATATTGATTATCGG	900
AZ118      ATAAGAAACGGTATGGCAAGAGTCAGAATTTGATAAGACTTCAGTAATATTGATTATCGG	900
AZ123      ATAAGAAACGGTATGGCAAGAGTCAGAATTTGATAAGACTTCAGTAATATTGATTATCGG	900
           ************************************************************

AZ65       TCGAACTAAGCTACCCTCTTGGTTAAATTCTAATGACTACCAGTGTGGTCAATATTTTAG	959
S288C      TCGAACTAAGCTACCCTCTTGGTTAAATTCTAATGACTACCAGTGTGGTCAATATTTTAG	960
AZ118      TCGAACTAAGCTACCCTCTTGGTTAAATTCTAATGACTATCAGTGTGGTCAATATTTTAG	960
AZ123      TCGAACTAAGCTACCCTCTTGGTTAAATTCTAATGACTATCAGTGTGGTCAATATTTTAG	960
           *************************************** ********************

AZ65       GGTTATCGGTGAACGATAACAAAATCCGTAATTGTTTCCTACGCTGGTTATTACTAAGTA	1019
S288C      GGTTATCGGTGAACGATAACAAAATCCGTAATTGTTTCCTACGCTGGTTATTACTAAGTA	1020
AZ118      GGTTATCGGTGAACGATAACAAAATCCGTAATTGTTTCCTACGCTGGTTATTACTAAGTA	1020
AZ123      GGTTATCGGTGAACGATAACAAAATCCGTAATTGTTTCCTACGCTGGTTATTACTAAGTA	1020
           ************************************************************

AZ65       GCCACTCGGTGGCTATCTCTAGCCGTTCATCAATCATGTCAAAAGTACAATCACTGAGCT	1079
S288C      GCCACTCGGTGGCTATCTCTAGCCGTTCATCAATCATGTCAAAAGTACAATCACTGAGCT	1080
AZ118      GCCACTCGGTGGCTATCTCTAGCCGTTCATCAATCATGTCAAAAGTACAATCACTGAGCT	1080
AZ123      GCCACTCGGTGGCTATCTCTAGCCGTTCATCAATCATGTCAAAAGTACAATCACTGAGCT	1080
           ************************************************************

AZ65       TATTAAGTCATCCGCACGTGGAACTCACACAAGCCCCGAATTTTGTCAGCCATTCCGTGG	1139
S288C      TATTAAGTCATCCGCACGTGGAACTCACACAAGCCCCGAATTTTGTCAGCCATTCCGTGG	1140
AZ118      TATTAAGTCATCCGCACGTGGAACTCACACAAGCCCCGAATTTTGTCAGCCATTCCGTGG	1140
AZ123      TATTAAGTCATCCGCACGTGGAACTCACACAAGCCCCGAATTTTGTCAGCCATTCCGTGG	1140
           ************************************************************
AZ65       AAAATCCTTTATCTCGTCACTCTTGTACGGAAGTTGCACCTTTAGCCCCACTCCCAGATC	1199
S288C      AAAATCCTTTATCTCGTCACTCTTGTACGGAAGTTGCACCTTTAGCCCCACTCCCAGATC	1200
AZ118      AAAATCCTTTATCTCGTCACTCTTGTACGGAAGTTGCACCTTTAGCCCCACTCCCAGATC	1200
AZ123      AAAATCCTTTATCTCGTCACTCTTGTACGGAAGTTGCACCTTTAGCCCCACTCCCAGATC	1200
           ************************************************************

AZ65       TTCAATCGCAGAGGCAATTAGGGTGGCTGTGCAATGTAAACTTTCTTATCGCTTCCTGTT	1259
S288C      TTCAATCACAGAGGCAATGAGGGTGGCTGTGCAATGTAAACTTTCTTATCGCTTCCTGTT	1260
AZ118      TTCAATCGCAGAGGCAATGAGGGTGGCTGTGCAATGTAAACTTTCTTATCGCTTCCTGTT	1260
AZ123      TTCAATCACAGAGGCAATGAGGGTGGCTGTGCAATGTAAACTTTCTTATCGCTTCCTGTT	1260
           ******* ********** *****************************************

AZ65       ATGATTGATTACCATGCGCGGTAGAAGGATTGAATGAGGGGTGTGGGTGTGATAATTCAT	1319
S288C      ATGATTGATTACCATGCGCGGTAGAAGGATTGAATGAGGGGTGTGGGTGTGATAATTCAT	1320
AZ118      ATGATTGATTACCATGCGCGGTAGAAGGATTGAATGAGGGGTGTGGGTGTGATAATTCAT	1320
AZ123      ATGATTGATTACCATGCGCGGTAGAAGGATTGAATGAGGGGTGTGGGTGTGATAATTCAT	1320
           ************************************************************

AZ65       GTACGGACCCTTTCCTCCCAGAAGCTTCTGCACGAAGTGGTTACATTGTGAGTACAGGGT	1379
S288C      GTACGGACCCTTTCCTCCCAGAAGCTTCTGCACGAAGTGGTTACATTGTGAGTACAGGGT	1380
AZ118      GTACGGACCCTTTCCTCCCAGAAGCTTCTGCACGAAGTGGTTACATTGTGAGTACAGGGT	1380
AZ123      GTACGGACCCTTTCCTCCCAGAAGCTTCTGCACGAAGTGGTTACATTGTGAGTACAGGGT	1380
           ************************************************************

AZ65       CTTAAACATCGCCGTGCGGCGTCATTTTGCGGGGCTCATCATTTATTTCCTACTTCTATG	1439
S288C      CTTAAACATCGCCGTGCGGCGTCATTTTGCGGGGCTCATCATTTATTTCCTACTTCTATG	1440
AZ118      CTTAAACATCGCCGTGCGGCGTCATTTTGCGGGGCTCATCATTTATTTCCTACTTCTATG	1440
AZ123      CTTAAACATCGCCGTGCGGCGTCATTTTGCGGGGCTCATCATTTATTTCCTACTTCTATG	1440
           ************************************************************

AZ65       ACGTTTGTTAGGGCAGGGATGTAGATTTTGGTCATATACTTTCACCTGACAGAAGAAAAA	1499
S288C      ACGTTTGTTAGGGCAGGGATGTAGATTTTGGTCATATACTTTCACCTGACAGAAGAAAAA	1500
AZ118      ACGTTTGTTAGGGCAGGGATGTAGATTTTGGTCATATACTTTCACCTGACAGAAGAAAAA	1500
AZ123      ACGTTTGTTAGGGCAGGGATGTAGATTTTGGTCATATACTTTCACCTGACAGAAGAAAAA	1500
           ************************************************************

AZ65       ACAAGGTCTTAGAGCAGTATTCCCGAATTATTGCCTCTTTCCTTGGTGGGTGCGGGTTAA	1559
S288C      ACAAGGTCTTAGAGCAGTATTCCCGAATTATTGCCTCTTTCCTTGGTGGGTGCGGGTTAA	1560
AZ118      ACAAGGTCTTAGAGCAGTATTCCCGAATTATTGCCTCTTTCCTTGGTGAGTGCGGGTTAA	1560
AZ123      ACAAGGTCTTAGAGCAGTATTCCCGAATTATTGCCTCTTTCCTTGGTGGGTGCGGGTTAA	1560
           ************************************************ ***********

AZ65       TATTAGATGAACGGTGGGTTGTGGTCAAACCCGGGGTAGGTTTTAAAATGTCATGCGTTT	1619
S288C      TATTAGATGAACGGTGGGTTGTGGTCAAACCCGGGGTAGGTTTTAAAATGTCATGCGTTT	1620
AZ118      TATTAGATGAACGGTGGGTTGTGGTCAAACCCGGGGTAGGTTTTAAAATGTCATGCGTTT	1620
AZ123      TATTAGATGAACGGTGGGTTGTGGTCAAACCCGGGGTAGGTTTTAAAATGTCATGCGTTT	1620
           ************************************************************

AZ65       TCGTCATTTACATAGATATATACAGGGAAAATATATATA------------TATATATAT	1667
S288C      TCGTCATTTACATAGATATATACAGGGAAAATATATATATATATATATATATATATATAT	1680
AZ118      TCGTCATTTACATAGATATATACAGGGAAAATATATATATA----------TATATATAT	1670
AZ123      TCGTCATTTACATAGATATATACAGGGAAAATATATATA------------TATATATAT	1668
           *************************************              *********

AZ65       ACCTAAGGGAAACAATTTGAATAGATAAAAGGTCTTTTCCTCTTATTCCGTTCTTTCTTT	1727
S288C      ACCTAAGGGAAACAATTTGAATAGATAAAAGGTCTTTTCCTCTTATTCCGTTCTTTCTTT	1740
AZ118      ACCTAAGGGAAACAATTTGAATAGATAAAAGGTCTTTTCCTCTTATTCCGTTCTTTCTTT	1730
AZ123      ACCTAAGGGAAACAATTTGAATAGATAAAAGGTCTTTTCCTCTTATTCCGTTCTTTCTTT	1728
           ************************************************************
AZ65       TTAGAGTACTATTAATTTCTACTATTTCTCCTATCTTTCTTTTCTCATTAGTAATAGTCT	1787
S288C      TTAGAGTACTATTAATTTCTACTATTTCTCCTATCTTTCTTTTCTCATTAGTAATAGTCT	1800
AZ118      TTAGAGTACTATTAATTTCTACTATTTCTCCTATCTTTCTTTTCTCATTAGTAATAGTCT	1790
AZ123      TTAGAGTACTATTAATTTCTACTATTTCTCCTATCTTTCTTTTCTCATTAGTAATAGTCT	1788
           ************************************************************

AZ65       AATCATTTATATTCTAATATTAACTTTAATAAAGCAGATATCTTCATCAAGTGAACTGTT	1847
S288C      AATCATTTATATTCTAATATTAACTTTAATAAAGCAGATATCTTCATCAAGTGAACTGTT	1860
AZ118      AATCATTTATATTCTAATATTAACTTTAATAAAGCAGATATCTTCATCAAGTGAACTGTT	1850
AZ123      AATCATTTATATTCTAATATTAACTTTAATAAAGCAGATATCTTCATCAAGTGAACTGTT	1848
           ************************************************************

AZ65       CACACTTAATTCACATTTATTTGAGGCAACAATACGTATATTTACTAATTAAAGAAAAAA	1907
S288C      CACACTTAATTCACATTTATTTGAGGCAACAATACGTATATTTACTAATTAAAGAAAAAA	1920
AZ118      CACACTTAATTCACATTTATTTGAGGCAACAATACGTATATTTACTAATTAAAGAAAAAA	1910
AZ123      CACACTTAATTCACATTTATTTGAGGCAACAATACGTATATTTACTAATTAAAGAAAAAA	1908
           ************************************************************

AZ65       CTTCGTACACAGAATTGAAAATTTTCGATGGGCGAAGGAACTACTAAGGAAAACAATAAT	1967
S288C      CTTCGTACACAGAATTGAAAATTTTCGATGGGCGAAGGAACTACTAAGGAAAACAATAAT	1980
AZ118      CTTCGTACACAGAATTGAAAATTTTCGATGGGCGAAGGAACTACTAAGGAAAACAATAAT	1970
AZ123      CTTCGTACACAGAATTGAAAATTTTCGATGGGCGAAGGAACTACTAAGGAAAACAATAAT	1968
           ************************************************************

AZ65       GCAGAATTCAATGCTTATCACACGCTGACTGCAGAAGAAGCCGCTGAATTCATAGGCACA	2027
S288C      GCAGAATTCAATGCTTATCACACGCTGACTGCAGAAGAAGCCGCTGAATTCATAGGCACA	2040
AZ118      GCAGAATTCAATGCTTATCACACGCTGACTGCAGAAGAAGCCGCTGAATTCATAGGCACA	2030
AZ123      GCAGAATTCAATGCTTATCACACGCTGACTGCAGAAGAAGCCGCTGAATTCATAGGCACA	2028
           ************************************************************

AZ65       AGTCTAACTGAAGGTTTGACCCAAGATGAGTTCGTCCACAGATTGAAAACAGTGGGTGAG	2087
S288C      AGTCTAACTGAAGGTTTGACCCAAGATGAGTTCGTCCACAGATTGAAAACAGTGGGTGAG	2100
AZ118      AGTCTAACTGAAGGTTTGACCCAAGATGAGTTCGTCCACAGATTGAAAACAGTGGGTGAG	2090
AZ123      AGTCTAACTGAAGGTTTGACCCAAGATGAGTTCGTCCACAGATTGAAAACAGTGGGTGAG	2088
           ************************************************************

AZ65       AACACATTGGGTGATGACACTAAAATTGATTACAAGGCAATGGTCCTCCATCAGGTATGT	2147
S288C      AACACATTGGGTGATGACACTAAAATTGATTACAAGGCAATGGTCCTCCATCAGGTATGT	2160
AZ118      AACACATTGGGTGATGACACTAAAATTGATTACAAGGCAATGGTCCTCCATCAGGTATGT	2150
AZ123      AACACATTGGGTGATGACACTAAAATTGATTACAAGGCAATGGTCCTCCATCAGGTATGT	2148
           ************************************************************

AZ65       AATGCCATGATCATGGTCCTTCTAATATCCATGATAATCTCGTTTGCCATGCATGATTGG	2207
S288C      AATGCCATGATCATGGTCCTTCTAATATCCATGATAATCTCGTTTGCCATGCATGATTGG	2220
AZ118      AATGCCATGATCATGGTCCTTCTAATATCCATGATAATCTCGTTTGCCATGCATGATTGG	2210
AZ123      AATGCCATGATCATGGTCCTTCTAATATCCATGATAATCTCGTTTGCCATGCATGATTGG	2208
           ************************************************************

AZ65       ATTACTGGTGGTGTTATTTCCTTTGTCGTGGGTGTTAACGTCGGTATTGGCCTTGTTCAA	2267
S288C      ATTACTGGTGGCGTTATTTCTTTTGTTATCGCGGTCAATGTGCTCATTGGCCTAGTTCAA	2280
AZ118      ATTACTGGTGGTGTTATTTCCTTTGTCGTGGGTGTTAACGTCGGTATTGGCCTTGTTCAA	2270
AZ123      ATTACTGGTGGTGTTATTTCCTTTGTCGTGGGTGTTAACGTCGGTATTGGCCTTGTTCAA	2268
           *********** ******** *****  * *  ** ** **    ******** ******

AZ65       GAATATAAGGCTACAAAGACAATGAACTCTTTGAAAAACTTGAGCTCTCCCAATGCTCAT	2327
S288C      GAATATAAGGCTACCAAGACAATGAACTCTTTGAAAAACTTGAGCTCTCCCAATGCTCAT	2340
AZ118      GAATATAAGGCTACAAAGACAATGAACTCTTTGAAAAACTTGAGCTCTCCCAATGCTCAT	2330
AZ123      GAATATAAGGCTACAAAGACAATGAACTCTTTGAAAAACTTGAGCTCTCCCAATGCTCAT	2328
           ************** *********************************************
AZ65       GTTATTAGGAACGGGAAAAGTGAGACTATAAACTCCAAAGATGTGGTTCCAGGTGATATT	2387
S288C      GTTATTAGGAACGGGAAAAGTGAGACTATAAACTCAAAAGATGTGGTTCCAGGTGATATT	2400
AZ118      GTTATCAGAAACGGGAAGAGCGAAACTATAAATTCCAAAGATGTCGTTCCAGGTGATATT	2390
AZ123      GTTATCAGAAACGGGAAGAGCGAAACTATAAATTCCAAAGATGTCGTTCCAGGTGATATT	2388
           ***** ** ******** ** ** ******** ** ******** ***************

AZ65       TGTCTGGTGAAGGTCGGTGATACTATTCCTGCTGATTTGCGTTTGATTGAAACTAAGAAT	2447
S288C      TGTCTGGTGAAGGTCGGTGATACTATTCCTGCTGATTTGCGTTTAATTGAAACTAAGAAT	2460
AZ118      TGTCTAGTGAAGGTCGGTGACACTATTCCTGCTGATTTACGTTTGATTGAAACCAAGAAT	2450
AZ123      TGTCTAGTGAAGGTCGGTGACACTATTCCTGCTGATTTACGTTTGATTGAAACCAAGAAT	2448
           ***** ************** ***************** ***** ******** ******

AZ65       TTTGATACTGATGAATCACTGCTGACCGGTGAATCTTTGCCTGTCTCCAAAGACGCCGAC	2507
S288C      TTTGATACTGATGAATCACTACTGACCGGTGAATCTTTGCCTGTCTCCAAAGACGCCAAC	2520
AZ118      TTTGATACTGATGAATCACTGCTGACCGGTGAATCTTTGCCTGTCTCCAAAGACGCCGAC	2510
AZ123      TTTGATACTGATGAATCACTGCTGACCGGTGAATCTTTGCCTGTCTCCAAAGACGCCAAC	2508
           ******************** ************************************ **

AZ65       TTGGTATTTGGAAAAGAAGAAGAAACCTCTGTGGGTGATCGTTTGAATTTAGCATTCTCT	2567
S288C      TTAGTGTTTGGAAAAGAAGAAGAAACCTCCGTGGGTGATCGTTTGAATTTAGCATTTTCT	2580
AZ118      TTGGTGTTTGGAAAAGAAGAAGAAACCTCCGTGGGTGATCGTTTGAATTTAGCATTTTCT	2570
AZ123      TTAGTGTTTGGAAAAGAAGAAGAAACCTCCGTGGGTGATCGTTTGAATTTAGCATTTTCT	2568
           ** ** *********************** ************************** ***

AZ65       TCGTCTACCGTTGTCAAAGGAAGGGCGAAGGGTATTGTCATCAAGACAGCTTTGAACAGT	2627
S288C      TCATCCGCCGTTGTCAAGGGAAGAGCCAAGGGTATTGTCATCAAGACAGCTTTGAATAGT	2640
AZ118      TCATCCACCGTTGTCAAGGGAAGAGCCAAGGGTATTGTCATCAAGACAGCTTTGAACAGT	2630
AZ123      TCATCCACCGTTGTCAAGGGAAGAGCCAAGGGTATTGTCATCAAGACAGCTTTGAATAGT	2628
           ** **  ********** ***** ** ***************************** ***

AZ65       GAAATTGGTAAAATTGCAAAATCTCTACAAGGTGATTCAGGTCTCATTTCCCGCGATCCT	2687
S288C      GAAATTGGTAAAATTGCAAAATCTCTACAAGGTGATTCAGGTCTCATTTCCCGCGATCCT	2700
AZ118      GAAATTGGTAAAATTGCAAAATCTCTACAAGGTGATTCAGGTCTCATTTCCCGCGATCCT	2690
AZ123      GAAATTGGTAAAATTGCAAAATCTCTACAAGGTGATTCAGGTCTCATTTCCCGCGATCCT	2688
           ************************************************************

AZ65       AGTAAGTCGTGGTTACAGAATACATGGATATCTACAAAGAAAGTTACTGGTGCATTTTTA	2747
S288C      AGTAAGTCGTGGTTACAGAATACATGGATATCTACAAAGAAAGTTACTGGTGCATTTTTA	2760
AZ118      AGTAAGTCGTGGTTACAGAATACATGGATATCTACAAAGAAAGTTACTGGTGCATTTTTA	2750
AZ123      AGTAAGTCGTGGTTACAGAATACATGGATATCTACAAAGAAAGTTACTGGTGCATTTTTA	2748
           ************************************************************

AZ65       GGTACAAATGTCGGTACGCCCCTGCACAGGAAACTGTCTAAGTTGGCGGTATTGCTTTTC	2807
S288C      GGTACAAATGTCGGTACGCCCCTGCACAGGAAACTGTCTAAGTTGGCGGTATTGCTTTTC	2820
AZ118      GGTACAAATGTCGGTACGCCCCTGCACAGGAAACTGTCTAAGTTGGCGGTATTGCTTTTC	2810
AZ123      GGTACAAATGTCGGTACGCCCCTGCACAGGAAACTGTCTAAGTTGGCGGTATTGCTTTTC	2808
           ************************************************************

AZ65       TGGATAGCTGTATTTGTTTGCTATAATTGTTATGGCCTCTCAAAAGTTCAGTGTAGATAA	2867
S288C      TGGATTGCCGTACTT-TTTGCTATCATCGTCATGGCCTCTCAAAAGTTTGACGTAGATAA	2879
AZ118      TGGATTGCCGTACTT-TTTGCTATCATCGTCATGGCCTCTCAAAAGTTTGACGTAGATAA	2869
AZ123      TGGATTGCCGTACTT-TTTGCTATCATCGTCATGGCCTCTCAAAAGTTTGACGTAGATAA	2867
           ***** ** *** ** ******** ** ** *****************    ********

AZ65       GGGTATAGCTATCTATGCAATTTGCGTTGCCCTATCTATGATTCCCTCTTCATTAGTCGT	2927
S288C      AAGGGTAGCTATCTATGCCATTTGTGTGGCCCTATCCATGATTCCCTCTTCATTAGTCGT	2939
AZ118      AAGGGTAGCTATCTATGCCATTTGTGTGGCCCTATCCATGATTCCCTCTTCATTAGTCGT	2929
AZ123      AAGGGTAGCTATCTATGCCATTTGTGTGGCCCTATCCATGATTCCCTCTTCATTAGTCGT	2927
             *  ************* ***** ** ******** ***********************
AZ65       TGTCTTGACTATTACTATGTCTGTAGGAGCTGCCGTTATGGTTTCTAGACATGTTATTGT	2987
S288C      TGTCTTGACCATCACCATGTCTGTTGGGGCTGCTGTTATGGTTTCTAGAAACGTTATTGT	2999
AZ118      TGTCTTGACCATCACCATGTCTGTTGGGGCTGCTGTTATGGTTTCTAGAAACGTTATTGT	2989
AZ123      TGTCTTGACCATCACCATGTCTGTTGGGGCTGCTGTTATGGTTTCTAGAAACGTTATTGT	2987
           ********* ** ** ******** ** ***** *************** * ********

AZ65       AAGAAAATTAGATTCTTTGGAGGCCCTAGGTGCAGTTAACGACATTTGTTCTGACAAGAC	3047
S288C      AAGAAAATTAGATTCTCTGGAAGCTTTAGGTGCCGTTAACGATATCTGTTCTGACAAGAC	3059
AZ118      AAGAAAATTAGATTCTCTGGAAGCTTTAGGTGCCGTTAACGATATCTGTTCTGACAAGAC	3049
AZ123      AAGAAAATTAGATTCTCTGGAAGCTTTAGGTGCCGTTAACGATATCTGTTCTGACAAGAC	3047
           **************** **** **  ******* ******** ** **************

AZ65       CGGTACTCTTACACAGGGAAAAATGTTAGCAAGGCAGATTTGGATCCCTCGCTTTGGTAC	3107
S288C      CGGTACTCTTACACAGGGTAAAATGTTAGCGAGGCAAATTTGGATCCCTCGCTTTGGTAC	3119
AZ118      CGGTACTCTTACACAGGGTAAAATGTTAGCGAGGCAAATTTGGATCCCTCGCTTTGGTAC	3109
AZ123      CGGTACTCTTACACAGGGTAAAATGTTAGCGAGGCAAATTTGGATCCCTCGCTTTGGTAC	3107
           ****************** *********** ***** ***********************

AZ65       TATAACTATCTCGAATTCTGATGACCCCTTTAATCCCAATGAGGGCAACGTGAGTTTGAT	3167
S288C      CATAACTATCTCGAATTCTGATGACCCCTTTAATCCCAATGAGGGCAACGTGAGTTTGAT	3179
AZ118      CATAACTATCTCGAATTCTGATGACCCCTTTAATCCCAATGAGGGCAACGTGAGTTTGAT	3169
AZ123      CATAACTATCTCGAATTCTGATGACCCCTTTAATCCCAATGAGGGCAACGTGAGTTTGAT	3167
            ***********************************************************

AZ65       TCCAAGGTTTTCACCTTACGAGTACTCCCACAATGAGGATGGTGACGTCGGTATTCTCCA	3227
S288C      TCCAAGGTTTTCACCTTACGAATATTCTCATAATGAGGATGGTGACGTTGGTATTCTCCA	3239
AZ118      TCCAAGGTTTTCACCTTACGAATATTCTCATAATGAGGATGGTGACGTTGGTATTCTCCA	3229
AZ123      TCCAAGGTTTTCACCTTACGAATATTCTCATAATGAGGATGGTGACGTTGGTATTCTCCA	3227
           ********************* ** ** ** ***************** ***********

AZ65       AAACTTTCAAGGATCGCCTATACGAAAAGGATTTACCAGAAGATATTGACATGGATCTAT	3287
S288C      GA-ATTTCAAGGATCGCCTATACGAAAAAGATTTACCAGAAGATATTGACATGGATCTAT	3298
AZ118      GA-ATTTCAAGGATCGCCTATACGAAAAAGATTTACCAGAAGATATTGACATGGATCTAT	3288
AZ123      GA-ATTTCAAGGATCGCCTATACGAAAAAGATTTACCAGAAGATATTGACATGGATCTAT	3286
            *  ************************ *******************************

AZ65       TTCAAAAATGGCTCGAAACCGCCACTTTGGCTAACATTGCTACTGTTTTCAAAGATGACG	3347
S288C      TTCAAAAATGGCTCGAAACCGCCACTTTGGCTAACATTGCTACTGTTTTCAAAGATGACG	3358
AZ118      TTCAAAAATGGCTCGAAACCGCCACTTTGGCTAACATTGCTACTGTTTTCAAAGATGACG	3348
AZ123      TTCAAAAATGGCTCGAAACCGCCACTTTGGCTAACATTGCTACTGTTTTCAAAGATGACG	3346
           ************************************************************

AZ65       CAACTGACTGTTGGAAAGCTCACGGTGATCCAACAGAAATTGCCATTCAGGTATTTGCTA	3407
S288C      CAACTGACTGTTGGAAAGCTCATGGTGACCCAACAGAAATTGCGATTCAAGTGTTTGCTA	3418
AZ118      CAACTGACTGTTGGAAAGCTCATGGTGACCCAACAGAAATTGCGATTCAAGTGTTTGCTA	3408
AZ123      CAACTGACTGTTGGAAAGCTCATGGTGACCCAACAGAAATTGCGATTCAAGTGTTTGCTA	3406
           ********************** ***** ************** ***** ** *******

AZ65       CCAAGATGGATTTACCTCGCAATGCCCTTACCGGTGAGAAATCGAATAATCAAAGTAATG	3467
S288C      CTAAGATGGACTTGCCTCACAATGCCCTTACCGGTGAGAAATCGACTAATCAAAGTAATG	3478
AZ118      CTAAGATGGACTTGCCTCGCAATGCCCTTACCGGTGAGAAATCGACTAATCAAAGTAATG	3468
AZ123      CTAAGATGGACTTGCCTCGCAATGCCCTTACCGGTGAGAAATCGACTAATCAAAGTAATG	3466
           * ******** ** **** ************************** **************

AZ65       AAAGCGATAAATCTTCTCTACTGGGTCACAATGAGAAACCCAGTAATGCCCAATTCCAGC	3527
S288C      AGAATGACCAATCCTCTCTTTCACAACACAATGAGAAGCCTGGCAGTGCACAATTCGAAC	3538
AZ118      AGAATGATCAATCCTCTCTTTCACAACACAATGAGAAGCCTGGCAGTGCACAATTCGAAC	3528
AZ123      AGAATGATCAATCCTCTCTTTCACAACACAATGAGAAGCCTGGCAGTGCACAATTCGAAC	3526
           * *  **  **** *****       *********** **  * * *** ****** * *
AZ65       ATATCGCTGAATTTCCATTTGATTCGACAGTGAAGAGAATGTCATCTGTATACTACAACA	3587
S288C      ATATTGCTGAATTCCCATTCGACTCAACTGTGAAGCGAATGTCATCTGTCTACTACAACA	3598
AZ118      ATATTGCTGAATTCCCATTCGACTCAACTGTGAAGCGAATGTCATCTGTCTACTACAACA	3588
AZ123      ATATTGCTGAATTCCCATTCGACTCAACTGTGAAGCGAATGTCATCTGTCTACTACAACA	3586
           **** ******** ***** ** ** ** ****** ************* **********

AZ65       ATCACAACGAAACATACAACGTTTATGGAAAAGGTGCTTTTGAAAGCATTATCAGCTGTT	3647
S288C      ATCACAACGAAACATATAATATTTATGGCAAGGGTGCTTTCGAAAGCATCATCAGTTGTT	3658
AZ118      ATCACAACGAAACATATAATATTTATGGCAAGGGTGCTTTCGAAAGCATCATCAGTTGTT	3648
AZ123      ATCACAACGAAACATATAATATTTATGGCAAGGGTGCTTTCGAAAGCATCATCAGTTGTT	3646
           **************** **  ******* ** ******** ******** ***** ****

AZ65       GCAGCTCTTGGTATGGCAAGGATGGTACAAAAATCACGCCTTTGACCGATTGTGATGTCG	3707
S288C      GCAGTTCTTGGTATGGTAAGGATGGTGTAAAAATCACACCATTGACCGATTGTGATGTCG	3718
AZ118      GCAGTTCTTGGTATGGTAAGGATGGTGTAAAAATCACACCATTGACCGATTGTGATGTCG	3708
AZ123      GCAGTTCTTGGTATGGTAAGGATGGTGTAAAAATCACACCATTGACCGATTGTGATGTCG	3706
           **** *********** *********  ********* ** *******************

AZ65       AAACTATAAGGAAAAATGTTTACAGTCTATCAAATGAAGGTTTAAGAGTCTTGGGGTTTG	3767
S288C      AAACGATAAGGAAAAATGTTTACAGTCTATCAAATGAGGGTTTAAGAGTCTTGGGTTTTG	3778
AZ118      AAACGATAAGGAAAAATGTTTACAGTCTATCAAATGAGGGTTTAAGAGTCTTGGGTTTTG	3768
AZ123      AAACGATAAGGAAAAATGTTTACAGTCTATCAAATGAGGGTTTAAGAGTCTTGGGTTTTG	3766
           **** ******************************** ***************** ****

AZ65       CTTCCAAATCTTTTACCAAGGATGAACAAAGTGACGAGCAATTGAAGAATAATACTTCGA	3827
S288C      CCTCCAAATCTTTCACTAAAGATCAAGTGAATGACGATCAATTGAAAAACATTACTTCAA	3838
AZ118      CCTCCAAATCTTTCACTAAAGATCAAGTGAATGACGATCAATTGAAAAACATTACTTCAA	3828
AZ123      CCTCCAAATCTTTCACTAAAGATCAAGTGAATGACGATCAATTGAAAAACATTACTTCAA	3826
           * *********** ** ** *** **   * ****** ******** ** * ****** *

AZ65       ACAGGGCTGCCGCAGAAAGCGATTTAGTTTTCCTAGGTTTGATTGGTATTTACGATCCAC	3887
S288C      ACAGGGCCACCGCAGAAAGTGATTTAGTTTTCCTAGGGTTGATTGGTATTTACGATCCAC	3898
AZ118      ACAGGGCCACCGCAGAAAGTGATTTAGTTTTCCTAGGGTTGATTGGTATTTACGATCCAC	3888
AZ123      ACAGGGCCACCGCAGAAAGTGATTTAGTTTTCCTAGGGTTGATTGGTATTTACGATCCAC	3886
           *******  ********** ***************** **********************

AZ65       CTAGAAACGAGACTGCCGGTGCAGTCAAGAAGTTTCATCAGGCCGGTATTAACGTCCATA	3947
S288C      CCAGAAATGAGACTGCCGGTGCAGTCAAGAAGTTTCACCAAGCTGGTATTAACGTTCATA	3958
AZ118      CCAGAAATGAGACTGCCGGTGCAGTCAAGAAGTTTCACCAAGCTGGTATTAACGTTCATA	3948
AZ123      CCAGAAATGAGACTGCCGGTGCAGTCAAGAAGTTTCACCAAGCTGGTATTAACGTTCATA	3946
           * ***** ***************************** ** ** *********** ****

AZ65       TGTTAACTGGGGACTTTGTGGGTACAGCAAAGGCTATTGCTCAAGAAGTTGGTATCTTAC	4007
S288C      TGTTAACTGGGGACTTTGTGGGTACAGCAAAGGCTATCGCTCAGGAGGTTGGCATCTTAC	4018
AZ118      TGTTAACTGGGGACTTTGTGGGTACAGCAAAGGCTATCGCTCAGGAGGTTGGCATCTTAC	4008
AZ123      TGTTAACTGGGGACTTTGTGGGTACAGCAAAGGCTATCGCTCAGGAGGTTGGCATCTTAC	4006
           ************************************* ***** ** ***** *******

AZ65       CCACCAATTTGTACCATTACTCCCAAGAGATTGTTGACAGTATGGTCATGACCGGATCTC	4067
S288C      CCACCAATTTGTACCATTACTCCCAAGAGATTGTTGACAGTATGGTCATGACCGGATCTC	4078
AZ118      CCACCAATTTGTACCATTACTCCCAAGAGATTGTTGACAGTATGGTCATGACCGGATCTC	4068
AZ123      CCACCAATTTGTACCATTACTCCCAAGAGATTGTTGACAGTATGGTCATGACCGGATCTC	4066
           ************************************************************

AZ65       AGTTTGACGGACTAAGTGAGGAGGAAGTGGACGATTTGCCCGTCTTACCTTTAGTTATTG	4127
S288C      AGTTTGACGGACTAAGTGAGGAGGAAGTGGACGATTTGCCCGTCTTACCTTTAGTTATTG	4138
AZ118      AGTTTGACGGACTAAGTGAGGAGGAAGTGGACGATTTGCCCGTCTTACCTTTAGTTATTG	4128
AZ123      AGTTTGACGGACTAAGTGAGGAGGAAGTGGACGATTTGCCCGTCTTACCTTTAGTTATTG	4126
           ************************************************************
AZ65       CCCGTTGCTCTCCGCAGACTAAGGTCAGAATGATCGAAGCTTTACACCGTAGGAAGAAGT	4187
S288C      CACGTTGCTCTCCGCAGACTAAGGTGAGAATGATCGAAGCTTTACACCGTAGGAAGAAGT	4198
AZ118      CACGTTGCTCTCCGCAGACTAAGGTGAGAATGATCGAAGCTTTACACCGTAGGAAGAAGT	4188
AZ123      CACGTTGCTCTCCGCAGACTAAGGTGAGAATGATCGAAGCTTTACACCGTAGGAAGAAGT	4186
           * *********************** **********************************

AZ65       TCTGCGCAATGACAGGTGACGGTGTCAACGATTCTCCATCTCTAAAAATGGCCAATGTTG	4247
S288C      TCTGCACAATGACAGGTGACGGTGTTAACGATTCTCCATCTCTAAAAATGGCCAATGTTG	4258
AZ118      TCTGCGCAATGACAGGTGACGGTGTTAATGATTCTCCATCTCTAAAAATGGCCAATGTTG	4248
AZ123      TCTGCGCAATGACAGGTGACGGTGTTAATGATTCTCCATCTCTAAAAATGGCCAATGTTG	4246
           ***** ******************* ** *******************************

AZ65       GTATTGCAATGGGTATTAATGGTTCAGATGTTTCCAAAGAAGCGTCTGATATTGTTCTAA	4307
S288C      GTATTGCAATGGGTATTAATGGTTCAGATGTTTCCAAAGAAGCGTCTGATATTGTTCTAA	4318
AZ118      GTATTGCAATGGGTATTAATGGTTCAGATGTTTCCAAAGAAGCGTCTGATATTGTTCTAA	4308
AZ123      GTATTGCAATGGGTATTAATGGTTCAGATGTTTCCAAAGAAGCGTCTGATATTGTTCTAA	4306
           ************************************************************

AZ65       GCGATGACAATTTTGCTTCTATTTTGAATGCTGTCGAAGAAGGTCGTAGGATGACGGATA	4367
S288C      GCGATGACAATTTTGCTTCTATTTTGAATGCTGTCGAAGAAGGTCGTAGGATGACGGATA	4378
AZ118      GCGATGACAATTTTGCTTCTATTTTGAATGCTGTCGAAGAAGGTCGTAGGATGACGGATA	4368
AZ123      GCGATGACAATTTTGCTTCTATTTTGAATGCTGTCGAAGAAGGTCGTAGGATGACGGATA	4366
           ************************************************************

AZ65       ACATTCAGAAGTTTGTCCTACAATTATTGGCAGAAAATGTTGCTCAGGCTTTGTATTTGA	4427
S288C      ACATTCAGAAGTTTGTCCTACAATTATTGGCAGAAAATGTTGCTCAGGCTTTGTATTTGA	4438
AZ118      ACATTCAGAAGTTTGTCCTACAATTATTGGCAGAAAATGTTGCTCAGGCTTTGTATTTGA	4428
AZ123      ACATTCAGAAGTTTGTCCTACAATTATTGGCAGAAAATGTTGCTCAGGCTTTGTATTTGA	4426
           ************************************************************

AZ65       TCATTGGTTTAGTATTCAGAGATGAGAACGGAAAATCAGTGTTTCCCTTATCGCCTGTAG	4487
S288C      TCATTGGTTTAGTATTCAGAGATGAGAACGGAAAATCAGTATTTCCCTTATCACCAGTGG	4498
AZ118      TCATTGGTTTAGTATTCAGAGATGAGAACGGAAAATCAGTATTTCCCTTATCACCAGTGG	4488
AZ123      TCATTGGTTTAGTATTCAGAGATGAGAACGGAAAATCAGTATTTCCCTTATCACCAGTGG	4486
           **************************************** *********** ** ** *

AZ65       AAGTTTTATGGATTATCGTTGTCACCTCTTGTTTCCCTGCTATGGGGCTAGGTCTAGAAA	4547
S288C      AAGTATTGTGGATTATTGTCGTCACCTCTTGTTTTCCTGCTATGGGGCTAGGTCTAGAAA	4558
AZ118      AAGTATTGTGGATTATTGTCGTCACCTCTTGTTTTCCTGCTATGGGGCTAGGTCTAGAAA	4548
AZ123      AAGTATTGTGGATTATTGTCGTCACCTCTTGTTTTCCTGCTATGGGGCTAGGTCTAGAAA	4546
           **** ** ******** ** ************** *************************

AZ65       AGGCTGCTCCAGATTTAATGGATAGACCTCCTAACGATTCAGAAGTTGGTATTTTCACCT	4607
S288C      AGGCTGCTCCAGATTTGATGGATAGACCTCCTCATGATTCAGAGGTTGGTATTTTCACGT	4618
AZ118      AGGCTGCTCCAGATTTGATGGATAGACCTCCTAATGATTCAGAGGTTGGTATTTTCACAT	4608
AZ123      AGGCTGCTCCAGATTTGATGGATAGACCTCCTAATGATTCAGAGGTTGGTATTTTCACAT	4606
           **************** *************** * ******** ************** *

AZ65       GGGAAGTTATTATCGATACATTTGCATACGGGTTTTTGATGGCCGGTTCTTGTATGGCCT	4667
S288C      GGGAGGTTATTATAGATACATTTGCATATGGGATTATAATGACAGGGTCCTGTATGGCTT	4678
AZ118      GGGAGGTTATTATAGATACATTTGCATATGGGATTATAATGACAGGGTCCTGTATGGCTT	4668
AZ123      GGGAGGTTATTATAGATACATTTGCATATGGGATTATAATGACAGGGTCCTGTATGGCTT	4666
           **** ******** ************** *** ** * *** * ** ** ******** *

AZ65       CCTTCACTGGGTCGCTCTACGGAATAAATAATGGTGAATTGGGAGTTAATTGTGATAGAT	4727
S288C      CATTTACTGGATCACTGTATGGAATAAATAGTGGTAGATTGGGGCACGATTGTGATGGCA	4738
AZ118      CATTTACTGGATCACTGTATGGAATAAATAGTGGTAGATTGGGGCACGATTGTGATGGCA	4728
AZ123      CATTTACTGGATCACTGTATGGAATAAATAGTGGTAGATTGGGGCACGATTGTGATGGCA	4726
           * ** ***** ** ** ** ********** ****  ******     ******** *
AZ65       CTTATAATGGTAGCTGTCATGATGTTTATAGATCACGTTCTGCAGCCTTCGCAACTATGA	4787
S288C      CCTATAACAGCAGTTGTCGTGATGTTTATAGATCACGTTCTGCGGCTTTCGCAACCATGA	4798
AZ118      CCTATAACAGCAGTTGTCGTGATGTTTATAGATCACGTTCTGCGGCTTTCGCAACTATGA	4788
AZ123      CCTATAACAGCAGTTGTCGTGATGTTTATAGATCACGTTCTGCGGCTTTCGCAACTATGA	4786
           * *****  * ** **** ************************ ** ******** ****

AZ65       CATGGTGTGCTTTGATTTTGGCTTGGGAAGTGGTTGATATGAGAAGATCTTTTTTCAGAA	4847
S288C      CGTGGTGCGCTTTGATTCTGGCTTGGGAAGTGGTTGACATGAGAAGATCCTTTTTCAGAA	4858
AZ118      CGTGGTGCGCTTTGATTCTGGCTTGGGAAGTGGTTGACATGAGAAGATCTTTTTTCAGAA	4848
AZ123      CGTGGTGCGCTTTGATTCTGGCTTGGGAAGTGGTTGACATGAGAAGATCTTTTTTCAGAA	4846
           * ***** ********* ******************* *********** **********

AZ65       TGCATCCAGACACTGACAGCCCAGTTAAGGAATTTTTTAAGAGTATTTGGGGAAACCAGT	4907
S288C      TGCATCCAGACACTGACAGCCCAGTCAAGGAATTTTTCAGAAGCATTTGGGGAAACCAGT	4918
AZ118      TGCATCCAGACACTGACAGCCCAGTCAAGGAATTTTTCAGAAGTATTTGGGGAAACCAGT	4908
AZ123      TGCATCCAGACACTGACAGCCCAGTCAAGGAATTTTTCAGAAGTATTTGGGGAAACCAGT	4906
           ************************* *********** *  ** ****************

AZ65       TTTTGTTTTGGTCAATCATTTTTGGATTTGTGTCTGCCTTCCCTGTCGTCTATATTCCGG	4967
S288C      TTTTGTTCTGGTCAATCATTTTTGGATTTGTGTCAGCCTTCCCCGTCGTCTATATTCCGG	4978
AZ118      TTTTGTTCTGGTCAATCATTTTTGGATTTGTGTCAGCCTTCCCCGTCGTCTATATTCCGG	4968
AZ123      TTTTGTTCTGGTCAATCATTTTTGGATTTGTGTCAGCCTTCCCCGTCGTCTATATTCCGG	4966
           ******* ************************** ******** ****************

AZ65       TTATTAATGATAAAGTGTTTTTGCATAAACCAATTGGTCCTGAATGGGGTCTCGCCATTG	5027
S288C      TTATTAATGATAAAGTGTTTTTGCATAAACCAATTGGTGCTGAATGGGGTCTCGCCATTG	5038
AZ118      TTATTAATGATAAAGTGTTTTTGCATAAACCAATTGGTGCTGAATGGGGTCTCGCCATTG	5028
AZ123      TTATTAATGATAAAGTGTTTTTGCATAAACCAATTGGTGCTGAATGGGGTCTCGCCATTG	5026
           ************************************** *********************

AZ65       CATTCACAATTGCATTCTGGATTGGTGCTGAACTTTACAAGTTAGGGAAGAGGCGCTACT	5087
S288C      CATTCACAATTGCATTCTGGATAGGTGCTGAACTTTACAAGTGTGGAAAGAGGCGCTATT	5098
AZ118      CATTCACAATTGCATTCTGGATAGGTGCTGAACTTTACAAGTGTGGAAAGAGGCGCTATT	5088
AZ123      CATTCACAATTGCATTCTGGATAGGTGCTGAACTTTACAAGTGTGGAAAGAGGCGCTATT	5086
           ********************** *******************  ** *********** *

AZ65       TCAAAACTCAGAGAGCACACAACCCGGAAAACGATTTGGAGAGTAACAATAAGCGCGATC	5147
S288C      TCAAAACTCAGAGAGCGCACAACCCGGAGAATGATTTGGAGAGTAACAATAAGCGCGATC	5158
AZ118      TCAAAACTCAGAGAGCGCACAACCCGGAGAATGATTTGGAGAGTAACAATAAGCGCGATC	5148
AZ123      TCAAAACTCAGAGAGCGCACAACCCGGAGAATGATTTGGAGAGTAACAATAAGCGCGATC	5146
           **************** *********** ** ****************************

AZ65       CATTCGAAGCGTACAGTACTTCTACTACAATCCATACAGAAATTAATATTGGTATCAAAC	5207
S288C      CATTCGAAGCGTATAGTACTTCTACTACAATCCATACAGAAGTTAATATTGGTATTAAAC	5218
AZ118      CATTCGAAGCGTATAGTACTTCTACTACAATCCATACAGAAGCTAATATTGGTATTAAAC	5208
AZ123      CATTCGAAGCGTATAGTACTTCTACTACAATCCATACAGAAGCTAATATTGGTATTAAAC	5206
           ************* ***************************  ************ ****

AZ65       AATGA	5212
S288C      AATGA	5223
AZ118      AATGA	5213
AZ123      AATGA	5211
           *****
```
