## Supplemental Figure 4 for "Analysis of genome sequences of marine isolates of *Saccharomyces cerevisiae*"

**Supplementary Figure S4**: Alignment of internal promoter regions (intergenic regions between two *ENA* copies). No sequence variations were detected. The stop codon of the preceding *ENA* copy and start codon of the subsequent *ENA* copy are highlighted in green.

```
AZ123A2      TAGGCAGGGCCTATTAAGTGCTCCCTATCTTTTTCCTTATTCATTTCATATATTTAGATA	60
AZ65A4       TAGGCAGGGCCTATTAAGTGCTCCCTATCTTTTTCCTTATTCATTTCATATATTTAGATA	60
ENA5         TAGGCAGGGCCTATTAAGTGCTCCCTATCTTTTTCCTTATTCATTTCATATATTTAGATA	60
AZ118A2      TAGGCAGGGCCTATTAAGTGCTCCCTATCTTTTTCCTTATTCATTTCATATATTTAGATA	60
AZ65A2       TAGGCAGGGCCTATTAAGTGCTCCCTATCTTTTTCCTTATTCATTTCATATATTTAGATA	60
AZ65A3       TAGGCAGGGCCTATTAAGTGCTCCCTATCTTTTTCCTTATTCATTTCATATATTTAGATA	60
ENA2         TAGGCAGGGCCTATTAAGTGCTCCCTATCTTTTTCCTTATTCATTTCATATATTTAGATA	60
             ************************************************************

AZ123A2      TTTACATTTATTTTGTTATTTGTTTTTTTATTTACTATAAACGAACTTTTCACAGGTTCT	120
AZ65A4       TTTACATTTATTTTGTTATTTGTTTTTTTATTTACTATAAACGAACTTTTCACAGGTTCT	120
ENA5         TTTACATTTATTTTGTTATTTGTTTTTTTATTTACTATAAACGAACTTTTCACAGGTTCT	120
AZ118A2      TTTACATTTATTTTGTTATTTGTTTTTTTATTTACTATAAACGAACTTTTCACAGGTTCT	120
AZ65A2       TTTACATTTATTTTGTTATTTGTTTTTTTATTTACTATAAACGAACTTTTCACAGGTTCT	120
AZ65A3       TTTACATTTATTTTGTTATTTGTTTTTTTATTTACTATAAACGAACTTTTCACAGGTTCT	120
ENA2         TTTACATTTATTTTGTTATTTGTTTTTTTATTTACTATAAACGAACTTTTCACAGGTTCT	120
             ************************************************************

AZ123A2      GAACATAAACAAGCCTAAATAGTATGTTTTCTAGTTCAAATATGTCTTAGAGGAATACTA	180
AZ65A4       GAACATAAACAAGCCTAAATAGTATGTTTTCTAGTTCAAATATGTCTTAGAGGAATACTA	180
ENA5         GAACATAAACAAGCCTAAATAGTATGTTTTCTAGTTCAAATATGTCTTAGAGGAATACTA	180
AZ118A2      GAACATAAACAAGCCTAAATAGTATGTTTTCTAGTTCAAATATGTCTTAGAGGAATACTA	180
AZ65A2       GAACATAAACAAGCCTAAATAGTATGTTTTCTAGTTCAAATATGTCTTAGAGGAATACTA	180
AZ65A3       GAACATAAACAAGCCTAAATAGTATGTTTTCTAGTTCAAATATGTCTTAGAGGAATACTA	180
ENA2         GAACATAAACAAGCCTAAATAGTATGTTTTCTAGTTCAAATATGTCTTAGAGGAATACTA	180
             ************************************************************

AZ123A2      CTGCTATACTCTTATTGCTGCCGTTGTAAATCATGAAACAGATACAGTTACGTTACGCGT	240
AZ65A4       CTGCTATACTCTTATTGCTGCCGTTGTAAATCATGAAACAGATACAGTTACGTTACGCGT	240
ENA5         CTGCTATACTCTTATTGCTGCCGTTGTAAATCATGAAACAGATACAGTTACGTTACGCGT	240
AZ118A2      CTGCTATACTCTTATTGCTGCCGTTGTAAATCATGAAACAGATACAGTTACGTTACGCGT	240
AZ65A2       CTGCTATACTCTTATTGCTGCCGTTGTAAATCATGAAACAGATACAGTTACGTTACGCGT	240
AZ65A3       CTGCTATACTCTTATTGCTGCCGTTGTAAATCATGAAACAGATACAGTTACGTTACGCGT	240
ENA2         CTGCTATACTCTTATTGCTGCCGTTGTAAATCATGAAACAGATACAGTTACGTTACGCGT	240
             ************************************************************

AZ123A2      ATGACAGGGTTCTTAACTAAAATTATGAATTTCAATACAGGATTAGTTAAATTCTTGTAG	300
AZ65A4       ATGACAGGGTTCTTAACTAAAATTATGAATTTCAATACAGGATTAGTTAAATTCTTGTAG	300
ENA5         ATGACAGGGTTCTTAACTAAAATTATGAATTTCAATACAGGATTAGTTAAATTCTTGTAG	300
AZ118A2      ATGACAGGGTTCTTAACTAAAATTATGAATTTCAATACAGGATTAGTTAAATTCTTGTAG	300
AZ65A2       ATGACAGGGTTCTTAACTAAAATTATGAATTTCAATACAGGATTAGTTAAATTCTTGTAG	300
AZ65A3       ATGACAGGGTTCTTAACTAAAATTATGAATTTCAATACAGGATTAGTTAAATTCTTGTAG	300
ENA2         ATGACAGGGTTCTTAACTAAAATTATGAATTTCAATACAGGATTAGTTAAATTCTTGTAG	300
             ************************************************************

AZ123A2      AATTAGAAAATTAATGCGTTTATTTCGTCAAGACTTTTCCTTCCAGGACGTTCAAATTCA	360
AZ65A4       AATTAGAAAATTAATGCGTTTATTTCGTCAAGACTTTTCCTTCCAGGACGTTCAAATTCA	360
ENA5         AATTAGAAAATTAATGCGTTTATTTCGTCAAGACTTTTCCTTCCAGGACGTTCAAATTCA	360
AZ118A2      AATTAGAAAATTAATGCGTTTATTTCGTCAAGACTTTTCCTTCCAGGACGTTCAAATTCA	360
AZ65A2       AATTAGAAAATTAATGCGTTTATTTCGTCAAGACTTTTCCTTCCAGGACGTTCAAATTCA	360
AZ65A3       AATTAGAAAATTAATGCGTTTATTTCGTCAAGACTTTTCCTTCCAGGACGTTCAAATTCA	360
ENA2         AATTAGAAAATTAATGCGTTTATTTCGTCAAGACTTTTCCTTCCAGGACGTTCAAATTCA	360
             ************************************************************
```

AZ123A2	ACTCACATATATTCTTTTATCAAAATCCTTATATCACCGCGCGGTGTAATTGTGAGGCTG	420

```
AZ65A4       ACTCACATATATTCTTTTATCAAAATCCTTATATCACCGCGCGGTGTAATTGTGAGGCTG	420
ENA5         ACTCACATATATTCTTTTATCAAAATCCTTATATCACCGCGCGGTGTAATTGTGAGGCTG	420
AZ118A2      ACTCACATATATTCTTTTATCAAAATCCTTATATCACCGCGCGGTGTAATTGTGAGGCTG	420
AZ65A2       ACTCACATATATTCTTTTATCAAAATCCTTATATCACCGCGCGGTGTAATTGTGAGGCTG	420
AZ65A3       ACTCACATATATTCTTTTATCAAAATCCTTATATCACCGCGCGGTGTAATTGTGAGGCTG	420
ENA2         ACTCACATATATTCTTTTATCAAAATCCTTATATCACCGCGCGGTGTAATTGTGAGGCTG	420
             ************************************************************

AZ123A2      ATGTTTTCTTCTTTAAATCTCCATATTATCTCTCGAGGCAAAGAAAGAGGAAAGAATTTT	480
AZ65A4       ATGTTTTCTTCTTTAAATCTCCATATTATCTCTCGAGGCAAAGAAAGAGGAAAGAATTTT	480
ENA5         ATGTTTTCTTCTTTAAATCTCCATATTATCTCTCGAGGCAAAGAAAGAGGAAAGAATTTT	480
AZ118A2      ATGTTTTCTTCTTTAAATCTCCATATTATCTCTCGAGGCAAAGAAAGAGGAAAGAATTTT	480
AZ65A2       ATGTTTTCTTCTTTAAATCTCCATATTATCTCTCGAGGCAAAGAAAGAGGAAAGAATTTT	480
AZ65A3       ATGTTTTCTTCTTTAAATCTCCATATTATCTCTCGAGGCAAAGAAAGAGGAAAGAATTTT	480
ENA2         ATGTTTTCTTCTTTAAATCTCCATATTATCTCTCGAGGCAAAGAAAGAGGAAAGAATTTT	480
             ************************************************************

AZ123A2      ATTTTCCTTCATCCTTTACATCGAGAATACGTTAACCAAATCAACGTTATTCATATTCGT	540
AZ65A4       ATTTTCCTTCATCCTTTACATCGAGAATACGTTAACCAAATCAACGTTATTCATATTCGT	540
ENA5         ATTTTCCTTCATCCTTTACATCGAGAATACGTTAACCAAATCAACGTTATTCATATTCGT	540
AZ118A2      ATTTTCCTTCATCCTTTACATCGAGAATACGTTAACCAAATCAACGTTATTCATATTCGT	540
AZ65A2       ATTTTCCTTCATCCTTTACATCGAGAATACGTTAACCAAATCAACGTTATTCATATTCGT	540
AZ65A3       ATTTTCCTTCATCCTTTACATCGAGAATACGTTAACCAAATCAACGTTATTCATATTCGT	540
ENA2         ATTTTCCTTCATCCTTTACATCGAGAATACGTTAACCAAATCAACGTTATTCATATTCGT	540
             ************************************************************

AZ123A2      ATTTTGCTTATCAGATAACGTATGTACTCACTGAGGAAAAATCTTTGTACACATAATTGA	600
AZ65A4       ATTTTGCTTATCAGATAACGTATGTACTCACTGAGGAAAAATCTTTGTACACATAATTGA	600
ENA5         ATTTTGCTTATCAGATAACGTATGTACTCACTGAGGAAAAATCTTTGTACACATAATTGA	600
AZ118A2      ATTTTGCTTATCAGATAACGTATGTACTCACTGAGGAAAAATCTTTGTACACATAATTGA	600
AZ65A2       ATTTTGCTTATCAGATAACGTATGTACTCACTGAGGAAAAATCTTTGTACACATAATTGA	600
AZ65A3       ATTTTGCTTATCAGATAACGTATGTACTCACTGAGGAAAAATCTTTGTACACATAATTGA	600
ENA2         ATTTTGCTTATCAGATAACGTATGTACTCACTGAGGAAAAATCTTTGTACACATAATTGA	600
             ************************************************************

AZ123A2      AAATTTTCGATGAGCGAGGGAACTGTCAAAGAAAACAATAATGAAGAATTCAATGCTTAT	660
AZ65A4       AAATTTTCGATGAGCGAGGGAACTGTCAAAGAAAACAATAATGAAGAATTCAATGCTTAT	660
ENA5         AAATTTTCGATGAGCGAGGGAACTGTCAAAGAAAACAATAATGAAGAATTCAATGCTTAT	660
AZ118A2      AAATTTTCGATGAGCGAGGGAACTGTCAAAGAAAACAATAATGAAGAATTCAATGCTTAT	660
AZ65A2       AAATTTTCGATGAGCGAGGGAACTGTCAAAGAAAACAATAATGAAGAATTCAATGCTTAT	660
AZ65A3       AAATTTTCGATGAGCGAGGGAACTGTCAAAGAAAACAATAATGAAGAATTCAATGCTTAT	660
ENA2         AAATTTTCGATGAGCGAGGGAACTGTCAAAGAAAACAATAATGAAGAATTCAATGCTTAT	660
             ************************************************************
```
